# Analysis of protein aging reveals rates of subcellular organelle renewal and selective post-translational modification in Arabidopsis

**DOI:** 10.1101/2025.06.09.658759

**Authors:** Nathan D. Tivendale, Xiaoyu Liu, A. Harvey Millar

**Affiliations:** School of Molecular Sciences, The University of Western Australia, 35 Stirling Hwy, Crawley, WA, 6009, Australia; Tasmanian Institute of Agriculture, University of Tasmania, Newnham Dr, Newnham, TAS, 7248 Australia

**Keywords:** proteomics, BONCAT, Arabidopsis, posttranslational modification, protein turnover

## Abstract

The cellular proteome represents a mixture of older and newer copies of each protein type and turnover of this mixture occurs by cycles of protein synthesis and degradation. There is considerable research on new protein synthesis, the nature of nascent proteins and cellular machinery of protein degradation. However, we have limited insights into older proteins or the protein aging process in plants at scale. In this study we use pulse chase biorthogonal non-canonical amino acid tagging (BONCAT) in Arabidopsis cells coupled to affinity purification to capture and analyse snapshots of the cellular proteome as it ages over a two-week period. Each snapshot was subjected to peptide mass spectrometry-based identification, quantitation and characterisation. We show that there are a broad range of lifespans among the 1688 proteins studied and that their subcellular location correlates strongly with protein longevity. Mitochondria, plastids and the extracellular environment contained the longest lived sub-proteomes while the vesicular pathway to ER, PM and peroxisomes contained the shortest-lived protein sets. Abundant primary metabolic enzymes have considerable longevity, while kinases and ubiquitination machinery do not. Through analysis of the aging profiles, we demonstrate that many proteins selectively accumulate posttranslational modifications (PTMs) as they age and that these are mostly oxidative in nature. We show by analysis of exemplar proteins that distinct PTM profiles and proportional changes with age exist between proteins, likely dictated by differences in subcellular environment and protein function. Implications of these insights for understanding cellular function and for biotechnological modification of the plant proteome are discussed.

## Introduction

Understanding the nature of proteins as they age *in vivo* in plants has broad implications for plant biology, biotechnology, agriculture, and environmental science. In all cellular organisms, protein pools are constantly being renewed through a cyclical process of synthesis, degradation and re-synthesis (Basisty et al., 2020; Tivendale et al., 2021; Tivendale et al., 2020). Turnover rates in plants vary greatly between proteins, with some proteins displaying half-lives of several hours and others showing half-lives up to several months (Li et al., 2017). This diversity of protein turnover rates means that there is a diversity of protein ages in each protein pool. Developmental processes in plants are intrinsically linked to protein aging. For example, rapid proliferation of cells in meristems leads to forever young proteomes, while in leaf senescence and seed maturation protein aging and functional loss become important rhythms of the process (Nelson and Millar, 2015). During abiotic stress, plant growth rate slows, aging the proteome and proteins are also vulnerable to acceleration of the aging process due to drought, heat and cold that can accelerate protein damage through oxidation, glycation, and misfolding (Sweetlove and Møller, 2009). Photosynthetic and respiratory efficiency requires optimal protein function in environments with light and redox induced reactive oxygen chemistry that damages protein machinery as well as phospholipids and DNA (Møller and Kristensen, 2004; Watson et al., 2018). These processes dictate complex regulation of protein synthesis and renewal of specific components for plastids and mitochondria. Protein degradation occurs through different systems in the cytosol, endosymbiotic organelles and the vacuole (Vierstra, 1996). But it is still unclear how these systems sift and detect exactly which protein molecules from a pool are targeted through autophagic or ubiquitin proteosome conjugation systems for removal from the cell (Isono et al., 2024). Crop quality and end-use relates to the harvest and post-harvest nature of proteins and their properties but limited markers of protein functional loss are currently known or measured. Biotechnology and synthetic biology aim to alter the complement of proteins in cells and maintain recombinant proteins plus their abundances or locations in cells. But these are in balance with endogenous protein degradation systems that often blunt engineering in favour of homeostasis.

One of the chief reasons that when studying a native protein of interest or designing a synthetic biology device, the lifespan of the proteins involved and the structural changes that occur over a protein’s lifetime are not considered, is technological. The diversity of protein turnover rates (Li *et al*., 2017; Tivendale *et al*., 2021) has critical implications for the efficiency and effectiveness of enzyme systems, since protein turnover can account for a large portion of the cellular maintenance energy budget (De Vries, 1975). While it is established that protein molecules go through numerous changes as they age, including post-translational modifications (PTMs) (Schöneich, 2006), quantification of these changes at scale has not been reported. We know from specific examples that age-related PTMs affect protein structure, turnover and localization (Schöneich, 2006) and may diminish the catalytic rate of enzymes (Gafni and Noy, 1984; Machado et al., 1991) or increase protein promiscuity (Nobeli et al., 2009), and this may cause decreased efficiency, but it is hard to study this beyond specific examples.

All of this raises three questions: what does a stable protein look like? How do the structures of protein molecules change over their lifetime? And are there ways to predict this for a given protein of interest? In much the same way as archaeology provides an understanding of what people did, where they lived, what sorts of tools they used, what sort of lives they led and how the environment has changed, examining the old proteome—protein archaeology—could teach us about the lives of protein molecules and the environment they have lived through. If we are able to extract and study the old plant proteome, we can learn which proteins remain in cells for long periods of time, how they might be altered in abundance or PTMs and how this might impact protein function.

Past studies have given insight into the degradation rates of large protein sets in plants (Li *et al*., 2017), but these studies are based on calculations from mass resolved spectral data and do not allow us to biochemically separate and independently examine old and new protein molecules. We previously demonstrated that homopropargylglycine (HPG) was a suitable non-canonical amino acid for performing biorthogonal non-canonical amino acid tagging (BONCAT) in plants, enabling separation of nascent proteins from the rest of the proteome (Tivendale *et al*., 2021). We demonstrated that HPG is superior to azidohomoalanine (AHA, a non-canonical amino acid commonly used for BONCAT in mammalian systems) for BONCAT in plants and showed how the nascent protein pool differs from the bulk proteome. In this study, we use this tool, typically employed to look at nascent proteins through pulse experiments, to examine the longevity of *Arabidopsis thaliana* proteins after they are made and study time-resolved data on PTM changes that occur as proteins age. Based on our analysis, we uncover features of the natural dynamics of protein aging within the substructure of Arabidopsis cells and provide recommendations for incorporating age-related protein data analysis into future synthetic biology device design.

## Results

### Protein longevity quotient reveals a broad range of protein lifespans

In this study, we performed a pulse-chase experiment using HPG to tag nascent proteins with an alkyne handle over a 24-h period in Arabidopsis cell culture. Specifically, five-day-old cell cultures were sampled, and the remainder of each culture was labelled with the non-canonical amino acid HPG (pulse) and sampled after 24 hours, after which cells were washed free of HPG-containing media and, grown on HPG-free media for two weeks (chase). Cultures were sampled at 1, 2, 4, 7, 9, 12 and 14 days during the chase. By collecting samples at these nine timepoints over the two weeks, we were then able to enrich the tagged proteins from each sample using azide-alkyne cycloaddition agarose enrichment media to collect protein pools of various ages. Once HPG-containing proteins were covalently attached to the resin beads they could be washed with high stringency removing non-specifically bound proteins. Trypsin digestion of the bead-bound proteins yielded a highly enriched population of peptides from HPG-containing proteins. Analysis of the peptides found in these different sample pools by mass spectrometry enabled us to see the range of protein lifespans and determine how protein abundance varies and PTM profiles change over time for different protein groups (Supplemental Table S1 and S2).

After analysis, the proteins identified and quantified at each time point were found to be consistent, with approximately 1945–2400 unique proteins (AGIs) being shared between all three replicates at a timepoint. The only notable exceptions were for the 4-day sample and 7-day samples, which showed 1690 and 1643 AGIs, respectively, shared between all replicates. At the 4-day sampling, an additional 955 AGIs were shared between only replicates 1 and 3 and at the 7-day sampling, an additional 933 AGIs were shared between only replicates 2 and 3. These results indicate that, while the sampling and analysis was consistent overall, there was some stochasticity in the enriched protein datasets (Supplemental Figure S1).

After quality filtering, we settled on a set of 1688 unique proteins found across the eight timepoints. Enriched protein fractions isolated at different timepoints were subject to differential abundant protein (DAP) analysis. The filtering included removing 242 proteins that appeared to change in abundance in response to the HPG pulse itself. We extracted total proteins immediately post-pulse and 7 and 14 days after the pulse and compared them, without enrichment, to the immediately post-pulse proteome. Any proteins with a log_2_(fold-change) ≥ 1.6 or ≤ –1.6 at the 14-day timepoint compared to the pre-pulse proteome was omitted from further analysis (Supplemental Figure S2). The group of 242 proteins was analysed using Fisher’s exact test to determine if any subcellular locations or MapMan functional categories were overrepresented in the group. Extracellular and peroxisomal proteins were found to be significantly (p < 0.05) overrepresented. Proteins in the ribosomal subunits functional category were significantly (p < 0.05) underrepresented in the group.

By comparing the relative intensity at a given timepoint for a given protein in the enriched sample the relative intensity for the same protein immediately post-pulse, we determined which proteins are longer or shorter lived than the average. This value, which we call the longevity quotient, is defined for each protein in a sample as:

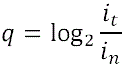

where *q* is ‘the longevity quotient’, *i_n_* is ‘the relative intensity immediately post-pulse’ and *i_t_* is ‘the relative intensity at time *t*’. To assess the longevity of different proteins, we defined short-lived as having *q* ≤ –1.6, median-lived as have –1.6 < *q* < 1.6 (a *q* value of 0 implies no change in relative abundance between timepoints) and long-lived as having *q* ≥ 1.6. Analysis of longevity quotients, revealed a wide spread of mean lifespans, ranging from –4.57 to 3.43 (Figure A). As expected, when assessing longevity quotients over time, the spread away from the centre became more apparent (Figure 1B).

**Figure 1.**
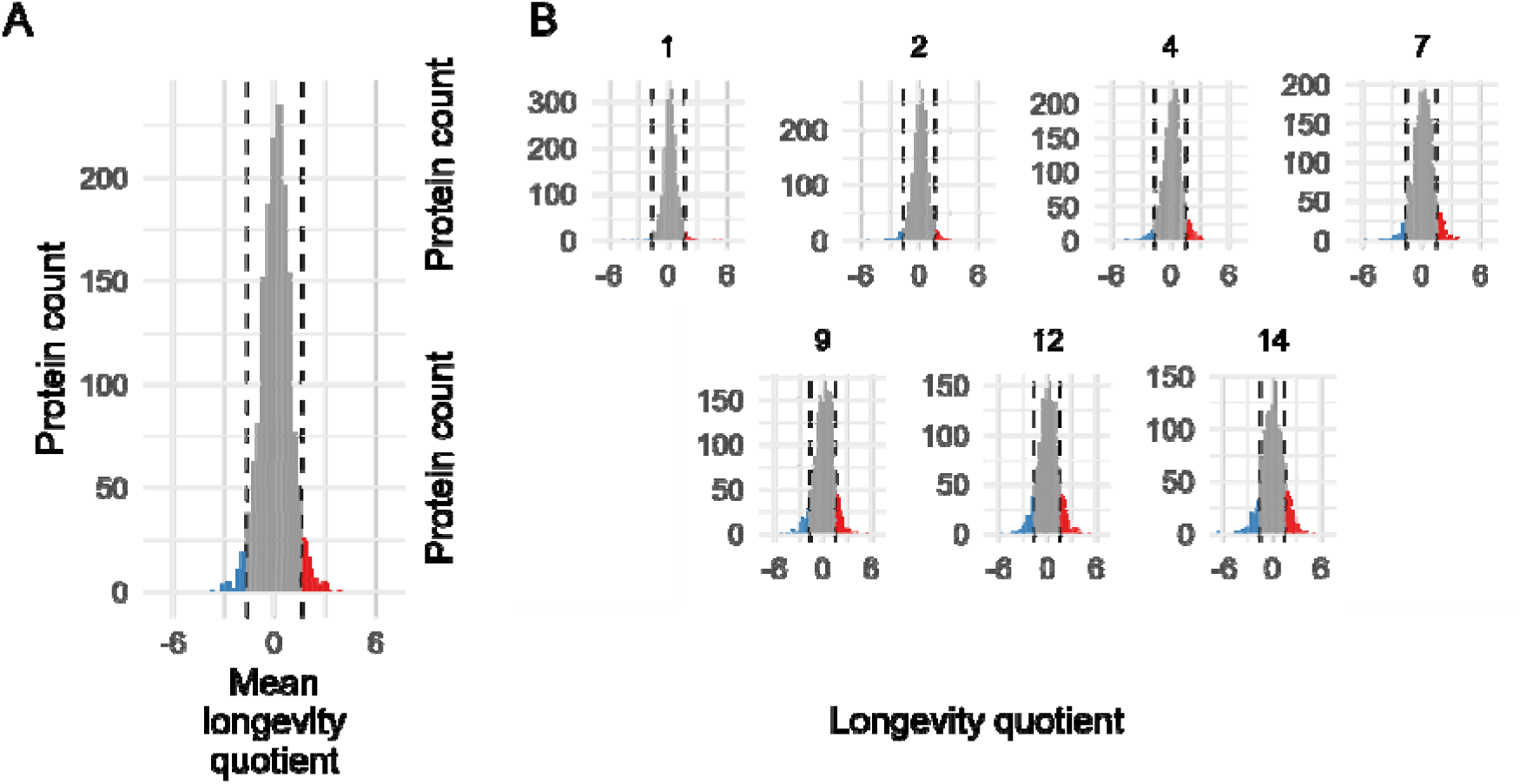
(A) Mean longevity quotient (*q*) across all timepoints. Dashed vertical lines are at –1.6 and 1.6 and indicate cutoff between short-lived (blue, *q* ≤ –1.6), average longevity (–1.6 < *q* < 1.6) and long-lived (red, *q* ≥ 1.6) proteins. (B) Longevity quotient for all proteins faceted by time (days). Dashed vertical lines as for A.

### Mitochondrial, plastid and extracellular proteins tend to be longer-lived

To understand the wide range of longevities apparent in the data, we grouped the proteins by their subcellular compartment, based on SUBAcon classification (Hooper et al., 2014). Looking at the analysis, it became clear many of the abundant proteins that increased in abundance over the course of the experiment were located in the mitochondria and plastids and many of those that decreased in abundance were located in the cytosol (Figure 2A). We then examined the spread of longevity across whole subcellular compartments and found that the histograms for mitochondria, plastids and extracellular locations were right skewed, indicating a greater proportion of proteins are longer lived in these compartments, most obviously at the two-week timepoint (Supplemental Figure S3A, B). By separating mitochondrial and plastid proteins from the bulk and observing the changes in longevity quotient distribution over time (Figure 2C), it appears that the distributions become increasingly right skewed as time progresses and the proteins become older. To confirm this observation, the spread of longevity quotients was compared across subcellular locations and there was found to be a significant location affect using ANOVA (p < 10^-15^; Figure 2C). Pairwise t-tests revealed that mitochondrial and plastid compartments contained proteins with significantly different longevity quotients, compared to the other 7 compartments. The difference between these two compartments and others was quite profound, with 29 % of mitochondrial proteins and 22 % of plastid proteins showing a long lifespan (*q* ≥ 1.6; Table 1). Twenty-one percent of extracellular AGIs were also long-lived, but this group was statistically inseparable from all other compartments, except the ER (Figure 2D, Table 1).

**Figure 2.**
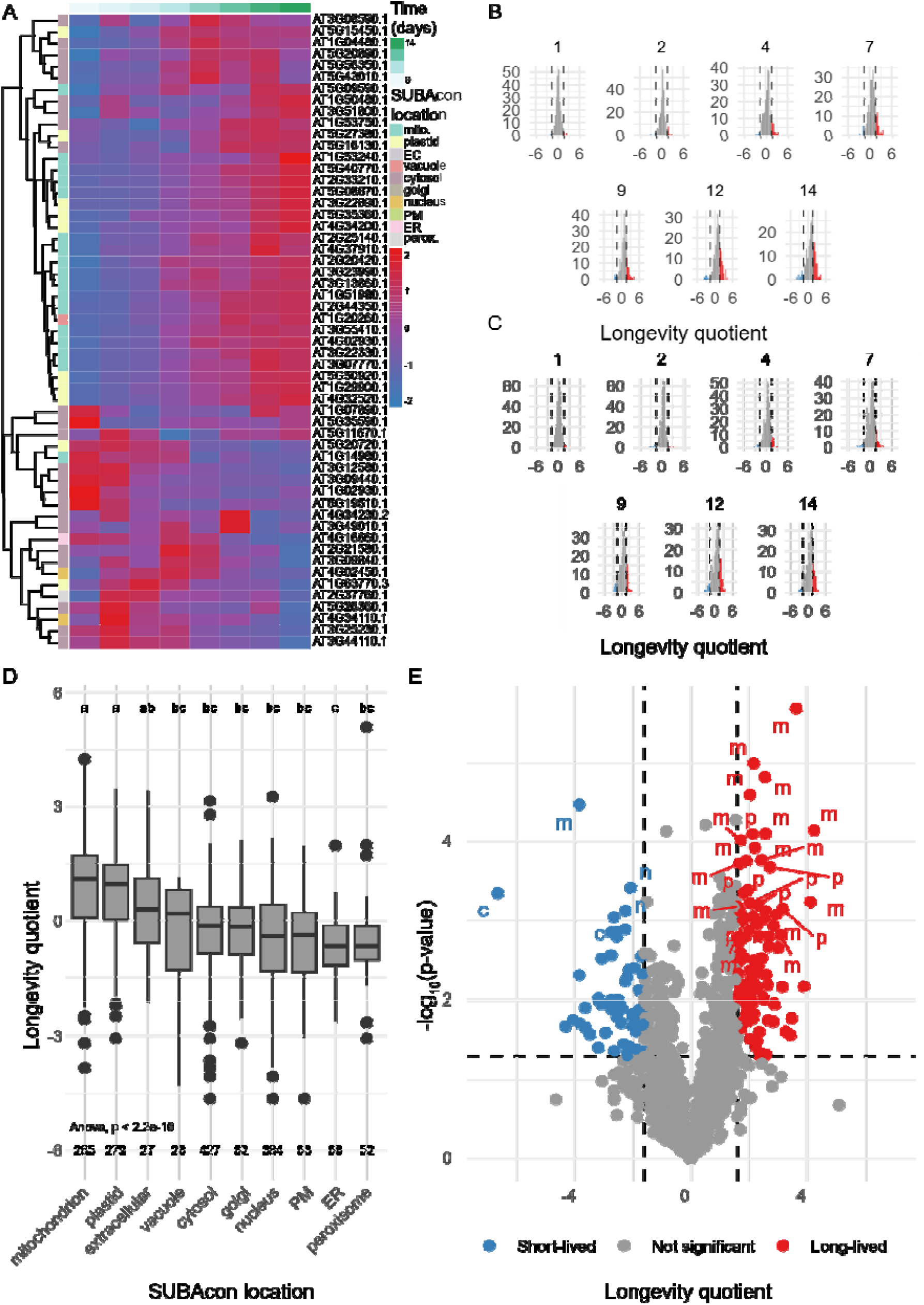
Longevity data for Arabidopsis cell culture proteins. (A) Heatmap of longevity quotients of selected highly abundant proteins (relative abundance ≥8 across all timepoints). (B) Longevity quotient for mitochondrial proteins, faceted by time. Dashed vertical lines are at –1.6 and 1.6 and indicate cutoff between short-lived (blue, *q* ≤ –1.6), moderate longevity (grey, –1.6 < *q* < 1.6) and long-lived (red, *q* ≥ 1.6) proteins. (C) Longevity quotient for plastid proteins, faceted by time. Dashed vertical lines as for B. (D) Boxplot and ANOVA of longevity quotients for proteins belonging to different subcellular locations (based on SUBAcon prediction). Letters indicate statistically different groups (pairwise t-tests, p-value ≤ 0.05), numbers indicate the number of proteins in subcellular location. (E) Volcano plot for all proteins. SUBAcon predicted locations (indicated by lowercase letters (m: mitochondrion, p: plastid, c: cytosol)) are shown for the 30 proteins with the most significant (lowest p-value) fold-change between the start and the end of the two-week chase. Dashed lines are at 1.3 (–log(0.05), y-axis) and –1.6 and 1.6 (x-axis). Proteins with p-values and longevity quotients outside above these parameters were considered short or long-lived as shown in blue and red.

**Table 1.** Summary statistics for proteomics data taken two weeks after switching the cell cultures back to normal media. Proteins were considered long-lived if q ≥ 1.6.

| Subcellular Location<br>(SUBAcon) | Mean longevity<br>quotient | Median longevity<br>quotient | Longevity quotient<br>standard deviation | Maximum longevity<br>quotient | Minimum longevity<br>quotient | Total AGIs | Long lived AGIs (%) |
| --- | --- | --- | --- | --- | --- | --- | --- |
| mitochondrion | 0.90 | 1.08 | 1.36 | 4.24 | -3.84 | 234 | 29 |
| plastid | 0.70 | 0.96 | 1.19 | 3.17 | -3.06 | 230 | 22 |
| extracellular | 0.37 | 0.29 | 1.44 | 3.42 | -2.12 | 19 | 21 |
| vacuole | -0.39 | 0.17 | 1.36 | 1.12 | -4.32 | 26 | 0 |
| cytosol | -0.37 | -0.23 | 1.17 | 3.13 | -6.64 | 331 | 2 |
| golgi | -0.19 | -0.15 | 0.98 | 2.1 | -3.18 | 59 | 3 |
| nucleus | -0.44 | -0.30 | 1.18 | 3.26 | -4.64 | 312 | 3 |
| PM | -0.55 | -0.35 | 1.39 | 1.96 | -4.64 | 40 | 8 |
| ER | -0.68 | -0.73 | 0.89 | 1.97 | -2.64 | 56 | 2 |
| peroxisome | -0.52 | -0.69 | 1.31 | 5.09 | -3.06 | 43 | 7 |

The same preponderance of mitochondrial and plastid proteins amongst the long-lived protein set could be observed by a series of other types of analyses. Volcano plot of longevity quotient revealed that most of the significantly longer-lived proteins were from mitochondrion, or plastid (Figure 2E). When only the mitochondrial and plastid proteins were graphed, 9 of the 10 most significantly differentially abundant proteins in these organelles were long-lived (Supplemental Figure S4A, C). GO term enrichment analysis of two-week longevity quotients of greater than 1.6 as a foreground list and the entire enriched dataset as the background found that the mitochondria and plastid GO terms were significantly enriched, especially true for mitochondria; 46/97 (47 %) of proteins in the foreground were mitochondrial compared to 248/1645 (15 %) in the background (Supplemental Figure S4B).

### Primary metabolism enzymes are longer lived and selected protein domains are associated with longevity

Analysis of the relative intensities of the most abundant proteins revealed distinct clusters of proteins that became more abundant over time and others that diminished over time. Parsing the data by MapMan functional groups showed proteins involved in primary metabolism commonly increased in relative abundance and those involved in protein biosynthesis, homeostasis and modification commonly decreased in abundance (Figure 3A). We statistically compared the differences in longevity between different MapMan functional categories (Figure 3B). Proteins involved in primary metabolism had a significantly longer lifespan than the basemean (i.e., the mean q-value across all MapMan categories). In contrast, proteins involved in RNA biosynthesis and processing, protein biosynthesis and homeostasis and protein modification and translocation, and ribosomal subunits had significantly shorter lifespans than the basemean. Long-lived proteins comprised 22 % of the primary metabolism group; in contrast, long-lived proteins comprised 2–11 % of the other groups. Given that proteins in the plastid and mitochondrion tended to be longer lived than proteins in other compartments, and they contain many primary metabolism enzymes, we wanted to determine if functional category or location was driving our observations. So, we separated by functional category and by subcellular location (Figure 3C). For most protein classes, proteins were longer lived in the mitochondria and plastids. This indicates that location may be more important than function when it comes to understanding protein lifespan.

**Figure 3.**
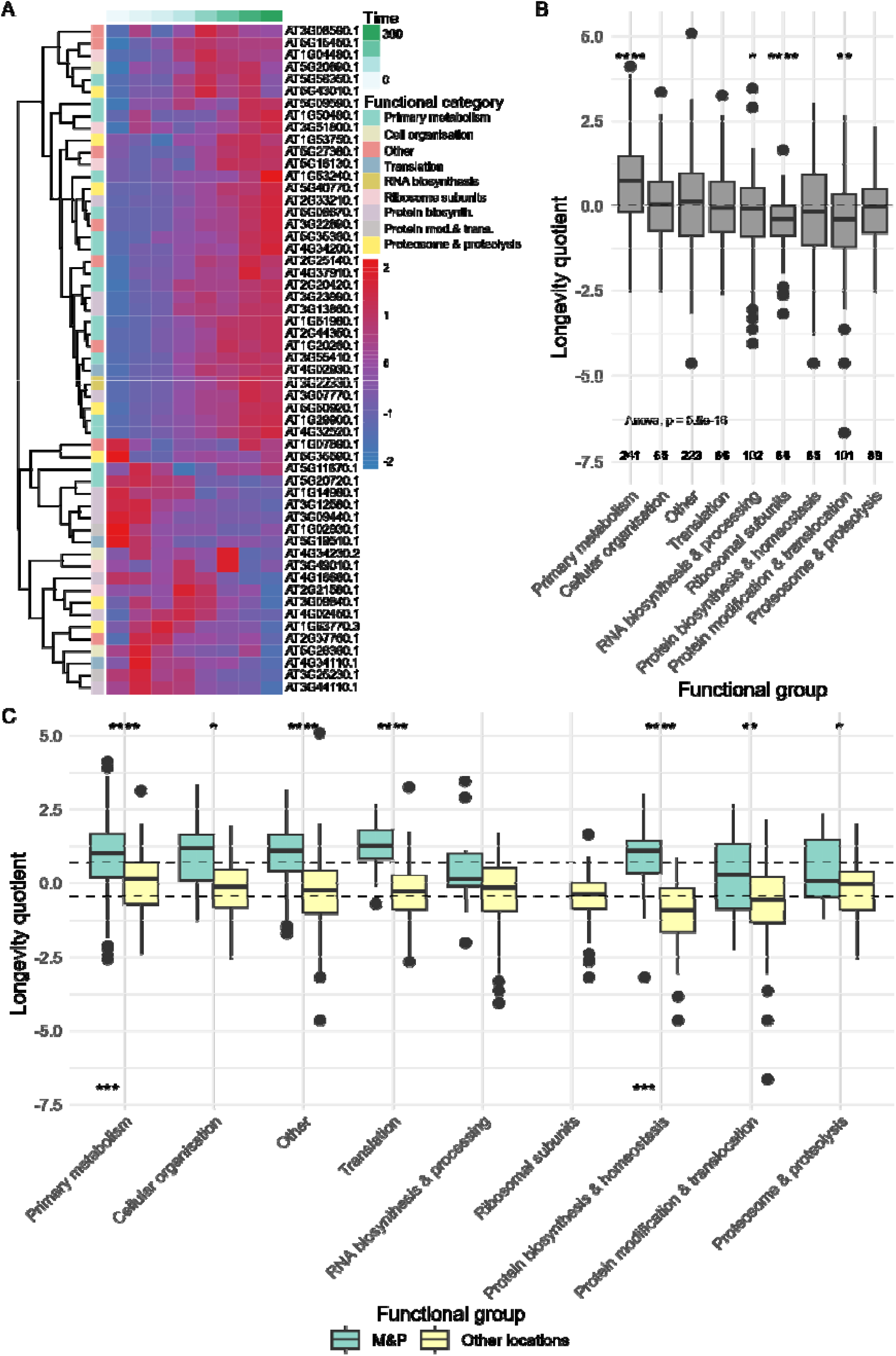
Longevity data for Arabidopsis cell culture proteins. (A) Heatmap of “longevity quotients” of highly abundant proteins (relative abundance ≥ 8 across all timepoints, as shown in Figure 2A) annotated for different MapMan functional categories. (B) Boxplot and ANOVA of longevity quotients for proteins in different MapMan functional categories. Statistically different groups (t-test) compared to the mean (dashed line) are denoted by asterisks. *p ≤ 0.05, **p ≤ 0.01, ***p ≤ 0.001, ****p ≤ 0.0001. (C) Longevity quotient for proteins in different functional categories in mitochondria and plastids (M&P), and all other cellular locations; dashed lines show mean longevity quotients for the two subcellular groups. Asterisks at the top indicate significantly different longevity quotients within functional categories between the two locations and asterisks at the bottom indicate significantly different longevity quotients between functional categories within each location. Significance marks as for B.

The longevity quotient was also found to correlate with certain Pfam and Interpro protein domains that appeared in 10 or more proteins in the datasets (Supplemental Figure S5A). The longevity quotients proteins belonging to PF00118 (TCP-1/cpn60 chaperonin family), PF01535 (pentatricopeptide repeat family; PPR) and PF13041 (PPR family) were significantly higher than the mean for the Pfams tested. Although this particular PPR subfamily does not have known functions, the PPR superfamily of proteins are well known for their functions in RNA processing in mitochondria and plastids (Barkan and Small, 2014) Interestingly, there was one Pfam that correlated with significantly lower longevity quotients than the basemean for the Pfams tested: PF00179 (ubiquitin-conjugating enzyme domain) which are central players in the trio of enzymes responsible for the attachment of ubiquitin (Ub) to cellular proteins to initiate proteosome linked degradation. Several Interpro domain also correlated with protein’s longevity (Supplemental Figure S5B). These included a pentatricopeptide repeat (PPR; IPR002885), which overlaps with the Pfam domains and a TIM β/α barrel domain (IPR013785). Some Pfam domains also correlated with shorter lifespans and most of these related to kinase activity, including a Serine/threonine-protein kinase active site (IPR008271), a protein kinase-like domain (IPR011009) and a protein kinase ATP binding site (IPR017441).

### Peptide spectra searches revealed changes in the proportion of post-translational modified peptides for proteins over time

We performed open mass searches of our MS/MS data with MSFragger to identify PTMs of peptides that could be correlated with protein aging (Supplemental Table S3). This revealed some PTMs which correlated strongly with age and others that showed weaker correlations (Supplemental Figure S6). We used these data to select PTMs for subsequent closed searches. Peptide N-terminal E-to-Pyro-Glu, W, M and C dihydroxylation and C and W oxidation were all selected for inclusion in the subsequent closed searches, along with other PTMs which we were interested in and suspected would change over time: M, K, P and T oxidation, K methylation and *N*-terminal acetylation (NTA).

To understand how protein modification changes over time, we selected a set of 1248 proteins that were consistently found in the enriched samples at each time point. This helped to eliminate the confounding effects of proteins (modified or not) that were present at only some timepoints. In this dataset, protein modification of all proteins generally increasesd over time, from 21 % after the 24-h pulse to 30 % after the two-week chase (Figure 4A). Amongst these, the long-lived proteins showed a greater PTM frequency compared to the bulk proteome and an increase from 21 % to 36 % over the two-week period (Figure 4B). In contrast, the short-lived proteins showed no correlation between age and PTM frequency (Figure 4C). This suggested that long-lived proteins not only accumulate more modifications as they age because they exist in the cell for longer, but that their modification in the cell may contribute to their longevity. The click reaction itself introduces a known degree of protein oxidation which will be evident following enrichment of HPG-containing proteins from samples (Meldal and Diness, 2020). The modification statuses of unenriched proteins in the control data showed lower percentages of modified spectra, consistent with this method-dependent oxidation (Supplemental Figure S7). Therefore, we focused our analysis and conclusions on PTMs in the test data that showed strong and significant trends up and down over time in the chase while the control data show no substantial change in PTM status during the chase period (Supplemental Figure S7). This represented cases where residue oxidation correlated with *in vivo* age of the samples being studied.

**Figure 4.**
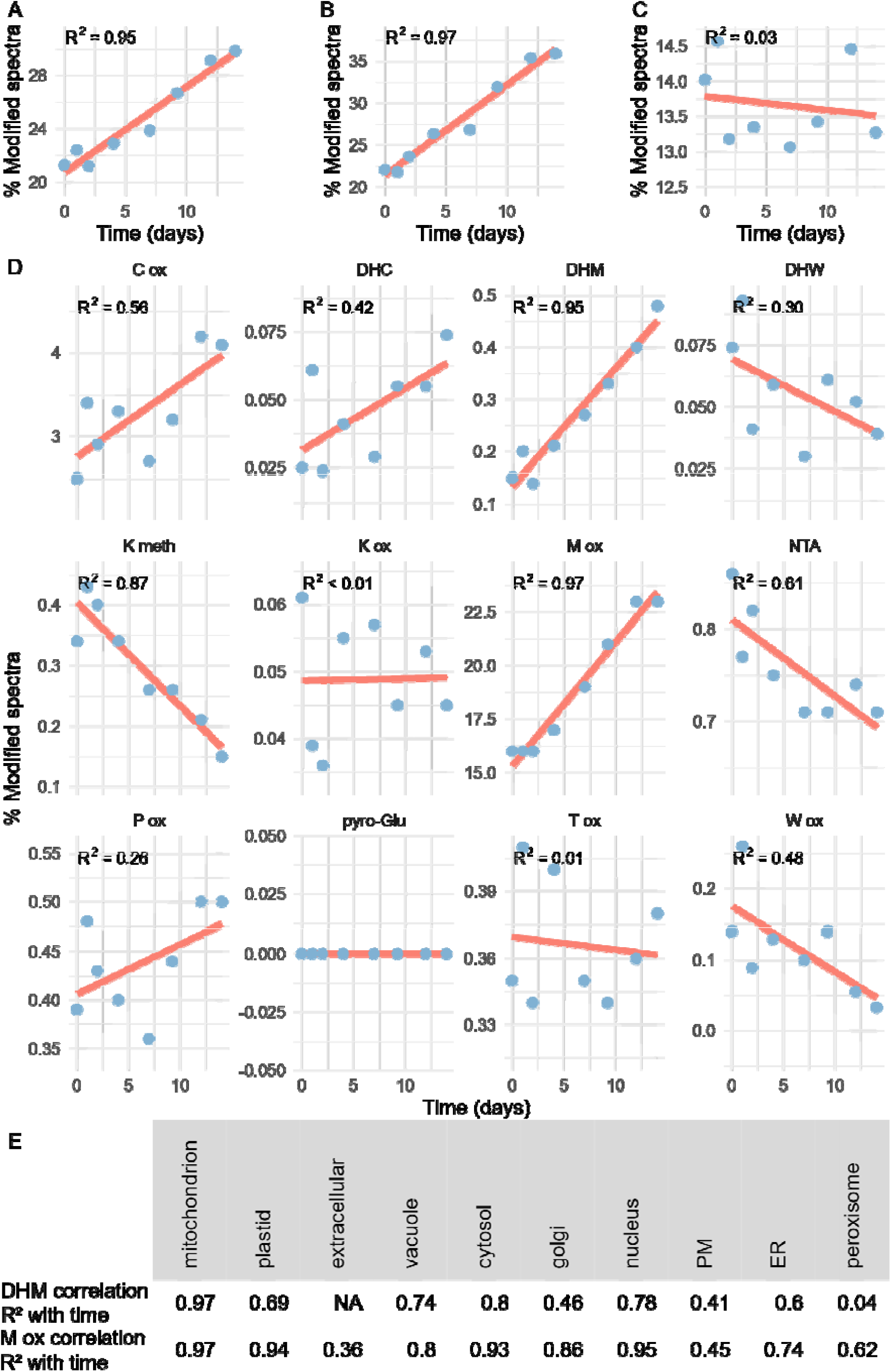
PTM frequency (total modified spectral count across three replicates relative to total spectral count across three replicates) for all selected proteins (A), long-lived proteins (B) and short-lived proteins (C) across the time series. PTM frequencies of different PTM types in selected proteins over time (D); correlations were significant (p ≤ 0.05) for C oxidation, DHM, K methylation, M oxidation and NTA; correlations were approaching significance (p ≤ 0.08) for DHC and W ox. Correlation (R^2^) between PTM frequency and time for different subcellular locations for DHM and M oxidation (E); DHM correlations were significant (p ≤ 0.05 for all locations, except extracellular, Golgi body (p = 0.07), PM (p = 0.09) and peroxisome (p = 0.62); M oxidation correlations were significant (p ≤ 0.05) in all locations, except extracellular and PM. Frequencies were calculated based on modified spectral counts, relative to total spectral counts. DHC: dihydroxycysteine, DHM: dihydroxymethionine, DHW: dihydroxytryptophan.

Breaking these time dependent relationships down into the different PTMs examined (Figure 4D), it was apparent that this increase was mainly in the form of oxidative modifications, implying that oxidation profile was the clearest marker of protein age. While M oxidation and dihydoxylation increased over time (R^2^= 0.97 and 0.95), other positively correlated oxidative modifications were less strong, with some, such as P oxidation and T oxidation, showing no discernible correlation. Other types of modification appear to be slowly or rapidly eliminated from the protein pool as it ages. The strongest negative correlation was between K methylation and age, that decreased 3-fold over 2 weeks. There was also a strong and significant correlation between K methylation and age in the control indicating that this effect may be due to the HPG treatment, rather than aging of the proteins (Supplemental Figure S7). However, in the control, the perfect fit make this correlation unreliable. Furthermore, the decrease in K methylation in the control was only 1.2-fold over 2 weeks, so we conclude that the decrease observed in the experimental data was likely due to a combination of HPG treatment and the aging of the proteome. NTA showed a moderate negative correlation over time, which is consistent with its co-translational nature. Once non-N-terminal pyroglutamine from E PTMs were eliminated from the dataset, no pyroglutamines remained, indicating this is unlikely to be biologically significant. W dihydroxylation and W oxidation showed weak to moderate negative correlations with protein age.

To better understand how PTM status changes as proteins age, we separated the PTM data by subcellular location, MapMan functional category, PFam ID and Interpro domains (see Supplemental Tables S4, S5, S7 for all correlations). Separating by subcellular location (Supplemental Table S4) revealed that M oxidation increased over time for proteins in all subcellular locations, except extracellular and PM (Figure 4E, Supplemental Figure S8A), with particularly strong and significant correlations observed in mitochondria, plastids, cytosol and the nucleus. This indicates that M oxidation is probably a chemical bioproduct of protein aging inside cells, rather a result of enzymatic modification to mark age. Another implication from this finding is that increasing M oxidation with protein age does not lead to decreased protein lifespan, especially since the level of M oxidation in mitochondria and plastids, where proteins tend to be longer lived, mirrors that of the cytosol and nucleus, where proteins have shorter lifespans. It was also observed that dihydroxymethionine (DHM) increased as proteins aged in all locations, except extracellular, Golgi body, PM and peroxisome. (Figure 4D, E). Interestingly, the correlation was moderate in the plastid, whereas it was strong for M oxidation in this compartment. The strongest correlation was seen in the mitochondria, where the percentage of DHM PSMs increased eight-fold over the two-week aging period (Supplemental Figure S8B). This is most likely because proteins in the mitochondria tend to live longer, and they are exposed to an environment containing reactive oxygen species and transition metals that are known to lead to methionine oxidation (Møller and Kristensen, 2004). The correlation for DHM was also strong in the cytosol, doubling over the two-week period. In the vacuole, the increase was also substantial (0–0.5 %) but the increase appeared non-linear over the sampling period.

When considering the MapMan functional categories of these proteins, we observed strong correlations between M oxidation frequency and time for all categories considered, except secondary metabolism (Supplemental Figure S9A). Similarly, DHM frequency was strongly and significantly correlated with time, especially for proteins in the translation, ‘protein biosynthesis & homeostasis’ and ‘protein modification & translocation’ groups (R^2^ ≥ 0.84). All correlations between PTM frequencies and time for MapMan functional categories are listed in Supplemental Table S5.

The frequency of M oxidation also increased over time for proteins grouped by the Pfams and Interpro domains. For proteins in the 17 Pfam groups previously investigated, M oxidation increased over time in 13 (Supplemental Figure S5, Supplemental Table S6). The correlations were strongest for PF00076, PF00226, PF00270 and PF17862. The correlation between M oxidation and time was also strong (p = 0.73) for PF01535, which is consistent with the long-lived nature of proteins with this Pfam (Supplemental Figure S5A). All correlations between PTM frequencies and time for PFam IDs are listed in Supplemental Table S6. Similarly, when parsing the data by Interpro domains, the correlation between M oxidation and time was observed for multiple Interpro domains (all correlations shown in Supplemental Table S7).

### Different aging characteristics of exemplar proteins

To more precisely understand the changes to PTM status as proteins age, we selected exemplar proteins with different characteristics for additional analysis. Four of these provide a range of the data and illustrate key features of the differences observed (Figure 5).

**Figure 5.**
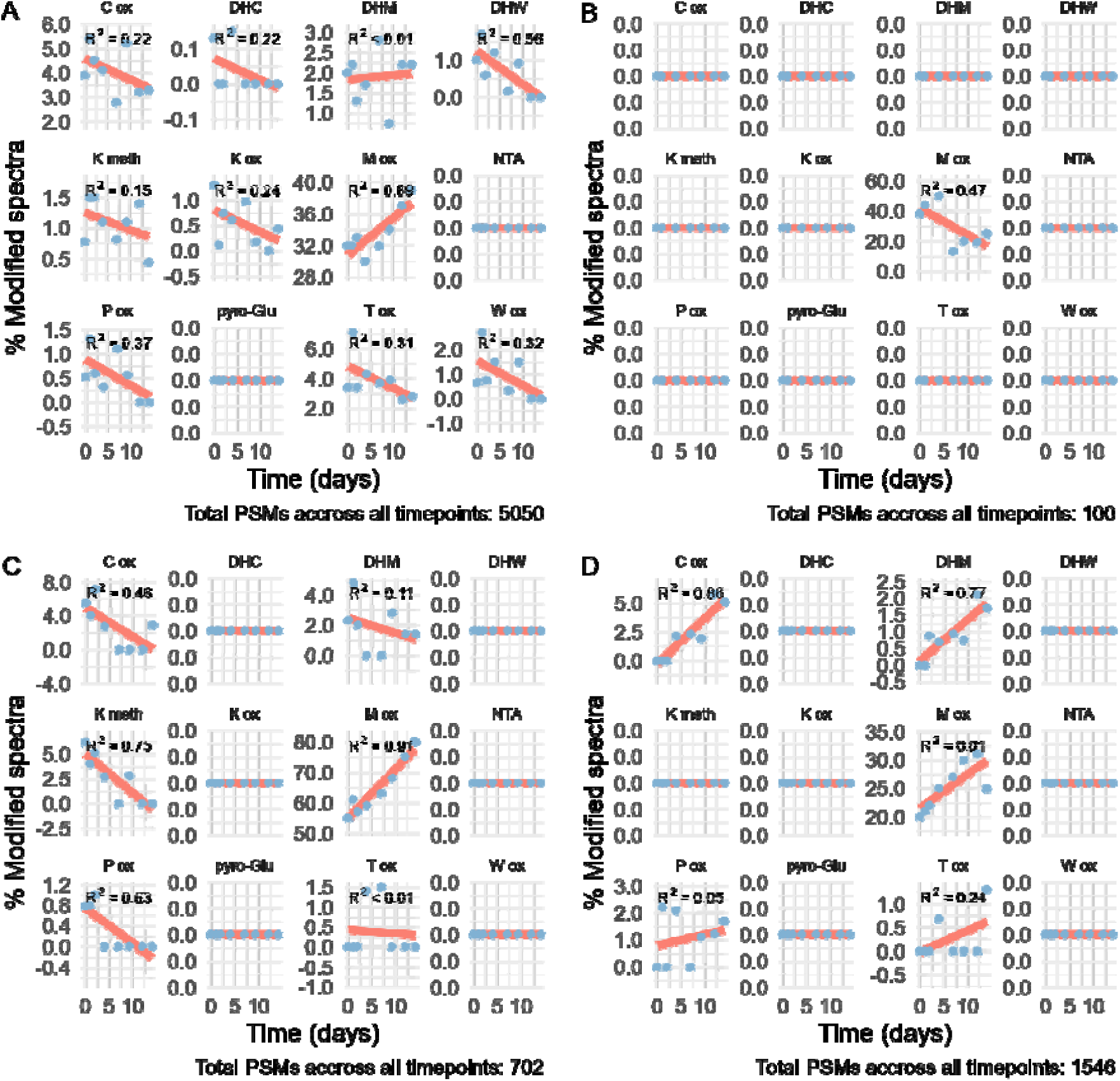
PTM frequencies of peptides for selected proteins across the time series. (A) HEAT SHOCK PROTEIN 70 (HSP70, AT3G12580.1); (B) CINNAMYL ALCOHOL DEHYDROGENASE 1(CAD1, AT1G72680.1); (C) thiazole synthase (THI4, AT5G54770.1); (D) NADH-dependent glutamate synthase 1 (GLT1, AT5G53460.1). Frequencies were calculated based on total modified spectral counts (across three biological replicates), relative to total spectral counts (across three biological replicates). DHC: dihydroxycysteine, DHM: dihydroxymethionine, DHW: dihydroxytryptophan.

The protein with the most PSMs in the entire experiment was HEAT SHOCK PROTEIN 70 (HSP70; AT3G12580.1), a cytosolic ATP-dependent chaperone involved in cellular protein quality control and degradation regulation (Usman et al., 2017). Interestingly, despite its high number of spectral counts, HSP70 was shown to be short-lived in this experiment, with a 60 % reduction in relative abundance over the course of the experiment (Supplemental Table S1). HSP70 showed PTM trends consistent with its short-lived nature: weak negative correlations were observed for most of the PTMs in our data (Figure 5A). Notable exceptions included pyro-Glu from E and NTA, which were not detected, DHC, which showed a weak positive correlation, DHW, which showed a moderate negative correlation and M oxidation, which showed a moderate positive correlation. Interestingly, other proteins in the ‘protein biosynthesis and homeostasis group’ showed a wide range of longevities, with longer-lived proteins generally being found in the mitochondria or plastids. This further demonstrates that the longevity of a protein is primarily determined by its location, rather than its function.

The longest-lived protein in the dataset was CAD1; CINNAMYL ALCOHOL DEHYDROGENASE 1 (AT1G72680.1). It is a member of a nine-member family of NADPH-dependent enzymes in Arabidopsis involved in final step of monolignol biosynthesis and known to be expressed in lignifying tissues (Kim et al., 2007). In the enriched protein samples, this protein increased in relative abundance 34-fold over the two weeks of the experiment. It has been shown to be a cytosolic protein (Ito et al., 2011), but it can adhere, presumably to the outside, of mitochondria (Fuchs et al., 2020). In contrast to general trends, this enzyme showed a weak positive correlation between M oxidation and age (Figure 5B), despite the protein containing nine M residues. This age-dependent PTM profile contrasts with the general trend for long-lived proteins and is indicative of the diversity amongst the different protein groups.

One of the most intriguing proteins in the data was THI4; a plastid localised thiazole synthase enzyme (AT5G54770.1). THI4 uses a suicidal reaction mechanism in the synthesis of thiazole, a precursor to thiamin (vitamin B_1_) (Amthor et al., 2019), and the enzyme had a 3-fold reduction in relative abundance over the course of the experiment (q = –1.6). In our dataset, THI4 was one of the most highly post-translationally modified proteins. There was a strong negative correlation between THI4 age and K methylation and a very strong positive correlation between THI4 age and M oxidation (Figure 5C). Since THI4 becomes non-functional after catalysing one reaction, the THI4 present in our samples contains a greater and greater portion of nonfunctional enzyme, assuming that old and new proteins are degraded at the same rate and the cell cannot sense which enzyme molecules are functional and which are not.

Another contrasting example is NADH-dependent glutamate synthase 1 (GLT1; AT5G53460.1), a long-lived plastid localised enzyme. This protein contains at TIM β/α barrel domain (IPR013785) found in aldolase proteins. Our data contained 23 proteins with this domain, and it correlates with longevity. Compared to the short lived THI4, the modification patterns over time are quite different. C oxidation in THI4 is weakly negatively correlated with time, whereas for GLT1, there is a very strong positive correlation. K methylation decreases in THI4 but is not present in GLT1 (Figure 5C, D). M oxidation increases over time, but the percentage of modified spectral counts and the strength of the correlation is much higher in THI4 than in GLT1. This may indicate that the cell makes no effort to reverse M oxidation in THI4 because it has a short useful longevity, and a high level of M oxidation indicates that a high proportion of the enzyme is old and therefore likely nonfunctional. DHM patterns also varied between the two exemplars. DHM did not correlate with age in THI4 (Figure 5C) but there was a strong correlation with age in GLT1 (Figure 5D). Because GLT1 remains longer in the cell, there is potentially more time for M oxidations to be further oxidised to DHM. This observation may also help explain why the M oxidation correlation over time was weaker for GLT1—because oxidised M residues are further oxidised to DHM.

K-means clustering was used to identify nine protein groups that had similar modification patterns over time (Supplemental Figure S10). The exemplar proteins (Figure 5) are members of four of these groups. The group memberships are shown in Supplemental Table S8 Comparison of subcellular location, functional categories and Pfam and Interpro domains did not clearly identify a unifying factor that explained the protein groups (χ^2^ p-value ≥ 0.21 and Cramer’s V ≤ 0.28 in all cases), indicating complex factors contribute to the PTM profile of different types of protein as they age.

## Discussion

### Kinetics of protein delivery and installation in different organelles

Protein turnover is thought to be a product of multiple interacting factors, including the presence or absence of certain degrons (Isono *et al*., 2024), the presence or absence of specific domains (Li *et al*., 2017), membership of a complex (Li *et al*., 2017), organism age (Basisty et al., 2018; Dhondt et al., 2017), environmental factors (Fan et al., 2024; Li et al., 2022b), phytohormone signalling (de Roij et al., 2024) and PTM status (Xu et al., 2024). Arraying our longevity data as subcellular proteome sets showed that relative to the cytosol, the old proteome accumulated in mitochondria, plastids and outside the cell in extracellular spaces, but depleted in the ER and peroxisomes. This is generally consistent with a broad understanding of protein trafficking is cells. Plastids and mitochondria are final destinations with their own protein degradation machinery. Nuclear-encoded organelle targeted proteins are sent to these organelles, functionally incorporated into machinery and will be degraded in time through the independent ATP-dependent proteases of FtsH, Clp and Lon families (van Wijk, 2015). Given the sheer bulk of the cellular proteome in these organelles, especially plastids, it is imperative that they represent relatively slow sites of degradation to conserve cellular energy used in protein synthesis/degradation cycles. The extracellular matrix is also a final destination and often contains tough enzymes and structural proteins that can operate without chaperones, limited cofactors and in adverse and changing osmotic and pH conditions. In contrast the ER is the chief highway of the secretory system with a role to delivery proteins rather than accumulate them. An exception in our data might be seen in peroxisomes that on average were actively depleted from the proteome across the two-week period (Figure 2D). Peroxisomes are sometimes considered along with mitochondria and plastids in metabolic discussions as similar structures. But, in fact, peroxisomes have a complex relationship with the ER (Geuze et al., 2003), can arise from the ER, house a range of oxidising reactions in plants that might necessitate regular renewal (Pan et al., 2020) and are often seen as selective targets for autophagy, known as pexophagy (Goto-Yamada et al., 2023). Overall, the data shown here quantify this proteome latency dynamics across the cell sub-structure in a new way.

In some respects, it is unsurprising that proteins in the mitochondria and plastids tend to be longer lived, since proteins in these organelles are sheltered from the proteosome. However, recent evidence suggests that the ubiquitin-proteosome system (UPS) for degradation of proteins may have a role to play even in plastid-localised protein degradation (Jarvis et al., 2024), so being sheltered from the proteosome may not be the only reason we observed significant stability differences in mitochondrial and plastid proteins compared to those in the rest of the cell. Inside mitochondria and plastids, prokaryotic-type proteases (Huang et al., 2020) facilitate degradation of target proteins through recognition of degrons in their targets or delivery of target proteins to catalytic subunits (Isono *et al*., 2024). Mitochondrial and plastid proteins can also be degraded by autophagy (Li et al., 2022a). According to our data, these systems have higher tolerances for aging proteins than the UPS system. This means that overall, the mitochondrial and plastid proteomes are older than the proteomes of the cytosol, nucleus and all the other compartments examined in this study. Nevertheless, it has been shown that mitochondrial and plastid proteins and proteins in other subcellular compartments all display a similar increase in degradation rate upon exposure to certain stressors, such as high light (Li *et al*., 2022b), which may indicate shared signalling pathways for protein degradation induction.

Although proteins in mitochondria and plastids tend to be longer lived, within these organelles, proteins are turned over selectively, indicating that location is not the only driver of protein longevity, although our data indicate that it is a major driver (Figure 3C). Another factor that appears to correlate with longevity is MapMan functional category. While longevity quotients were significantly higher in mitochondria and plastids compared to the rest of the cell for all MapMan functional categories tested, except the ‘RNA biosynthesis and processing’ category, the function of proteins correlated with differences in longevity within these two groups of compartments. In the plastids and mitochondria, primary metabolism proteins were not significantly longer lived compared to the baseline mean for these compartments. However, in the non-plastid/non-mitochondrial compartments, primary metabolism proteins showed significantly increased longevity, indicating that location and function both affect their longevity. In contrast, proteins involved in protein modification and translocation were significantly shorter lived than the respective baseline means within the two compartment groups, although the proteins in this functional category in the mitochondria and plastids were significantly longer lived than those in the non-plastid/non-mitochondrial group. Within the non-plastid/non-mitochondrial group, proteins involved in protein biosynthesis and homeostasis were significantly shorter lived compared to the baseline mean, but in the mitochondria and plastids, these proteins longevity quotients did not differ from the baseline mean. Overall, our data show that there is more consistency in protein longevity within the mitochondria and plastids compared to the rest of the cell and that proteins in all MapMan functional categories display the same or significantly longer lifespans in these organelles compared to the rest of the cell. Ribosomal subunits showed shorter lifespans than the baseline mean when all the compartments were considered, but this difference vanished when mitochondria and plastids were removed from the analysis (compare Figure 3B and 3C). Consistent with previous studies (Salih et al., 2020), our data show that ribosomal subunits tend to have similar longevity, with 87 % having moderate longevity (–1.6 < q < 1.6; Supplemental Table S1). Notable proteins in this group that had longer, or shorter, lifespans were a core component (Gar1) of the H/ACA small nucleolar ribonucleoprotein RNA pseudouridylation complex (AT3G03920.1, *q* = 1.66) and NOC3, a component of the ribosomal subunit nuclear export complex (AT1G79150.1, q = –3.18).

In the same way as location and function are correlated with differences in longevity, certain Pfams and Interpro domains also correlated with increased or decreased longevity. However, closer inspection revealed that these were mostly a result of location rather than the domains themselves. Two exceptions to this trend were the TIM β/α barrel domain (IPR013785) and AAA+ ATPase domain (IPR003593).

### Linking PTMs and organelle functions to degradation

A range of PTMs were found in the proteome as it aged (Figure 4D, Supplemental Figure S6). Abundant PTM processes in plants can be viewed as early, such as N-terminal acetylation (NTA) that is typically co-translational, as structural such as may other types of acetylation and methylation, as regulatory such as phosphorylation, or as cumulative with time or function such as oxidation (Millar et al., 2019). Various methylations and acetylations were identified but most did not show clear responses to aging (Figure 4, Supplemental Figure S6). An exception was K methylation that was depleted 3 fold between young to old protein sets (Figure 4D). K methylation causes minimal changes in the size and electrostatic status of K residues but there are reports in a range of organisms methyllysine reader proteins that can selectively bind to them, enabling recognition. These are best exemplified by the methylation and demethylation of histones (Luo, 2018). M, C oxidations accumulated with age, as did DHM and DHC (Figure 4D), indicating that protein molecules containing them were not selectively. or at least not efficiently, removed by degradation machinery from the protein pool. But this was not the case of W oxidation, which depleted in old proteome samples (Figure 4D). Intensive study of W oxidation has discovered how it can structurally alter proteins enabling them to be recognised for degradation (Ehrenshaft et al., 2015). In plants W oxidation of P43 and D1 core polypeptides of PSII have been shown to be a key signal for photosynthetic damage repair through protein turnover of selected D1 polypeptides (Kasson and Barry, 2012).

### Implications for studying changes in PTM abundance in plant proteomes

PTMs are a ‘cost-effective’ way of modulating protein function (Mehta et al., 2021) because they do not require degradation of an unneeded protein and subsequent resynthesis. Some PTMs are acquired by enzymatic catalysis but others are enzyme-independent chemical reactions (Millar *et al*., 2019). Here, we used open searches to identify 12 PTMs that appeared to modulate with protein ages. These PTMs were mostly oxidative (nine of the 12). Notably absent from the list generated by the open searches were common PTMs, such as phosphorylation, glycosylation, ubiquitination and nitrosylation (Bobalova et al., 2023). This indicates that these modifications did not modulate with protein age with any discernible pattern at a proteome level, although the degree of phosphorylation on specific proteins has been previously linked with plant organ age (Rankenberg et al., 2024).

Using closed searches containing the 12 PTMs suggested by open searches to be age-dependent, we generated time-resolved proteomics data, allowing evaluation of the PTM status of proteins as they age. We showed that proteins generally accumulate PTMs as they age but the rate of PTM accumulation and the nature of the PTMs accumulated vary depending on lifespan, subcellular location, function and presence of specific domains.

Unexpectedly, the number of PTM-containing PSMs and the rate of PTM accumulation varied with protein longevity. Longer-lived proteins tended to accumulate more PTMs over their lifetimes and the rate of accumulation was greater for these proteins. PTM accumulation did not correlate with age for shorter lived proteins. If accumulation of PTMs was merely a chemical by-product of the cellular environment, the rate of PTM accumulation should be the same for all proteins in the same subcellular compartment, regardless of the longevity of the proteins in question. But this is not what we found; in all compartments, shorter-lived proteins did not accumulate PTMs in a consistent way as the proteins aged, but longer-lived proteins did. Given that this accumulation of PTMs is mostly oxidative, the most likely explanation for this observation is that cells remove oxidations on shorter lived proteins, but they do not remove these PTMs on longer-lived proteins. This implies that the removal oxidation from C, M, P, K, T and/or W residues contributes to protein turnover.

Our observations in the aging proteome also indicate that the type of sample and the growth rate of the tissue should be taken into consideration when interpreting apparent changes in PTM abundances. For example, abiotic stress studies, that are very popular to profile in PTM research, are typically associated with slowing or cessation of plant growth rate over a period of days. This will incrementally age the protein complement through the decrease in the ability of protein synthesis to dilute the pre-existing protein pool. Accumulation of PTMs in such scenarios thus has a temporal component associated with growth rate, rather than only a plant response to the stress imposed. Oxidation of proteins is often found in abiotic stress studies and attributed to oxidative damage caused by the stress. But our data show that over a few days without new protein replacement, an aging proteome can deliver the same effect without any specific oxidative stress having been induced.

### Designing efficient and effective SynBio systems

Synthetic biology (SynBio) is a burgeoning discipline that applies engineering principles to biology and is focused on developing useful biological parts, devices and systems. Examples of SynBio devices and systems include biosensors (Jones et al., 2019), memory circuits (Lloyd et al., 2022) and plants with improved energy use efficiency (South et al., 2019).

SynBio parts are well-characterised biomolecules with defined functions and includes promoters, repressors and enzymes. Parts are assembled into devices and devices can be further assembled into complex systems. When designing SynBio systems, engineers select or create parts that are encoded by certain genes that are designed to work together to produce and efficient and effective system. However, there has been comparatively little research into the post-translational fates of the protein products of recombinant genes. Our understanding of the varying longevity of proteins and what drives it is limited, as is our understanding of post-translational structural changes. Understanding what happens to these proteins over the course of their lifetime is of paramount importance when designing synthetic biology systems because it can impact the structure and functioning of the recombinant proteins.

The ideal recombinant protein for a SynBio system would be one that is long lived or could have its life extended by optimal localisation but does not accumulate modifications that lead to structural changes. While many of the mitochondrial and plastid proteins displayed the former characteristic, they did not display the latter, generally being subjected to high levels of oxidation as they aged. However, there were some long-lived proteins, such as CAD1, that did not accumulate modifications during aging. Learning the best locations and the folds of proteins that resist modification would make excellent SynBio parts in terms of efficiency and effectiveness. Equally, the techniques used here can be deployed in SynBio systems to study the fate of specific proteins or interest and provide detailed information on the nature of components aging to engage design changes to optimise functional protein longevity.

## Methods

### Arabidopsis cell culture

PSB-D cell cultures were maintained on sterile MSMO media, which contained: sucrose (30 g/L; Chem-Supply, Gillman, SA, Australia), 4-morpholineethanesulfonic acid (0.5 g/L; Astral Scientific; Taren Point, NSW, Australia), Linsmeier & Skoog Basal Medium (4.43 g/L; PhytoTech Labs, Shawnee Missions, KS, USA), α-naphthaleneacetic acid (0.5 mg/L; Sigma, St. Louis, MO, USA) and kinetin (0.05 mg/L; Sigma); the pH was adjusted to 5.7 with aqueous KOH, and the media was sterilised by autoclaving before use.

### BONCAT pulse-chase

Five five-day-old cell cultures were pulsed with HPG as follows: cultures (120 mL each) were sampled (2 x 5 mL per culture) and the cells were collected by centrifugation (3 x 10^3^ RCF, 2 min) followed by aspiration of the supernatant and snap-freezing of the pellet in liquid N_2_. Cultures were treated with HPG by addition of 1.1 mL of HPG solution (100 mM in water, filter-sterilised). After 6 h, the cultures were sampled, as above (1 x 5 mL per culture). After 24 h, the cultures were again sampled, as above and then washed with and transferred to fresh MSMO media (volume of new media equivalent to the remaining volume before the media switch). The cultures were then sampled by centrifugation 1, 2, 4, 7, 9, 12 and 14 days after the beginning of the chase period (5 mL per culture each time, except at 14 days, when 4 mL was collected for each replicate). At each time point, a 1-mL sample of each replicate was taken for viability staining using propidium iodide and fluorescein diacetate followed by microscopic analysis (Supplemental Figure S11). Fresh weights were obtained for each sample (Supplemental Figure S12).

### Enrichment of alkyne-modified proteins

Three replicate samples from each timepoint were selected for azide-alkyne cycloaddition enrichment as follows. Tissue samples (200–1000 mg) were lyophilised, homogenised and extracted essentially as in Tivendale *et al*. (2021; see the Protein extraction and LC-MS preparation sub-section), vortexing frequently and repeatedly over the course of an hour for extraction using 0.8–2.3 µL/mg (FW) of extraction buffer, up to completion of the acetone wash steps. These acetone-washed protein pellets were resuspended in 100–200 µL of Lysis Buffer (made up as per the Click Chemistry Tools Click-&-Go Protein Enrichment Kits (Product number 1153, alkyne-modified protein capture) protocol) and the protein concentrations determined using Amido Black assays. The click reaction was performed essentially as described in the protocol, except the reaction mixtures contained 800 µL of protein solution in Lysis Buffer (containing 500 µg of protein), instead of 800 µL of cell lysate. At the completion of the protocol, enriched peptide samples were dried under reduced pressure and stored at –80 or –20 °C until they could be analysed.

### Control BONCAT pulse-chase

To account for proteomic changes induced by the HPG pulse-chase, a separate labelling experiment was performed, essentially as described above, but the samples were not subject to the enrichment procedure. Instead, they were digested and analysed directly by LC-MS.

### Peptide LC-MS

Peptide samples were analysed by UHPLC-MS, as follows. Unless otherwise stated, the conditions were the same for experimental and control peptide samples. Peptide samples were injected into an online nanoflow (0.225 µL/min for experimental samples and 0.3 µL/min for control samples) capillary column (Picofrit with 50 μm tip opening/75 μm diameter, New Objective, ICT36015030F-50) packed in-house with 15cm C18 silica material (3 µm; Dr. Maisch GmbH) connected to Thermo Exploris 480, in-line with a Dionex Ultimate 3000 series UHPLC. The column flow rate was 0.225 mL/min for experimental samples and 0.3 mL/min for control samples, with the following elution profile: 0–1 min % v/v acetonitrile, 0.1 % v/v formic acid to 2 % v/v acetonitrile, 0.1 % v/v formic acid; 1– 70 min 2 % v/v acetonitrile for experimental samples and 5 % v/v for control samples, % v/v formic acid to 23 % v/v acetonitrile, 0.1 % v/v formic acid; 70–78 min 23 % v/v acetonitrile, 0.1 % v/v formic acid to 35 % v/v acetonitrile, 0.1 % v/v formic acid; 78–80 min 35 % v/v acetonitrile, 0.1 % v/v formic acid to 95 % v/v acetonitrile, 0.1 % v/v formic acid; 80–82 min 95 % v/v acetonitrile, 0.1 % v/v formic acid, 82–83 min 95 % v/v acetonitrile, % v/v formic acid to 2 % v/v acetonitrile, 0.1 % v/v formic acid; 83–105 min for experimental samples and 83–90 min for control samples 2 % v/v acetonitrile, 0.1 % v/v formic acid. Spray voltage was set to 2 kV, and heated capillary temperature was at 275 °C. DDA was used, and full MS resolutions were set to 60,000 at m/z 200 and full MS AGC target was 300 % with an auto injection time. Mass range was set to 350–1500. AGC target value for fragment spectra was set to ‘standard’ with a resolution of 15,000 and auto injection time. Intensity threshold was kept at 5E4. Isolation width was set at 1.6 m/z. Normalized collision energy was set at 30%.

### Data processing

Before processing, Thermo.raw files were converted to mzML using the Msconvert package (v3.0.1908-43e675997; peakPicking filter vendor msLevel = 1) and the resulting files were analysed using MSFragger run in the FragPipe wrapper (Kong et al., 2017). Peptide mass spectra were searched against the Araport11 database. Results were checked using the For time-resolved analysis of proteomics data from cultures that had been treated with HPG, open searchers were performed and Philosopher (da Veiga Leprevost et al., 2020) and PTM Shepherd (Geiszler et al., 2021) were used to identify mass shifts in open searches; this allowed identification of mass deltas (representing PTMs) that varied with increasing protein age. After identifying these data, modifications of interest were included closed searches of the data. For our purposes, to be included in the subsequent closed searches, mass deltas had to meet the following criteria: must have at least 1000 PSMs in the data, must be a known modification, must not be in combination with isotopic peaks or isotopic peak errors, must be organic, must make logical sense (e.g. 2-amino-3-oxo-butanoic acid cannot logically replace tryptophan or cysteine). Potentially modified residues were selected on interest and probability score (if the probability score for one residue was ≥ 10x that of the rest, then the others were ignored). M acetylation was not considered since it was anticipated that this would be captured in the N-terminal acetylation (NTA) variable modification. Graphs were used to determine which mass deltas changed significantly over time.

Variable modifications included in closed searches were M, C, P, K, W, and T oxidation (+15.9949), protein NTA (+42.0106, NTA), peptide *N*-terminal pyro-Glu from Glu (–18.0106), W, M and C dihydroxylation (+31.9896), K methylation (+14.0157), and M-to-HPG substitution (–21.9877). These results were also checked using the Philosopher program (da Veiga Leprevost *et al*., 2020) and then further analysed with an in-house R-script (available on GitHub; https://github.com/ndtivendale/oldProteome). Proteins that showed ≥ 3-fold or ≤ 0.3-fold relative abundance fold-change in the control BONCAT pulse-chase over the course of the experiment were omitted from the analysis of the test samples to remove any proteomic changes induced by the HPG pulse.

Percent modified spectra were calculated on a per protein basis in two ways: 1) based on total spectral counts containing each modification across the three replicates for each time point relative to the total spectral counts and 2) as for 1, except relative the total spectral counts that contained potentially modifiable residues.

## Data availability

The MS proteomics data were deposited to the ProteomeXchange Consortium via the PRIDE partner repository (Perez-Riverol et al., 2019) with the dataset identifiers PXD062151 (main experiment) and PXD062152 (control).

## Author contributions

NDT and AHM conceived of the project and designed the experiments. NDT conducted the main experiments and analysed the data. XL performed the control experiment and NDT analysed the control data. NDT and AHM wrote the manuscript.

## Acknowledgements

Peptide quantitation in this work was performed as a service by the WA Proteomics Facility as a node of Proteomics Australia, supported by infrastructure funding from the Western Australian State Government in partnership with Bioplatforms Australia under the Commonwealth Government National Collaborative Research Infrastructure Strategy

