## Supplemental Information for "Analysis of protein aging reveals rates of subcellular organelle renewal and selective post-translational modification in Arabidopsis"

Supplemental Table S3. Mass shifts with >1000 PSMs from open searches.

| Mass shift | No. PSMss | Chemical identity | Most likely residue | 2^nd^ most likely residue | 3^rd^ most likely residue |
| --- | --- | --- | --- | --- | --- |
| -18.0106 | 1746 | Dehydration/Pyro-glu from E | E | T | D |
| -17.0272 | 6115 | Pyro-glu from Q/Loss of ammonia | Q | C | H |
| -2.015 | 1604 | 2-amino-3-oxo-butanoic acid | W | C |  |
| -1.031 | 1017 | Lysine oxidation to aminoadipic semialdehyde | C | H | E |
| 0.9842 | 13033 | Deamidation | N |  |  |
| 12.0004 | 1239 | formaldehyde adduct | W | H | N |
| 14.0152 | 2999 | Methylation | K | P | D |
| 15.9946 | 20651 | Oxidation or Hydroxylation | C | W |  |
| 27.995 | 3702 | Formylation | H | S |  |
| 28.0316 | 2324 | di-Methylation/Acetaldehyde +28/Ethylation | K | A | F |
| 31.9896 | 4637 | dihydroxy | W | M | C |
| 42.009 | 2474 | Acetylation | M | K |  |
| 42.0476 | 1033 | tri-Methylation/Propyl | K | P |  |

Supplemental Table S4. Correlations between PTM frequencies and time for proteins in different subcellular (SUBAcon) locations.

| PTM | SUBAcon location | R^2^ | Slope | p |
| --- | --- | --- | --- | --- |
| C dihydroxylation | mitochondrion | 0.61 | 7.14E-03 | 0.02 |
| C dihydroxylation | plastid | 0.4 | 1.88E-03 | 0.09 |
| C dihydroxylation | extracellular | NA | 0.00E+00 | NA |
| C dihydroxylation | vacuole | NA | 0.00E+00 | NA |
| C dihydroxylation | cytosol | 0.07 | 1.47E-03 | 0.53 |
| C dihydroxylation | golgi | 0.1 | -1.49E-03 | 0.44 |
| C dihydroxylation | nucleus | 0.23 | -1.46E-03 | 0.23 |
| C dihydroxylation | PM | NA | 0.00E+00 | NA |
| C dihydroxylation | ER | 0.08 | -1.99E-03 | 0.49 |
| C dihydroxylation | peroxisome | 0.23 | -2.50E-03 | 0.23 |
| C oxidation | mitochondrion | 0.8 | 1.51E-01 | 0 |
| C oxidation | plastid | 0.66 | 1.29E-01 | 0.01 |
| C oxidation | extracellular | 0.02 | -1.80E-02 | 0.74 |
| C oxidation | vacuole | 0.02 | 1.91E-02 | 0.75 |
| C oxidation | cytosol | 0.36 | 6.78E-02 | 0.11 |
| C oxidation | golgi | 0.09 | -1.88E-02 | 0.48 |
| C oxidation | nucleus | 0.5 | 4.12E-02 | 0.05 |
| C oxidation | PM | 0.3 | 7.63E-02 | 0.16 |
| C oxidation | ER | 0.18 | -3.09E-02 | 0.29 |
| C oxidation | peroxisome | 0.12 | 3.72E-02 | 0.39 |
| K methylation | mitochondrion | 0.07 | -4.03E-03 | 0.54 |
| K methylation | plastid | 0.85 | -2.05E-02 | 0 |
| K methylation | extracellular | NA | 0.00E+00 | NA |
| K methylation | vacuole | 0.67 | -3.73E-02 | 0.01 |
| K methylation | cytosol | 0.83 | -2.85E-02 | 0 |
| K methylation | golgi | 0.11 | 8.43E-03 | 0.42 |
| K methylation | nucleus | 0.58 | -1.35E-02 | 0.03 |
| K methylation | PM | 0.04 | -1.44E-03 | 0.63 |
| K methylation | ER | 0.01 | -1.96E-03 | 0.84 |
| K methylation | peroxisome | 0.15 | 6.16E-03 | 0.35 |
| K oxidation | mitochondrion | 0.02 | 4.17E-04 | 0.73 |
| K oxidation | plastid | 0.01 | -1.12E-04 | 0.8 |
| K oxidation | extracellular | NA | 0.00E+00 | NA |
| K oxidation | vacuole | 0.1 | -8.56E-03 | 0.44 |
| K oxidation | cytosol | 0 | -1.66E-05 | 0.99 |
| K oxidation | golgi | 0.01 | 1.87E-03 | 0.84 |
| K oxidation | nucleus | 0.32 | 2.35E-03 | 0.15 |
| K oxidation | PM | 0.18 | -9.03E-03 | 0.3 |
| K oxidation | ER | 0 | 2.33E-04 | 0.94 |
| K oxidation | peroxisome | 0.39 | 9.28E-03 | 0.1 |
| M dihydroxylation | mitochondrion | 0.97 | 4.44E-02 | 0 |
| M dihydroxylation | plastid | 0.69 | 1.43E-02 | 0.01 |
| M dihydroxylation | extracellular | NA | 0.00E+00 | NA |
| M dihydroxylation | vacuole | 0.74 | 2.86E-02 | 0.01 |
| M dihydroxylation | cytosol | 0.8 | 1.59E-02 | 0 |
| M dihydroxylation | golgi | 0.46 | 1.84E-02 | 0.07 |
| M dihydroxylation | nucleus | 0.78 | 1.56E-02 | 0 |
| M dihydroxylation | PM | 0.41 | 9.05E-03 | 0.09 |
| M dihydroxylation | ER | 0.6 | 1.23E-02 | 0.02 |
| M dihydroxylation | peroxisome | 0.04 | -2.27E-03 | 0.62 |
| M oxidation | mitochondrion | 0.97 | 6.48E-01 | 0 |
| M oxidation | plastid | 0.94 | 5.67E-01 | 0 |
| M oxidation | extracellular | 0.36 | 1.53E-01 | 0.12 |
| M oxidation | vacuole | 0.8 | 5.50E-01 | 0 |
| M oxidation | cytosol | 0.93 | 5.00E-01 | 0 |
| M oxidation | golgi | 0.86 | 6.12E-01 | 0 |
| M oxidation | nucleus | 0.95 | 4.86E-01 | 0 |
| M oxidation | PM | 0.45 | 1.60E-01 | 0.07 |
| M oxidation | ER | 0.74 | 2.88E-01 | 0.01 |
| M oxidation | peroxisome | 0.62 | 1.96E-01 | 0.02 |
| NTA | mitochondrion | 0 | -1.15E-04 | 0.98 |
| NTA | plastid | 0.11 | -8.36E-04 | 0.43 |
| NTA | extracellular | NA | 0.00E+00 | NA |
| NTA | vacuole | 0.7 | -3.58E-02 | 0.01 |
| NTA | cytosol | 0.02 | 1.95E-03 | 0.75 |
| NTA | golgi | 0.08 | 9.01E-03 | 0.49 |
| NTA | nucleus | 0.46 | -1.46E-02 | 0.06 |
| NTA | PM | 0.01 | -1.61E-03 | 0.77 |
| NTA | ER | NA | 0.00E+00 | NA |
| NTA | peroxisome | 0.42 | 2.81E-02 | 0.08 |
| P oxidation | mitochondrion | 0.35 | 1.19E-02 | 0.12 |
| P oxidation | plastid | 0.4 | 1.30E-02 | 0.09 |
| P oxidation | extracellular | 0.07 | 6.87E-03 | 0.53 |
| P oxidation | vacuole | 0.01 | -2.31E-03 | 0.8 |
| P oxidation | cytosol | 0.2 | 6.24E-03 | 0.27 |
| P oxidation | golgi | 0.19 | 2.43E-02 | 0.28 |
| P oxidation | nucleus | 0 | -3.03E-04 | 0.95 |
| P oxidation | PM | 0.45 | -4.02E-02 | 0.07 |
| P oxidation | ER | 0.28 | -3.56E-02 | 0.18 |
| P oxidation | peroxisome | 0.06 | -6.49E-03 | 0.57 |
| T oxidation | mitochondrion | 0.13 | 2.57E-03 | 0.37 |
| T oxidation | plastid | 0.23 | 2.88E-03 | 0.23 |
| T oxidation | extracellular | 0.37 | 1.92E-02 | 0.11 |
| T oxidation | vacuole | 0.04 | 1.50E-02 | 0.63 |
| T oxidation | cytosol | 0.04 | 2.57E-03 | 0.64 |
| T oxidation | golgi | 0.42 | -1.02E-02 | 0.08 |
| T oxidation | nucleus | 0.18 | 3.87E-03 | 0.29 |
| T oxidation | PM | 0.01 | -7.83E-04 | 0.77 |
| T oxidation | ER | 0.78 | -4.63E-02 | 0 |
| T oxidation | peroxisome | 0.12 | -1.60E-02 | 0.4 |
| W dihydroxylation | mitochondrion | 0.67 | 3.80E-03 | 0.01 |
| W dihydroxylation | plastid | 0.14 | 2.96E-04 | 0.36 |
| W dihydroxylation | extracellular | NA | 0.00E+00 | NA |
| W dihydroxylation | vacuole | 0.26 | -1.83E-02 | 0.2 |
| W dihydroxylation | cytosol | 0.34 | -5.18E-03 | 0.13 |
| W dihydroxylation | golgi | 0.05 | 2.28E-03 | 0.58 |
| W dihydroxylation | nucleus | 0.06 | 3.29E-04 | 0.55 |
| W dihydroxylation | PM | 0.07 | 2.90E-03 | 0.54 |
| W dihydroxylation | ER | 0.64 | -3.05E-02 | 0.02 |
| W dihydroxylation | peroxisome | NA | 0.00E+00 | NA |
| W oxidation | mitochondrion | 0.14 | 3.00E-03 | 0.36 |
| W oxidation | plastid | 0.03 | -2.63E-04 | 0.7 |
| W oxidation | extracellular | NA | 0.00E+00 | NA |
| W oxidation | vacuole | 0.28 | -2.05E-02 | 0.18 |
| W oxidation | cytosol | 0.27 | -1.12E-02 | 0.19 |
| W oxidation | golgi | 0.02 | -3.14E-03 | 0.74 |
| W oxidation | nucleus | 0.21 | 9.36E-04 | 0.25 |
| W oxidation | PM | 0.33 | -2.27E-02 | 0.14 |
| W oxidation | ER | 0.78 | -1.25E-01 | 0 |
| W oxidation | peroxisome | 0.03 | -1.46E-03 | 0.69 |
| pyro-Glu | mitochondrion | NA | 0.00E+00 | NA |
| pyro-Glu | plastid | NA | 0.00E+00 | NA |
| pyro-Glu | extracellular | NA | 0.00E+00 | NA |
| pyro-Glu | vacuole | NA | 0.00E+00 | NA |
| pyro-Glu | cytosol | NA | 0.00E+00 | NA |
| pyro-Glu | golgi | NA | 0.00E+00 | NA |
| pyro-Glu | nucleus | NA | 0.00E+00 | NA |
| pyro-Glu | PM | NA | 0.00E+00 | NA |
| pyro-Glu | ER | NA | 0.00E+00 | NA |
| pyro-Glu | peroxisome | NA | 0.00E+00 | NA |

Supplemental Table S5. Correlations between PTM frequencies and time for proteins in different MapMan functional categories.

| PTM | Functional category | R^2^ | Slope | p |
| --- | --- | --- | --- | --- |
| C dihydroxylation | Primary metabolism | 0.04 | 9.62E-04 | 0.62 |
| C dihydroxylation | Cellular organisation | 0.04 | 1.91E-03 | 0.64 |
| C dihydroxylation | Other | 0.23 | 1.19E-03 | 0.23 |
| C dihydroxylation | Translation | 0.07 | 3.70E-03 | 0.53 |
| C dihydroxylation | RNA biosynthesis & processing | 0.76 | 3.24E-02 | 0 |
| C dihydroxylation | Ribosomal subunits | 0.1 | -1.19E-03 | 0.44 |
| C dihydroxylation | Protein biosynthesis & homeostasis | 0.02 | 5.30E-04 | 0.74 |
| C dihydroxylation | Protein modification & translocation | 0.5 | 9.20E-03 | 0.05 |
| C dihydroxylation | Proteosome & proteolysis | NA | 0.00E+00 | NA |
| C dihydroxylation | Secondary metabolism | NA | 0.00E+00 | NA |
| C dihydroxylation | NA | 0.6 | -2.29E-03 | 0.02 |
| C oxidation | Primary metabolism | 0.62 | 1.20E-01 | 0.02 |
| C oxidation | Cellular organisation | 0.29 | 8.11E-02 | 0.17 |
| C oxidation | Other | 0.78 | 1.19E-01 | 0 |
| C oxidation | Translation | 0.02 | 7.34E-03 | 0.77 |
| C oxidation | RNA biosynthesis & processing | 0.93 | 1.66E-01 | 0 |
| C oxidation | Ribosomal subunits | 0.12 | 4.62E-02 | 0.4 |
| C oxidation | Protein biosynthesis & homeostasis | 0.38 | 7.27E-02 | 0.1 |
| C oxidation | Protein modification & translocation | 0.67 | 1.02E-01 | 0.01 |
| C oxidation | Proteosome & proteolysis | 0.36 | 6.50E-02 | 0.12 |
| C oxidation | Secondary metabolism | 0.01 | 1.16E-02 | 0.81 |
| C oxidation | NA | 0.26 | 4.77E-02 | 0.2 |
| K methylation | Primary metabolism | 0.17 | -5.48E-03 | 0.31 |
| K methylation | Cellular organisation | 0.04 | -1.50E-03 | 0.62 |
| K methylation | Other | 0.5 | -9.01E-03 | 0.05 |
| K methylation | Translation | 0.21 | -7.75E-03 | 0.26 |
| K methylation | RNA biosynthesis & processing | 0.09 | 3.55E-03 | 0.46 |
| K methylation | Ribosomal subunits | 0.38 | -2.25E-02 | 0.1 |
| K methylation | Protein biosynthesis & homeostasis | 0.76 | -6.00E-02 | 0 |
| K methylation | Protein modification & translocation | 0.88 | -3.32E-02 | 0 |
| K methylation | Proteosome & proteolysis | 0.67 | -1.54E-02 | 0.01 |
| K methylation | Secondary metabolism | NA | 0.00E+00 | NA |
| K methylation | NA | 0.81 | -2.28E-02 | 0 |
| K oxidation | Primary metabolism | 0 | 1.40E-04 | 0.88 |
| K oxidation | Cellular organisation | 0.03 | -9.19E-04 | 0.69 |
| K oxidation | Other | 0.43 | 2.95E-03 | 0.08 |
| K oxidation | Translation | 0 | 2.81E-04 | 0.88 |
| K oxidation | RNA biosynthesis & processing | 0.28 | 4.12E-03 | 0.18 |
| K oxidation | Ribosomal subunits | 0.27 | 2.83E-03 | 0.19 |
| K oxidation | Protein biosynthesis & homeostasis | 0.08 | -4.69E-03 | 0.51 |
| K oxidation | Protein modification & translocation | 0.13 | -3.00E-03 | 0.38 |
| K oxidation | Proteosome & proteolysis | NA | 0.00E+00 | NA |
| K oxidation | Secondary metabolism | 0.14 | 1.30E-02 | 0.36 |
| K oxidation | NA | 0 | 6.81E-05 | 0.97 |
| M dihydroxylation | Primary metabolism | 0.74 | 2.69E-02 | 0.01 |
| M dihydroxylation | Cellular organisation | 0.17 | 1.25E-02 | 0.31 |
| M dihydroxylation | Other | 0.63 | 1.00E-02 | 0.02 |
| M dihydroxylation | Translation | 0.94 | 2.33E-02 | 0 |
| M dihydroxylation | RNA biosynthesis & processing | 0.19 | 5.04E-03 | 0.28 |
| M dihydroxylation | Ribosomal subunits | 0.44 | 2.11E-02 | 0.07 |
| M dihydroxylation | Protein biosynthesis & homeostasis | 0.84 | 6.10E-02 | 0 |
| M dihydroxylation | Protein modification & translocation | 0.88 | 6.13E-02 | 0 |
| M dihydroxylation | Proteosome & proteolysis | 0.43 | 1.10E-02 | 0.08 |
| M dihydroxylation | Secondary metabolism | NA | 0.00E+00 | NA |
| M dihydroxylation | NA | 0.93 | 1.40E-02 | 0 |
| M oxidation | Primary metabolism | 0.95 | 5.78E-01 | 0 |
| M oxidation | Cellular organisation | 0.94 | 5.67E-01 | 0 |
| M oxidation | Other | 0.97 | 5.91E-01 | 0 |
| M oxidation | Translation | 0.88 | 5.89E-01 | 0 |
| M oxidation | RNA biosynthesis & processing | 0.95 | 5.61E-01 | 0 |
| M oxidation | Ribosomal subunits | 0.9 | 4.49E-01 | 0 |
| M oxidation | Protein biosynthesis & homeostasis | 0.87 | 4.46E-01 | 0 |
| M oxidation | Protein modification & translocation | 0.98 | 5.28E-01 | 0 |
| M oxidation | Proteosome & proteolysis | 0.97 | 5.30E-01 | 0 |
| M oxidation | Secondary metabolism | 0.14 | 1.58E-01 | 0.36 |
| M oxidation | NA | 0.94 | 5.37E-01 | 0 |
| NTA | Primary metabolism | 0.04 | -3.11E-03 | 0.63 |
| NTA | Cellular organisation | 0.45 | -2.73E-02 | 0.07 |
| NTA | Other | 0.21 | -8.48E-03 | 0.26 |
| NTA | Translation | 0.01 | -3.41E-03 | 0.84 |
| NTA | RNA biosynthesis & processing | 0.39 | -1.84E-02 | 0.1 |
| NTA | Ribosomal subunits | 0.33 | 2.64E-02 | 0.13 |
| NTA | Protein biosynthesis & homeostasis | 0.9 | -2.41E-02 | 0 |
| NTA | Protein modification & translocation | 0.81 | 4.88E-02 | 0 |
| NTA | Proteosome & proteolysis | 0.01 | -3.51E-03 | 0.79 |
| NTA | Secondary metabolism | NA | 0.00E+00 | NA |
| NTA | NA | 0.87 | -2.07E-02 | 0 |
| P oxidation | Primary metabolism | 0.07 | 7.92E-03 | 0.52 |
| P oxidation | Cellular organisation | 0 | -4.50E-04 | 0.97 |
| P oxidation | Other | 0.78 | 2.61E-02 | 0 |
| P oxidation | Translation | 0.52 | 1.29E-02 | 0.04 |
| P oxidation | RNA biosynthesis & processing | 0.19 | -4.51E-03 | 0.29 |
| P oxidation | Ribosomal subunits | 0.13 | 1.01E-02 | 0.38 |
| P oxidation | Protein biosynthesis & homeostasis | 0.35 | -1.38E-02 | 0.12 |
| P oxidation | Protein modification & translocation | 0.28 | 2.37E-02 | 0.18 |
| P oxidation | Proteosome & proteolysis | 0.09 | 7.27E-03 | 0.47 |
| P oxidation | Secondary metabolism | NA | 0.00E+00 | NA |
| P oxidation | NA | 0.78 | -1.11E-02 | 0 |
| T oxidation | Primary metabolism | 0.65 | 3.39E-03 | 0.02 |
| T oxidation | Cellular organisation | 0.05 | -2.07E-03 | 0.6 |
| T oxidation | Other | 0.85 | 8.67E-03 | 0 |
| T oxidation | Translation | 0.06 | -4.27E-03 | 0.56 |
| T oxidation | RNA biosynthesis & processing | 0.37 | 1.81E-02 | 0.11 |
| T oxidation | Ribosomal subunits | 0.02 | 4.40E-03 | 0.71 |
| T oxidation | Protein biosynthesis & homeostasis | 0.56 | -3.93E-02 | 0.03 |
| T oxidation | Protein modification & translocation | 0.03 | 3.59E-03 | 0.67 |
| T oxidation | Proteosome & proteolysis | 0.77 | 1.90E-02 | 0 |
| T oxidation | Secondary metabolism | 0.42 | -5.67E-02 | 0.08 |
| T oxidation | NA | 0.08 | -3.94E-03 | 0.5 |
| W dihydroxylation | Primary metabolism | 0.68 | 1.93E-03 | 0.01 |
| W dihydroxylation | Cellular organisation | 0.04 | 7.56E-04 | 0.62 |
| W dihydroxylation | Other | 0 | 1.30E-04 | 0.88 |
| W dihydroxylation | Translation | 0.06 | 4.66E-04 | 0.57 |
| W dihydroxylation | RNA biosynthesis & processing | 0.82 | 3.58E-03 | 0 |
| W dihydroxylation | Ribosomal subunits | 0.01 | 1.18E-03 | 0.8 |
| W dihydroxylation | Protein biosynthesis & homeostasis | 0.11 | -8.36E-03 | 0.42 |
| W dihydroxylation | Protein modification & translocation | 0.1 | -2.72E-03 | 0.45 |
| W dihydroxylation | Proteosome & proteolysis | 0 | 8.58E-05 | 0.95 |
| W dihydroxylation | Secondary metabolism | NA | 0.00E+00 | NA |
| W dihydroxylation | NA | 0.66 | -7.58E-03 | 0.01 |
| W oxidation | Primary metabolism | 0 | -2.73E-04 | 0.91 |
| W oxidation | Cellular organisation | 0.23 | 4.62E-03 | 0.22 |
| W oxidation | Other | 0.1 | -3.31E-03 | 0.46 |
| W oxidation | Translation | 0.66 | 4.55E-03 | 0.01 |
| W oxidation | RNA biosynthesis & processing | 0.82 | 3.58E-03 | 0 |
| W oxidation | Ribosomal subunits | 0.01 | -5.41E-04 | 0.78 |
| W oxidation | Protein biosynthesis & homeostasis | 0.17 | -1.79E-02 | 0.3 |
| W oxidation | Protein modification & translocation | 0.21 | 1.29E-03 | 0.26 |
| W oxidation | Proteosome & proteolysis | 0.03 | -1.94E-03 | 0.67 |
| W oxidation | Secondary metabolism | NA | 0.00E+00 | NA |
| W oxidation | NA | 0.77 | -3.43E-02 | 0 |
| pyro-Glu | Primary metabolism | NA | 0.00E+00 | NA |
| pyro-Glu | Cellular organisation | NA | 0.00E+00 | NA |
| pyro-Glu | Other | NA | 0.00E+00 | NA |
| pyro-Glu | Translation | NA | 0.00E+00 | NA |
| pyro-Glu | RNA biosynthesis & processing | NA | 0.00E+00 | NA |
| pyro-Glu | Ribosomal subunits | NA | 0.00E+00 | NA |
| pyro-Glu | Protein biosynthesis & homeostasis | NA | 0.00E+00 | NA |
| pyro-Glu | Protein modification & translocation | NA | 0.00E+00 | NA |
| pyro-Glu | Proteosome & proteolysis | NA | 0.00E+00 | NA |
| pyro-Glu | Secondary metabolism | NA | 0.00E+00 | NA |
| pyro-Glu | NA | NA | 0.00E+00 | NA |

Supplemental Table S6. Correlations between PTM frequencies and time for proteins with different Pfam domains.

| PTM | Pfam ID | R^2^ | Slope | p |
| --- | --- | --- | --- | --- |
| C dihydroxylation | PF00004 | NA | 0.00E+00 | NA |
| C dihydroxylation | PF00012 | 0.2 | -3.42E-03 | 0.27 |
| C dihydroxylation | PF00069 | NA | 0.00E+00 | NA |
| C dihydroxylation | PF00071 | NA | 0.00E+00 | NA |
| C dihydroxylation | PF00076 | NA | 0.00E+00 | NA |
| C dihydroxylation | PF00085 | NA | 0.00E+00 | NA |
| C dihydroxylation | PF00118 | 0.22 | -2.30E-03 | 0.24 |
| C dihydroxylation | PF00160 | 0.21 | 4.07E-02 | 0.25 |
| C dihydroxylation | PF00179 | NA | 0.00E+00 | NA |
| C dihydroxylation | PF00226 | 0.2 | 8.81E-03 | 0.26 |
| C dihydroxylation | PF00270 | 0.67 | 5.78E-02 | 0.01 |
| C dihydroxylation | PF00271 | 0.68 | 5.85E-02 | 0.01 |
| C dihydroxylation | PF00400 | NA | 0.00E+00 | NA |
| C dihydroxylation | PF01535 | NA | 0.00E+00 | NA |
| C dihydroxylation | PF08240 | NA | 0.00E+00 | NA |
| C dihydroxylation | PF13041 | NA | 0.00E+00 | NA |
| C dihydroxylation | PF17862 | NA | 0.00E+00 | NA |
| C oxidation | PF00004 | 0.17 | 5.52E-02 | 0.3 |
| C oxidation | PF00012 | 0.15 | -4.12E-02 | 0.34 |
| C oxidation | PF00069 | 0 | 4.55E-03 | 0.94 |
| C oxidation | PF00071 | 0.29 | -1.66E-01 | 0.16 |
| C oxidation | PF00076 | 0.06 | 2.76E-02 | 0.55 |
| C oxidation | PF00085 | 0.09 | 1.43E-02 | 0.48 |
| C oxidation | PF00118 | 0.26 | 1.10E-01 | 0.2 |
| C oxidation | PF00160 | 0.51 | 3.43E-01 | 0.05 |
| C oxidation | PF00179 | 0.08 | 5.23E-02 | 0.5 |
| C oxidation | PF00226 | 0.1 | 8.17E-02 | 0.45 |
| C oxidation | PF00270 | 0.79 | 1.88E-01 | 0 |
| C oxidation | PF00271 | 0.79 | 1.86E-01 | 0 |
| C oxidation | PF00400 | 0.22 | 6.89E-02 | 0.25 |
| C oxidation | PF01535 | 0.59 | 1.21E-01 | 0.03 |
| C oxidation | PF08240 | 0.36 | 7.51E-02 | 0.12 |
| C oxidation | PF13041 | 0.41 | 1.09E-01 | 0.09 |
| C oxidation | PF17862 | 0.34 | 1.08E-01 | 0.13 |
| K methylation | PF00004 | 0.65 | -3.01E-02 | 0.02 |
| K methylation | PF00012 | 0.39 | -5.70E-02 | 0.1 |
| K methylation | PF00069 | NA | 0.00E+00 | NA |
| K methylation | PF00071 | NA | 0.00E+00 | NA |
| K methylation | PF00076 | 0.32 | -2.20E-02 | 0.15 |
| K methylation | PF00085 | 0 | -1.53E-03 | 0.93 |
| K methylation | PF00118 | 0.58 | -2.76E-02 | 0.03 |
| K methylation | PF00160 | 0.04 | -1.22E-02 | 0.62 |
| K methylation | PF00179 | NA | 0.00E+00 | NA |
| K methylation | PF00226 | NA | 0.00E+00 | NA |
| K methylation | PF00270 | 0 | 3.60E-04 | 0.88 |
| K methylation | PF00271 | 0 | 3.51E-04 | 0.88 |
| K methylation | PF00400 | NA | 0.00E+00 | NA |
| K methylation | PF01535 | NA | 0.00E+00 | NA |
| K methylation | PF08240 | NA | 0.00E+00 | NA |
| K methylation | PF13041 | NA | 0.00E+00 | NA |
| K methylation | PF17862 | 0.27 | -1.94E-02 | 0.19 |
| K oxidation | PF00004 | 0.2 | 9.72E-04 | 0.26 |
| K oxidation | PF00012 | 0.23 | -1.39E-02 | 0.23 |
| K oxidation | PF00069 | 0.16 | -2.93E-02 | 0.32 |
| K oxidation | PF00071 | NA | 0.00E+00 | NA |
| K oxidation | PF00076 | 0.21 | 5.19E-03 | 0.25 |
| K oxidation | PF00085 | NA | 0.00E+00 | NA |
| K oxidation | PF00118 | 0.03 | 2.13E-03 | 0.69 |
| K oxidation | PF00160 | 0.06 | 4.02E-03 | 0.57 |
| K oxidation | PF00179 | NA | 0.00E+00 | NA |
| K oxidation | PF00226 | NA | 0.00E+00 | NA |
| K oxidation | PF00270 | NA | 0.00E+00 | NA |
| K oxidation | PF00271 | NA | 0.00E+00 | NA |
| K oxidation | PF00400 | NA | 0.00E+00 | NA |
| K oxidation | PF01535 | NA | 0.00E+00 | NA |
| K oxidation | PF08240 | NA | 0.00E+00 | NA |
| K oxidation | PF13041 | NA | 0.00E+00 | NA |
| K oxidation | PF17862 | NA | 0.00E+00 | NA |
| M dihydroxylation | PF00004 | 0.37 | 7.37E-03 | 0.11 |
| M dihydroxylation | PF00012 | 0 | -2.05E-03 | 0.9 |
| M dihydroxylation | PF00069 | 0.2 | 9.42E-03 | 0.26 |
| M dihydroxylation | PF00071 | NA | 0.00E+00 | NA |
| M dihydroxylation | PF00076 | 0.16 | 5.52E-03 | 0.32 |
| M dihydroxylation | PF00085 | 0 | 8.77E-04 | 0.88 |
| M dihydroxylation | PF00118 | 0.83 | 5.59E-02 | 0 |
| M dihydroxylation | PF00160 | 0.3 | 4.93E-02 | 0.16 |
| M dihydroxylation | PF00179 | 0.17 | -4.83E-02 | 0.31 |
| M dihydroxylation | PF00226 | 0 | 1.23E-03 | 0.88 |
| M dihydroxylation | PF00270 | 0.03 | 1.98E-03 | 0.69 |
| M dihydroxylation | PF00271 | 0.03 | 2.04E-03 | 0.67 |
| M dihydroxylation | PF00400 | 0.46 | 1.10E-02 | 0.06 |
| M dihydroxylation | PF01535 | NA | 0.00E+00 | NA |
| M dihydroxylation | PF08240 | 0.04 | 7.07E-03 | 0.62 |
| M dihydroxylation | PF13041 | NA | 0.00E+00 | NA |
| M dihydroxylation | PF17862 | 0.33 | 9.81E-03 | 0.14 |
| M oxidation | PF00004 | 0.88 | 4.95E-01 | 0 |
| M oxidation | PF00012 | 0.84 | 5.11E-01 | 0 |
| M oxidation | PF00069 | 0.13 | 9.31E-02 | 0.39 |
| M oxidation | PF00071 | 0.86 | 8.21E-01 | 0 |
| M oxidation | PF00076 | 0.96 | 5.28E-01 | 0 |
| M oxidation | PF00085 | 0.04 | 4.25E-02 | 0.63 |
| M oxidation | PF00118 | 0.89 | 4.90E-01 | 0 |
| M oxidation | PF00160 | 0.69 | 4.97E-01 | 0.01 |
| M oxidation | PF00179 | 0.42 | 5.55E-01 | 0.08 |
| M oxidation | PF00226 | 0.91 | 6.52E-01 | 0 |
| M oxidation | PF00270 | 0.91 | 5.63E-01 | 0 |
| M oxidation | PF00271 | 0.9 | 5.58E-01 | 0 |
| M oxidation | PF00400 | 0.38 | 1.70E-01 | 0.1 |
| M oxidation | PF01535 | 0.73 | 1.02E+00 | 0.01 |
| M oxidation | PF08240 | 0.78 | 6.01E-01 | 0 |
| M oxidation | PF13041 | 0.74 | 1.07E+00 | 0.01 |
| M oxidation | PF17862 | 0.93 | 6.30E-01 | 0 |
| NTA | PF00004 | 0.13 | -9.74E-03 | 0.38 |
| NTA | PF00012 | 0.8 | -1.60E-02 | 0 |
| NTA | PF00069 | NA | 0.00E+00 | NA |
| NTA | PF00071 | NA | 0.00E+00 | NA |
| NTA | PF00076 | 0.24 | -2.09E-02 | 0.22 |
| NTA | PF00085 | 0.34 | -7.42E-02 | 0.13 |
| NTA | PF00118 | 0.02 | -2.03E-03 | 0.71 |
| NTA | PF00160 | 0.14 | -3.72E-02 | 0.37 |
| NTA | PF00179 | 0.17 | -7.27E-02 | 0.31 |
| NTA | PF00226 | NA | 0.00E+00 | NA |
| NTA | PF00270 | 0.55 | -1.99E-02 | 0.03 |
| NTA | PF00271 | 0.55 | -1.94E-02 | 0.04 |
| NTA | PF00400 | 0.04 | -9.12E-03 | 0.64 |
| NTA | PF01535 | NA | 0.00E+00 | NA |
| NTA | PF08240 | NA | 0.00E+00 | NA |
| NTA | PF13041 | NA | 0.00E+00 | NA |
| NTA | PF17862 | 0.04 | -9.76E-03 | 0.62 |
| P oxidation | PF00004 | 0.18 | -2.02E-02 | 0.29 |
| P oxidation | PF00012 | 0.81 | -5.16E-02 | 0 |
| P oxidation | PF00069 | 0.28 | -2.05E-02 | 0.18 |
| P oxidation | PF00071 | NA | 0.00E+00 | NA |
| P oxidation | PF00076 | 0.01 | -1.68E-03 | 0.82 |
| P oxidation | PF00085 | 0.04 | -1.09E-02 | 0.64 |
| P oxidation | PF00118 | 0.1 | -4.82E-03 | 0.44 |
| P oxidation | PF00160 | 0 | 3.41E-05 | 1 |
| P oxidation | PF00179 | NA | 0.00E+00 | NA |
| P oxidation | PF00226 | NA | 0.00E+00 | NA |
| P oxidation | PF00270 | 0.13 | -9.10E-03 | 0.38 |
| P oxidation | PF00271 | 0.13 | -8.85E-03 | 0.38 |
| P oxidation | PF00400 | 0.01 | -2.03E-03 | 0.86 |
| P oxidation | PF01535 | NA | 0.00E+00 | NA |
| P oxidation | PF08240 | 0.5 | -6.16E-02 | 0.05 |
| P oxidation | PF13041 | 0.06 | 4.66E-03 | 0.57 |
| P oxidation | PF17862 | 0.02 | 8.37E-03 | 0.73 |
| T oxidation | PF00004 | 0 | -2.45E-04 | 0.97 |
| T oxidation | PF00012 | 0.34 | -4.51E-02 | 0.13 |
| T oxidation | PF00069 | NA | 0.00E+00 | NA |
| T oxidation | PF00071 | NA | 0.00E+00 | NA |
| T oxidation | PF00076 | 0 | -1.18E-03 | 0.92 |
| T oxidation | PF00085 | 0.03 | -2.35E-03 | 0.69 |
| T oxidation | PF00118 | 0.21 | -1.78E-02 | 0.25 |
| T oxidation | PF00160 | 0.05 | 1.41E-02 | 0.61 |
| T oxidation | PF00179 | 0.66 | 1.52E-01 | 0.01 |
| T oxidation | PF00226 | 0.03 | -1.12E-02 | 0.67 |
| T oxidation | PF00270 | 0.35 | 1.92E-02 | 0.12 |
| T oxidation | PF00271 | 0.36 | 1.90E-02 | 0.11 |
| T oxidation | PF00400 | 0.03 | -5.27E-03 | 0.69 |
| T oxidation | PF01535 | NA | 0.00E+00 | NA |
| T oxidation | PF08240 | NA | 0.00E+00 | NA |
| T oxidation | PF13041 | NA | 0.00E+00 | NA |
| T oxidation | PF17862 | 0.49 | 1.51E-02 | 0.05 |
| W dihydroxylation | PF00004 | 0 | 1.18E-04 | 0.95 |
| W dihydroxylation | PF00012 | 0.63 | -4.29E-02 | 0.02 |
| W dihydroxylation | PF00069 | NA | 0.00E+00 | NA |
| W dihydroxylation | PF00071 | 0.13 | 3.76E-02 | 0.38 |
| W dihydroxylation | PF00076 | 0.82 | 7.45E-03 | 0 |
| W dihydroxylation | PF00085 | NA | 0.00E+00 | NA |
| W dihydroxylation | PF00118 | 0.37 | 1.22E-02 | 0.11 |
| W dihydroxylation | PF00160 | NA | 0.00E+00 | NA |
| W dihydroxylation | PF00179 | NA | 0.00E+00 | NA |
| W dihydroxylation | PF00226 | NA | 0.00E+00 | NA |
| W dihydroxylation | PF00270 | NA | 0.00E+00 | NA |
| W dihydroxylation | PF00271 | NA | 0.00E+00 | NA |
| W dihydroxylation | PF00400 | 0.04 | 2.76E-03 | 0.63 |
| W dihydroxylation | PF01535 | NA | 0.00E+00 | NA |
| W dihydroxylation | PF08240 | NA | 0.00E+00 | NA |
| W dihydroxylation | PF13041 | NA | 0.00E+00 | NA |
| W dihydroxylation | PF17862 | 0 | 3.22E-04 | 0.93 |
| W oxidation | PF00004 | 0.05 | -3.38E-03 | 0.61 |
| W oxidation | PF00012 | 0.68 | -8.25E-02 | 0.01 |
| W oxidation | PF00069 | NA | 0.00E+00 | NA |
| W oxidation | PF00071 | 0 | -1.23E-02 | 0.91 |
| W oxidation | PF00076 | 0.82 | 7.45E-03 | 0 |
| W oxidation | PF00085 | NA | 0.00E+00 | NA |
| W oxidation | PF00118 | 0.15 | 1.09E-02 | 0.35 |
| W oxidation | PF00160 | NA | 0.00E+00 | NA |
| W oxidation | PF00179 | NA | 0.00E+00 | NA |
| W oxidation | PF00226 | NA | 0.00E+00 | NA |
| W oxidation | PF00270 | NA | 0.00E+00 | NA |
| W oxidation | PF00271 | NA | 0.00E+00 | NA |
| W oxidation | PF00400 | 0.2 | 9.22E-03 | 0.27 |
| W oxidation | PF01535 | NA | 0.00E+00 | NA |
| W oxidation | PF08240 | NA | 0.00E+00 | NA |
| W oxidation | PF13041 | NA | 0.00E+00 | NA |
| W oxidation | PF17862 | 0.03 | -5.20E-03 | 0.69 |
| pyro-Glu | PF00004 | NA | 0.00E+00 | NA |
| pyro-Glu | PF00012 | NA | 0.00E+00 | NA |
| pyro-Glu | PF00069 | NA | 0.00E+00 | NA |
| pyro-Glu | PF00071 | NA | 0.00E+00 | NA |
| pyro-Glu | PF00076 | NA | 0.00E+00 | NA |
| pyro-Glu | PF00085 | NA | 0.00E+00 | NA |
| pyro-Glu | PF00118 | NA | 0.00E+00 | NA |
| pyro-Glu | PF00160 | NA | 0.00E+00 | NA |
| pyro-Glu | PF00179 | NA | 0.00E+00 | NA |
| pyro-Glu | PF00226 | NA | 0.00E+00 | NA |
| pyro-Glu | PF00270 | NA | 0.00E+00 | NA |
| pyro-Glu | PF00271 | NA | 0.00E+00 | NA |
| pyro-Glu | PF00400 | NA | 0.00E+00 | NA |
| pyro-Glu | PF01535 | NA | 0.00E+00 | NA |
| pyro-Glu | PF08240 | NA | 0.00E+00 | NA |
| pyro-Glu | PF13041 | NA | 0.00E+00 | NA |
| pyro-Glu | PF17862 | NA | 0.00E+00 | NA |

Table S7. Correlations between PTM frequencies and time for proteins with different Interpro domains.

| PTM | Interpro domai | R^2^ | Slope | p |
| --- | --- | --- | --- | --- |
| C dihydroxylation | IPR000504 | NA | 0.00E+00 | NA |
| C dihydroxylation | IPR000719 | NA | 0.00E+00 | NA |
| C dihydroxylation | IPR001650 | 0.68 | 5.85E-02 | 0.01 |
| C dihydroxylation | IPR001680 | NA | 0.00E+00 | NA |
| C dihydroxylation | IPR002885 | NA | 0.00E+00 | NA |
| C dihydroxylation | IPR003593 | 0.31 | 3.77E-03 | 0.15 |
| C dihydroxylation | IPR003959 | NA | 0.00E+00 | NA |
| C dihydroxylation | IPR003960 | NA | 0.00E+00 | NA |
| C dihydroxylation | IPR005225 | NA | 0.00E+00 | NA |
| C dihydroxylation | IPR008271 | NA | 0.00E+00 | NA |
| C dihydroxylation | IPR011009 | NA | 0.00E+00 | NA |
| C dihydroxylation | IPR011545 | 0.67 | 5.78E-02 | 0.01 |
| C dihydroxylation | IPR011989 | NA | 0.00E+00 | NA |
| C dihydroxylation | IPR011990 | 0 | 8.87E-05 | 0.98 |
| C dihydroxylation | IPR012340 | NA | 0.00E+00 | NA |
| C dihydroxylation | IPR012677 | NA | 0.00E+00 | NA |
| C dihydroxylation | IPR013766 | NA | 0.00E+00 | NA |
| C dihydroxylation | IPR013785 | 0.13 | 7.00E-03 | 0.38 |
| C dihydroxylation | IPR014001 | 0.67 | 5.36E-02 | 0.01 |
| C dihydroxylation | IPR014014 | 0.66 | 6.78E-02 | 0.01 |
| C dihydroxylation | IPR014729 | 0.06 | 1.42E-03 | 0.57 |
| C dihydroxylation | IPR015943 | NA | 0.00E+00 | NA |
| C dihydroxylation | IPR016024 | NA | 0.00E+00 | NA |
| C dihydroxylation | IPR017441 | NA | 0.00E+00 | NA |
| C dihydroxylation | IPR019734 | 0.01 | 9.18E-04 | 0.86 |
| C dihydroxylation | IPR027417 | 0.6 | 1.07E-02 | 0.02 |
| C dihydroxylation | IPR029063 | NA | 0.00E+00 | NA |
| C dihydroxylation | IPR035979 | NA | 0.00E+00 | NA |
| C dihydroxylation | IPR036249 | NA | 0.00E+00 | NA |
| C dihydroxylation | IPR036291 | 0 | -5.29E-04 | 0.95 |
| C dihydroxylation | IPR036322 | NA | 0.00E+00 | NA |
| C oxidation | IPR000504 | 0.06 | 2.76E-02 | 0.55 |
| C oxidation | IPR000719 | 0 | 7.46E-03 | 0.89 |
| C oxidation | IPR001650 | 0.79 | 1.86E-01 | 0 |
| C oxidation | IPR001680 | 0.19 | 5.05E-02 | 0.28 |
| C oxidation | IPR002885 | 0.65 | 1.14E-01 | 0.02 |
| C oxidation | IPR003593 | 0.25 | 4.88E-02 | 0.21 |
| C oxidation | IPR003959 | 0.17 | 5.52E-02 | 0.3 |
| C oxidation | IPR003960 | 0.31 | 9.29E-02 | 0.15 |
| C oxidation | IPR005225 | 0.6 | -8.01E-02 | 0.02 |
| C oxidation | IPR008271 | 0 | 4.55E-03 | 0.94 |
| C oxidation | IPR011009 | 0.02 | 1.73E-02 | 0.75 |
| C oxidation | IPR011545 | 0.79 | 1.88E-01 | 0 |
| C oxidation | IPR011989 | 0.15 | 2.51E-02 | 0.34 |
| C oxidation | IPR011990 | 0.08 | -2.41E-02 | 0.51 |
| C oxidation | IPR012340 | 0.62 | 9.66E-02 | 0.02 |
| C oxidation | IPR012677 | 0.06 | 2.75E-02 | 0.55 |
| C oxidation | IPR013766 | 0.1 | 9.06E-03 | 0.44 |
| C oxidation | IPR013785 | 0.58 | 1.50E-01 | 0.03 |
| C oxidation | IPR014001 | 0.77 | 1.71E-01 | 0 |
| C oxidation | IPR014014 | 0.73 | 1.87E-01 | 0.01 |
| C oxidation | IPR014729 | 0.01 | 2.01E-02 | 0.82 |
| C oxidation | IPR015943 | 0.24 | 4.90E-02 | 0.22 |
| C oxidation | IPR016024 | 0.05 | 1.58E-02 | 0.58 |
| C oxidation | IPR017441 | 0.01 | 1.27E-02 | 0.83 |
| C oxidation | IPR019734 | 0.07 | -2.52E-02 | 0.54 |
| C oxidation | IPR027417 | 0.39 | 5.37E-02 | 0.1 |
| C oxidation | IPR029063 | 0.14 | 3.52E-02 | 0.35 |
| C oxidation | IPR035979 | 0.06 | 2.76E-02 | 0.55 |
| C oxidation | IPR036249 | 0 | 7.03E-04 | 0.93 |
| C oxidation | IPR036291 | 0.48 | 7.20E-02 | 0.06 |
| C oxidation | IPR036322 | 0.1 | 2.97E-02 | 0.45 |
| K methylation | IPR000504 | 0.32 | -2.20E-02 | 0.15 |
| K methylation | IPR000719 | NA | 0.00E+00 | NA |
| K methylation | IPR001650 | 0 | 3.51E-04 | 0.88 |
| K methylation | IPR001680 | NA | 0.00E+00 | NA |
| K methylation | IPR002885 | NA | 0.00E+00 | NA |
| K methylation | IPR003593 | 0.16 | -1.08E-02 | 0.32 |
| K methylation | IPR003959 | 0.65 | -3.01E-02 | 0.02 |
| K methylation | IPR003960 | 0.29 | -1.87E-02 | 0.17 |
| K methylation | IPR005225 | 0 | -1.38E-03 | 0.92 |
| K methylation | IPR008271 | NA | 0.00E+00 | NA |
| K methylation | IPR011009 | NA | 0.00E+00 | NA |
| K methylation | IPR011545 | 0 | 3.60E-04 | 0.88 |
| K methylation | IPR011989 | 0.03 | -1.23E-03 | 0.69 |
| K methylation | IPR011990 | 0.61 | -2.67E-02 | 0.02 |
| K methylation | IPR012340 | 0.06 | 3.70E-03 | 0.57 |
| K methylation | IPR012677 | 0.35 | -2.32E-02 | 0.13 |
| K methylation | IPR013766 | 0 | -1.58E-03 | 0.91 |
| K methylation | IPR013785 | 0.73 | -2.80E-02 | 0.01 |
| K methylation | IPR014001 | 0 | 3.20E-04 | 0.88 |
| K methylation | IPR014014 | 0 | 4.21E-04 | 0.88 |
| K methylation | IPR014729 | 0.29 | 1.53E-02 | 0.17 |
| K methylation | IPR015943 | NA | 0.00E+00 | NA |
| K methylation | IPR016024 | 0.03 | -1.02E-03 | 0.69 |
| K methylation | IPR017441 | NA | 0.00E+00 | NA |
| K methylation | IPR019734 | 0.42 | -3.24E-02 | 0.08 |
| K methylation | IPR027417 | 0.06 | -3.53E-03 | 0.55 |
| K methylation | IPR029063 | NA | 0.00E+00 | NA |
| K methylation | IPR035979 | 0.32 | -2.20E-02 | 0.15 |
| K methylation | IPR036249 | 0.7 | -2.73E-02 | 0.01 |
| K methylation | IPR036291 | 0.9 | -4.02E-02 | 0 |
| K methylation | IPR036322 | NA | 0.00E+00 | NA |
| K oxidation | IPR000504 | 0.21 | 5.19E-03 | 0.25 |
| K oxidation | IPR000719 | 0.16 | -2.44E-02 | 0.32 |
| K oxidation | IPR001650 | NA | 0.00E+00 | NA |
| K oxidation | IPR001680 | NA | 0.00E+00 | NA |
| K oxidation | IPR002885 | NA | 0.00E+00 | NA |
| K oxidation | IPR003593 | 0.01 | -2.63E-04 | 0.8 |
| K oxidation | IPR003959 | 0.2 | 9.72E-04 | 0.26 |
| K oxidation | IPR003960 | NA | 0.00E+00 | NA |
| K oxidation | IPR005225 | 0 | 5.70E-04 | 0.88 |
| K oxidation | IPR008271 | 0.16 | -2.93E-02 | 0.32 |
| K oxidation | IPR011009 | 0.18 | -2.10E-02 | 0.3 |
| K oxidation | IPR011545 | NA | 0.00E+00 | NA |
| K oxidation | IPR011989 | NA | 0.00E+00 | NA |
| K oxidation | IPR011990 | 0.06 | 9.97E-04 | 0.57 |
| K oxidation | IPR012340 | NA | 0.00E+00 | NA |
| K oxidation | IPR012677 | 0.21 | 5.19E-03 | 0.25 |
| K oxidation | IPR013766 | NA | 0.00E+00 | NA |
| K oxidation | IPR013785 | NA | 0.00E+00 | NA |
| K oxidation | IPR014001 | NA | 0.00E+00 | NA |
| K oxidation | IPR014014 | NA | 0.00E+00 | NA |
| K oxidation | IPR014729 | NA | 0.00E+00 | NA |
| K oxidation | IPR015943 | NA | 0.00E+00 | NA |
| K oxidation | IPR016024 | NA | 0.00E+00 | NA |
| K oxidation | IPR017441 | 0.16 | -2.73E-02 | 0.33 |
| K oxidation | IPR019734 | 0.06 | 1.77E-03 | 0.57 |
| K oxidation | IPR027417 | 0.05 | 1.04E-03 | 0.58 |
| K oxidation | IPR029063 | NA | 0.00E+00 | NA |
| K oxidation | IPR035979 | 0.21 | 5.19E-03 | 0.25 |
| K oxidation | IPR036249 | 0.23 | -2.50E-03 | 0.23 |
| K oxidation | IPR036291 | 0.02 | -2.16E-03 | 0.76 |
| K oxidation | IPR036322 | NA | 0.00E+00 | NA |
| M dihydroxylation | IPR000504 | 0.16 | 5.52E-03 | 0.32 |
| M dihydroxylation | IPR000719 | 0.2 | 7.59E-03 | 0.26 |
| M dihydroxylation | IPR001650 | 0.03 | 2.04E-03 | 0.67 |
| M dihydroxylation | IPR001680 | 0.46 | 8.29E-03 | 0.06 |
| M dihydroxylation | IPR002885 | NA | 0.00E+00 | NA |
| M dihydroxylation | IPR003593 | 0.82 | 2.42E-02 | 0 |
| M dihydroxylation | IPR003959 | 0.37 | 7.37E-03 | 0.11 |
| M dihydroxylation | IPR003960 | 0.3 | 7.94E-03 | 0.16 |
| M dihydroxylation | IPR005225 | 0.06 | 1.93E-03 | 0.57 |
| M dihydroxylation | IPR008271 | 0.2 | 9.42E-03 | 0.26 |
| M dihydroxylation | IPR011009 | 0.2 | 6.08E-03 | 0.26 |
| M dihydroxylation | IPR011545 | 0.03 | 1.98E-03 | 0.69 |
| M dihydroxylation | IPR011989 | 0.13 | 8.80E-03 | 0.39 |
| M dihydroxylation | IPR011990 | 0.04 | 4.75E-03 | 0.62 |
| M dihydroxylation | IPR012340 | 0.48 | 1.81E-02 | 0.06 |
| M dihydroxylation | IPR012677 | 0.16 | 5.52E-03 | 0.32 |
| M dihydroxylation | IPR013766 | 0.07 | -5.44E-03 | 0.51 |
| M dihydroxylation | IPR013785 | 0.41 | 3.01E-02 | 0.09 |
| M dihydroxylation | IPR014001 | 0.03 | 1.63E-03 | 0.71 |
| M dihydroxylation | IPR014014 | 0.03 | 2.25E-03 | 0.7 |
| M dihydroxylation | IPR014729 | 0.02 | 3.10E-03 | 0.71 |
| M dihydroxylation | IPR015943 | 0.46 | 7.28E-03 | 0.06 |
| M dihydroxylation | IPR016024 | 0.12 | 6.96E-03 | 0.4 |
| M dihydroxylation | IPR017441 | 0.2 | 9.11E-03 | 0.26 |
| M dihydroxylation | IPR019734 | 0.11 | 1.28E-02 | 0.41 |
| M dihydroxylation | IPR027417 | 0.86 | 1.58E-02 | 0 |
| M dihydroxylation | IPR029063 | 0.06 | 6.92E-03 | 0.57 |
| M dihydroxylation | IPR035979 | 0.16 | 5.52E-03 | 0.32 |
| M dihydroxylation | IPR036249 | 0.05 | -3.52E-03 | 0.61 |
| M dihydroxylation | IPR036291 | 0.4 | 1.11E-02 | 0.09 |
| M dihydroxylation | IPR036322 | 0.46 | 8.29E-03 | 0.06 |
| M oxidation | IPR000504 | 0.96 | 5.28E-01 | 0 |
| M oxidation | IPR000719 | 0.57 | 1.83E-01 | 0.03 |
| M oxidation | IPR001650 | 0.9 | 5.58E-01 | 0 |
| M oxidation | IPR001680 | 0.54 | 2.21E-01 | 0.04 |
| M oxidation | IPR002885 | 0.8 | 1.09E+00 | 0 |
| M oxidation | IPR003593 | 0.89 | 5.39E-01 | 0 |
| M oxidation | IPR003959 | 0.88 | 4.95E-01 | 0 |
| M oxidation | IPR003960 | 0.89 | 5.76E-01 | 0 |
| M oxidation | IPR005225 | 0.99 | 6.25E-01 | 0 |
| M oxidation | IPR008271 | 0.13 | 9.31E-02 | 0.39 |
| M oxidation | IPR011009 | 0.64 | 2.54E-01 | 0.02 |
| M oxidation | IPR011545 | 0.91 | 5.63E-01 | 0 |
| M oxidation | IPR011989 | 0.52 | 2.52E-01 | 0.04 |
| M oxidation | IPR011990 | 0.94 | 7.25E-01 | 0 |
| M oxidation | IPR012340 | 0.68 | 4.53E-01 | 0.01 |
| M oxidation | IPR012677 | 0.96 | 5.28E-01 | 0 |
| M oxidation | IPR013766 | 0.15 | 6.14E-02 | 0.35 |
| M oxidation | IPR013785 | 0.89 | 6.61E-01 | 0 |
| M oxidation | IPR014001 | 0.91 | 5.70E-01 | 0 |
| M oxidation | IPR014014 | 0.93 | 5.62E-01 | 0 |
| M oxidation | IPR014729 | 0 | -1.12E-02 | 0.92 |
| M oxidation | IPR015943 | 0.63 | 2.30E-01 | 0.02 |
| M oxidation | IPR016024 | 0.58 | 2.90E-01 | 0.03 |
| M oxidation | IPR017441 | 0.16 | 9.77E-02 | 0.33 |
| M oxidation | IPR019734 | 0.89 | 5.83E-01 | 0 |
| M oxidation | IPR027417 | 0.9 | 5.02E-01 | 0 |
| M oxidation | IPR029063 | 0.67 | 1.11E+00 | 0.01 |
| M oxidation | IPR035979 | 0.96 | 5.28E-01 | 0 |
| M oxidation | IPR036249 | 0.44 | 1.18E-01 | 0.07 |
| M oxidation | IPR036291 | 0.99 | 5.52E-01 | 0 |
| M oxidation | IPR036322 | 0.45 | 2.10E-01 | 0.07 |
| NTA | IPR000504 | 0.24 | -2.09E-02 | 0.22 |
| NTA | IPR000719 | NA | 0.00E+00 | NA |
| NTA | IPR001650 | 0.55 | -1.94E-02 | 0.04 |
| NTA | IPR001680 | 0.04 | -6.74E-03 | 0.65 |
| NTA | IPR002885 | NA | 0.00E+00 | NA |
| NTA | IPR003593 | 0.16 | -1.05E-02 | 0.32 |
| NTA | IPR003959 | 0.13 | -9.74E-03 | 0.38 |
| NTA | IPR003960 | 0.02 | -5.97E-03 | 0.73 |
| NTA | IPR005225 | NA | 0.00E+00 | NA |
| NTA | IPR008271 | NA | 0.00E+00 | NA |
| NTA | IPR011009 | 0.09 | 1.38E-02 | 0.47 |
| NTA | IPR011545 | 0.55 | -1.99E-02 | 0.03 |
| NTA | IPR011989 | 0.09 | -1.02E-02 | 0.48 |
| NTA | IPR011990 | NA | 0.00E+00 | NA |
| NTA | IPR012340 | 0.19 | 5.40E-02 | 0.28 |
| NTA | IPR012677 | 0.18 | -1.78E-02 | 0.29 |
| NTA | IPR013766 | 0.36 | -5.19E-02 | 0.12 |
| NTA | IPR013785 | 0.06 | -1.14E-02 | 0.57 |
| NTA | IPR014001 | 0.54 | -1.75E-02 | 0.04 |
| NTA | IPR014014 | 0.5 | -1.66E-02 | 0.05 |
| NTA | IPR014729 | 0.28 | 8.67E-03 | 0.18 |
| NTA | IPR015943 | 0.04 | -6.11E-03 | 0.65 |
| NTA | IPR016024 | 0.15 | -9.85E-03 | 0.35 |
| NTA | IPR017441 | NA | 0.00E+00 | NA |
| NTA | IPR019734 | NA | 0.00E+00 | NA |
| NTA | IPR027417 | 0.19 | -4.74E-03 | 0.29 |
| NTA | IPR029063 | NA | 0.00E+00 | NA |
| NTA | IPR035979 | 0.24 | -2.09E-02 | 0.22 |
| NTA | IPR036249 | 0.55 | -4.54E-02 | 0.03 |
| NTA | IPR036291 | 0.25 | -1.08E-02 | 0.21 |
| NTA | IPR036322 | 0.04 | -6.38E-03 | 0.65 |
| P oxidation | IPR000504 | 0.01 | -1.68E-03 | 0.82 |
| P oxidation | IPR000719 | 0.28 | -1.73E-02 | 0.18 |
| P oxidation | IPR001650 | 0.13 | -8.85E-03 | 0.38 |
| P oxidation | IPR001680 | 0.01 | -1.70E-03 | 0.85 |
| P oxidation | IPR002885 | 0.06 | 3.38E-03 | 0.57 |
| P oxidation | IPR003593 | 0.04 | -7.12E-03 | 0.64 |
| P oxidation | IPR003959 | 0.18 | -2.02E-02 | 0.29 |
| P oxidation | IPR003960 | 0.03 | 1.01E-02 | 0.66 |
| P oxidation | IPR005225 | 0.32 | 2.38E-02 | 0.15 |
| P oxidation | IPR008271 | 0.28 | -2.05E-02 | 0.18 |
| P oxidation | IPR011009 | 0.28 | -1.45E-02 | 0.18 |
| P oxidation | IPR011545 | 0.13 | -9.10E-03 | 0.38 |
| P oxidation | IPR011989 | 0.2 | 6.68E-03 | 0.26 |
| P oxidation | IPR011990 | 0.18 | -1.19E-02 | 0.29 |
| P oxidation | IPR012340 | 0.05 | -3.95E-03 | 0.59 |
| P oxidation | IPR012677 | 0.01 | -1.74E-03 | 0.81 |
| P oxidation | IPR013766 | 0.04 | -6.99E-03 | 0.66 |
| P oxidation | IPR013785 | 0.44 | 5.96E-02 | 0.07 |
| P oxidation | IPR014001 | 0.12 | -8.09E-03 | 0.39 |
| P oxidation | IPR014014 | 0.14 | -1.12E-02 | 0.36 |
| P oxidation | IPR014729 | 0.64 | -5.86E-02 | 0.02 |
| P oxidation | IPR015943 | 0.01 | -1.32E-03 | 0.86 |
| P oxidation | IPR016024 | 0.2 | 5.16E-03 | 0.26 |
| P oxidation | IPR017441 | 0.27 | -1.95E-02 | 0.18 |
| P oxidation | IPR019734 | 0.21 | -1.94E-02 | 0.25 |
| P oxidation | IPR027417 | 0.12 | -9.86E-03 | 0.39 |
| P oxidation | IPR029063 | 0.2 | 3.04E-02 | 0.26 |
| P oxidation | IPR035979 | 0.01 | -1.68E-03 | 0.82 |
| P oxidation | IPR036249 | 0.21 | -1.01E-02 | 0.26 |
| P oxidation | IPR036291 | 0 | 8.98E-04 | 0.96 |
| P oxidation | IPR036322 | 0.01 | -1.59E-03 | 0.85 |
| T oxidation | IPR000504 | 0 | -1.18E-03 | 0.92 |
| T oxidation | IPR000719 | 0.1 | -9.94E-03 | 0.44 |
| T oxidation | IPR001650 | 0.36 | 1.90E-02 | 0.11 |
| T oxidation | IPR001680 | 0.03 | -4.37E-03 | 0.69 |
| T oxidation | IPR002885 | NA | 0.00E+00 | NA |
| T oxidation | IPR003593 | 0.24 | 5.40E-03 | 0.21 |
| T oxidation | IPR003959 | 0 | -2.45E-04 | 0.97 |
| T oxidation | IPR003960 | 0.16 | 1.07E-02 | 0.33 |
| T oxidation | IPR005225 | 0.06 | -4.88E-03 | 0.56 |
| T oxidation | IPR008271 | NA | 0.00E+00 | NA |
| T oxidation | IPR011009 | 0.1 | -8.43E-03 | 0.44 |
| T oxidation | IPR011545 | 0.35 | 1.92E-02 | 0.12 |
| T oxidation | IPR011989 | 0.13 | -9.59E-03 | 0.39 |
| T oxidation | IPR011990 | 0.02 | -3.83E-03 | 0.73 |
| T oxidation | IPR012340 | 0.2 | 7.90E-03 | 0.26 |
| T oxidation | IPR012677 | 0 | -1.18E-03 | 0.92 |
| T oxidation | IPR013766 | 0.03 | -1.57E-03 | 0.69 |
| T oxidation | IPR013785 | 0.27 | 1.16E-02 | 0.18 |
| T oxidation | IPR014001 | 0.37 | 1.80E-02 | 0.11 |
| T oxidation | IPR014014 | 0.34 | 2.13E-02 | 0.13 |
| T oxidation | IPR014729 | 0.29 | 1.41E-02 | 0.17 |
| T oxidation | IPR015943 | 0.03 | -3.70E-03 | 0.69 |
| T oxidation | IPR016024 | 0.14 | -8.00E-03 | 0.36 |
| T oxidation | IPR017441 | 0.1 | -1.15E-02 | 0.44 |
| T oxidation | IPR019734 | 0.01 | -4.40E-03 | 0.8 |
| T oxidation | IPR027417 | 0.2 | 4.28E-03 | 0.27 |
| T oxidation | IPR029063 | NA | 0.00E+00 | NA |
| T oxidation | IPR035979 | 0 | -1.18E-03 | 0.92 |
| T oxidation | IPR036249 | 0.14 | -3.06E-03 | 0.36 |
| T oxidation | IPR036291 | 0.01 | 1.61E-03 | 0.84 |
| T oxidation | IPR036322 | 0.03 | -4.04E-03 | 0.69 |
| W dihydroxylation | IPR000504 | 0.82 | 7.45E-03 | 0 |
| W dihydroxylation | IPR000719 | NA | 0.00E+00 | NA |
| W dihydroxylation | IPR001650 | NA | 0.00E+00 | NA |
| W dihydroxylation | IPR001680 | 0.04 | 2.19E-03 | 0.64 |
| W dihydroxylation | IPR002885 | NA | 0.00E+00 | NA |
| W dihydroxylation | IPR003593 | 0 | 7.51E-05 | 0.96 |
| W dihydroxylation | IPR003959 | 0 | 1.18E-04 | 0.95 |
| W dihydroxylation | IPR003960 | 0 | 3.48E-04 | 0.92 |
| W dihydroxylation | IPR005225 | 0.02 | 3.14E-03 | 0.73 |
| W dihydroxylation | IPR008271 | NA | 0.00E+00 | NA |
| W dihydroxylation | IPR011009 | NA | 0.00E+00 | NA |
| W dihydroxylation | IPR011545 | NA | 0.00E+00 | NA |
| W dihydroxylation | IPR011989 | NA | 0.00E+00 | NA |
| W dihydroxylation | IPR011990 | 0.29 | 4.95E-03 | 0.17 |
| W dihydroxylation | IPR012340 | NA | 0.00E+00 | NA |
| W dihydroxylation | IPR012677 | 0.82 | 7.29E-03 | 0 |
| W dihydroxylation | IPR013766 | NA | 0.00E+00 | NA |
| W dihydroxylation | IPR013785 | NA | 0.00E+00 | NA |
| W dihydroxylation | IPR014001 | NA | 0.00E+00 | NA |
| W dihydroxylation | IPR014014 | NA | 0.00E+00 | NA |
| W dihydroxylation | IPR014729 | 0.05 | 3.09E-03 | 0.59 |
| W dihydroxylation | IPR015943 | 0.05 | 2.11E-03 | 0.61 |
| W dihydroxylation | IPR016024 | NA | 0.00E+00 | NA |
| W dihydroxylation | IPR017441 | NA | 0.00E+00 | NA |
| W dihydroxylation | IPR019734 | 0.29 | 8.76E-03 | 0.17 |
| W dihydroxylation | IPR027417 | 0.03 | 5.07E-04 | 0.7 |
| W dihydroxylation | IPR029063 | NA | 0.00E+00 | NA |
| W dihydroxylation | IPR035979 | 0.82 | 7.45E-03 | 0 |
| W dihydroxylation | IPR036249 | 0.06 | -5.25E-03 | 0.56 |
| W dihydroxylation | IPR036291 | 0.67 | -1.58E-02 | 0.01 |
| W dihydroxylation | IPR036322 | 0.05 | 2.30E-03 | 0.61 |
| W oxidation | IPR000504 | 0.82 | 7.45E-03 | 0 |
| W oxidation | IPR000719 | NA | 0.00E+00 | NA |
| W oxidation | IPR001650 | NA | 0.00E+00 | NA |
| W oxidation | IPR001680 | 0.2 | 7.34E-03 | 0.27 |
| W oxidation | IPR002885 | NA | 0.00E+00 | NA |
| W oxidation | IPR003593 | 0.05 | -2.61E-03 | 0.61 |
| W oxidation | IPR003959 | 0.05 | -3.38E-03 | 0.61 |
| W oxidation | IPR003960 | 0.03 | -4.75E-03 | 0.67 |
| W oxidation | IPR005225 | 0.03 | -1.04E-02 | 0.66 |
| W oxidation | IPR008271 | NA | 0.00E+00 | NA |
| W oxidation | IPR011009 | NA | 0.00E+00 | NA |
| W oxidation | IPR011545 | NA | 0.00E+00 | NA |
| W oxidation | IPR011989 | NA | 0.00E+00 | NA |
| W oxidation | IPR011990 | 0.27 | 4.21E-03 | 0.19 |
| W oxidation | IPR012340 | NA | 0.00E+00 | NA |
| W oxidation | IPR012677 | 0.82 | 7.29E-03 | 0 |
| W oxidation | IPR013766 | NA | 0.00E+00 | NA |
| W oxidation | IPR013785 | NA | 0.00E+00 | NA |
| W oxidation | IPR014001 | NA | 0.00E+00 | NA |
| W oxidation | IPR014014 | NA | 0.00E+00 | NA |
| W oxidation | IPR014729 | 0.34 | -3.08E-02 | 0.13 |
| W oxidation | IPR015943 | 0.21 | 6.56E-03 | 0.26 |
| W oxidation | IPR016024 | NA | 0.00E+00 | NA |
| W oxidation | IPR017441 | NA | 0.00E+00 | NA |
| W oxidation | IPR019734 | 0.36 | 8.42E-03 | 0.12 |
| W oxidation | IPR027417 | 0.05 | -1.97E-03 | 0.59 |
| W oxidation | IPR029063 | NA | 0.00E+00 | NA |
| W oxidation | IPR035979 | 0.82 | 7.45E-03 | 0 |
| W oxidation | IPR036249 | 0.03 | 2.03E-03 | 0.69 |
| W oxidation | IPR036291 | 0.56 | -3.88E-02 | 0.03 |
| W oxidation | IPR036322 | 0.21 | 7.02E-03 | 0.25 |
| pyro-Glu | IPR000504 | NA | 0.00E+00 | NA |
| pyro-Glu | IPR000719 | NA | 0.00E+00 | NA |
| pyro-Glu | IPR001650 | NA | 0.00E+00 | NA |
| pyro-Glu | IPR001680 | NA | 0.00E+00 | NA |
| pyro-Glu | IPR002885 | NA | 0.00E+00 | NA |
| pyro-Glu | IPR003593 | NA | 0.00E+00 | NA |
| pyro-Glu | IPR003959 | NA | 0.00E+00 | NA |
| pyro-Glu | IPR003960 | NA | 0.00E+00 | NA |
| pyro-Glu | IPR005225 | NA | 0.00E+00 | NA |
| pyro-Glu | IPR008271 | NA | 0.00E+00 | NA |
| pyro-Glu | IPR011009 | NA | 0.00E+00 | NA |
| pyro-Glu | IPR011545 | NA | 0.00E+00 | NA |
| pyro-Glu | IPR011989 | NA | 0.00E+00 | NA |
| pyro-Glu | IPR011990 | NA | 0.00E+00 | NA |
| pyro-Glu | IPR012340 | NA | 0.00E+00 | NA |
| pyro-Glu | IPR012677 | NA | 0.00E+00 | NA |
| pyro-Glu | IPR013766 | NA | 0.00E+00 | NA |
| pyro-Glu | IPR013785 | NA | 0.00E+00 | NA |
| pyro-Glu | IPR014001 | NA | 0.00E+00 | NA |
| pyro-Glu | IPR014014 | NA | 0.00E+00 | NA |
| pyro-Glu | IPR014729 | NA | 0.00E+00 | NA |
| pyro-Glu | IPR015943 | NA | 0.00E+00 | NA |
| pyro-Glu | IPR016024 | NA | 0.00E+00 | NA |
| pyro-Glu | IPR017441 | NA | 0.00E+00 | NA |
| pyro-Glu | IPR019734 | NA | 0.00E+00 | NA |
| pyro-Glu | IPR027417 | NA | 0.00E+00 | NA |
| pyro-Glu | IPR029063 | NA | 0.00E+00 | NA |
| pyro-Glu | IPR035979 | NA | 0.00E+00 | NA |
| pyro-Glu | IPR036249 | NA | 0.00E+00 | NA |
| pyro-Glu | IPR036291 | NA | 0.00E+00 | NA |
| pyro-Glu | IPR036322 | NA | 0.00E+00 | NA |

Supplemental Table S8. Cluster analysis of proteomic data based on changes in PTM frequencies over time. 12 clusters were identified and four are shown. Exemplar proteins are shown in blue and bold.

| TAIR Header | Cluster |
| --- | --- |
| AT3G12580.1 \| Symbols: HSP70, ATHSP70, HSC70-4 \| ARABIDOPSIS HEAT SHOCK PROTEIN 70, heat shock protein 70 \| chr3:3991487-3993689 REVERSE LENGTH=650 | 1 |
| AT5G08670.1 \| Symbols: no symbol available \| no full name available \| chr5:2818395-2821149 REVERSE LENGTH=556 | 1 |
| AT5G20890.1 \| Symbols: CCT2 \| Chaperonin containing T-complex polypeptide-1 subunit 2 \| chr5:7087020-7089906 REVERSE LENGTH=527 | 1 |
| AT3G13860.1 \| Symbols: HSP60-3A \| heat shock protein 60-3A \| chr3:4561704-4565133 REVERSE LENGTH=572 | 1 |
| AT4G02930.1 \| Symbols: no symbol available \| no full name available \| chr4:1295751-1298354 REVERSE LENGTH=454 | 1 |
| AT4G34200.1 \| Symbols: EDA9, PGDH1 \| phosphoglycerate dehydrogenase 1, embryo sac development arrest 9 \| chr4:16374041-16376561 REVERSE LENGTH=603 | 1 |
| AT3G44110.1 \| Symbols: ATJ3, ATJ, J3 \| DNAJ homologue 3 \| chr3:15869115-15871059 REVERSE LENGTH=420 | 1 |
| AT5G26860.1 \| Symbols: LON1, LON_ARA_ARA \| lon protease 1 \| chr5:9451048-9456631 FORWARD LENGTH=985 | 1 |
| AT3G11830.1 \| Symbols: CCT7 \| Chaperonin containing T-complex polypeptide-1 subunit 7 \| chr3:3732734-3736156 FORWARD LENGTH=557 | 1 |
| AT4G22670.1 \| Symbols: AtHip1, HIP1, TPR11 \| HSP70-interacting protein 1, tetratricopeptide repeat 11 \| chr4:11918236-11920671 FORWARD LENGTH=441 | 1 |
| AT4G20850.1 \| Symbols: TPP2 \| tripeptidyl peptidase ii \| chr4:11160935-11169889 REVERSE LENGTH=1380 | 1 |
| AT5G19990.1 \| Symbols: RPT6A, ATSUG1 \| regulatory particle triple-A ATPase 6A \| chr5:6752144-6754918 FORWARD LENGTH=419 | 1 |
| AT2G35040.1 \| Symbols: no symbol available \| no full name available \| chr2:14765347-14768269 REVERSE LENGTH=596 | 1 |
| AT1G59610.1 \| Symbols: DL3, DRP2B, ADL3, CF1 \| Dynamin related protein 2B, dynamin-like 3 \| chr1:21893413-21900780 FORWARD LENGTH=920 | 1 |
| AT5G11170.1 \| Symbols: UAP56a \| homolog of human UAP56 a \| chr5:3553334-3556646 FORWARD LENGTH=427 | 1 |
| AT2G47510.1 \| Symbols: FUM1 \| fumarase 1 \| chr2:19498614-19502020 FORWARD LENGTH=492 | 1 |
| AT4G27585.1 \| Symbols: SLP1, AtSLP1 \| stomatin-like protein 1 \| chr4:13766984-13769832 REVERSE LENGTH=411 | 1 |
| AT1G09620.1 \| Symbols: no symbol available \| no full name available \| chr1:3113077-3116455 REVERSE LENGTH=1091 | 1 |
| AT5G50850.1 \| Symbols: MAB1 \| MACCI-BOU \| chr5:20689671-20692976 FORWARD LENGTH=363 | 1 |
| AT4G13430.1 \| Symbols: IIL1, ATLEUC1 \| isopropyl malate isomerase large subunit 1 \| chr4:7804194-7807789 REVERSE LENGTH=509 | 1 |
| AT5G55070.1 \| Symbols: E2-OGDH2 \| \| chr5:22347637-22350409 FORWARD LENGTH=464 | 1 |
| AT2G37760.1 \| Symbols: AKR4C8 \| Aldo-keto reductase family 4 member C8 \| chr2:15831995-15833742 FORWARD LENGTH=311 | 1 |
| AT1G24510.1 \| Symbols: CCT5 \| Chaperonin containing T-complex polypeptide-1 subunit 5 \| chr1:8685504-8688101 REVERSE LENGTH=535 | 1 |
| AT1G20630.1 \| Symbols: CAT1 \| catalase 1 \| chr1:7146812-7149609 FORWARD LENGTH=492 | 1 |
| AT1G04480.1 \| Symbols: no symbol available \| no full name available \| chr1:1216110-1217257 FORWARD LENGTH=140 | 1 |
| AT1G62380.1 \| Symbols: ATACO2, ACO2 \| ACC oxidase 2 \| chr1:23082340-23084068 FORWARD LENGTH=320 | 1 |
| AT5G46290.1 \| Symbols: KASI, KAS1 \| 3-ketoacyl-acyl carrier protein synthase I, KETOACYL-ACP SYNTHASE 1 \| chr5:18774439-18776629 REVERSE LENGTH=473 | 1 |
| AT5G58070.1 \| Symbols: TIL, ATTIL \| TEMPERATURE-INDUCED LIPOCALIN, temperature-induced lipocalin \| chr5:23500512-23501156 REVERSE LENGTH=186 | 1 |
| AT5G19510.1 \| Symbols: no symbol available \| no full name available \| chr5:6581854-6583137 REVERSE LENGTH=224 | 1 |
| AT1G62750.1 \| Symbols: ATSCO1/CPEF-G, ATSCO1, SCO1 \| SNOWY COTYLEDON 1 \| chr1:23233622-23236321 REVERSE LENGTH=783 | 1 |
| AT3G59760.1 \| Symbols: OASC, ATCS-C \| ARABIDOPSIS THALIANA CYSTEINSYNTHASE-C, O-acetylserine (thiol) lyase isoform C \| chr3:22072119-22075345 REVERSE LENGTH=433 | 1 |
| AT5G63680.1 \| Symbols: no symbol available \| no full name available \| chr5:25490507-25492530 FORWARD LENGTH=510 | 1 |
| AT1G63660.1 \| Symbols: no symbol available \| no full name available \| chr1:23604127-23607080 REVERSE LENGTH=534 | 1 |
| AT2G38040.1 \| Symbols: CAC3 \| acetyl Co-enzyme a carboxylase carboxyltransferase alpha subunit \| chr2:15917612-15920749 FORWARD LENGTH=769 | 1 |
| AT3G25860.1 \| Symbols: LTA2, PLE2 \| PLASTID E2 SUBUNIT OF PYRUVATE DECARBOXYLASE \| chr3:9460632-9462585 FORWARD LENGTH=480 | 1 |
| AT3G03060.1 \| Symbols: SBA1, ATAD3A1 \| ATPase family AAA Domain-containing protein 3A1, SHOT1 Binding ATPase 1 \| chr3:692188-695424 FORWARD LENGTH=628 | 1 |
| AT5G42890.1 \| Symbols: SCP2, ATSCP2 \| STEROL CARRIER PROTEIN 2, sterol carrier protein 2 \| chr5:17194458-17195910 REVERSE LENGTH=123 | 1 |
| AT5G04800.1 \| Symbols: no symbol available \| no full name available \| chr5:1389217-1389642 FORWARD LENGTH=141 | 1 |
| AT1G74030.1 \| Symbols: ENO1 \| enolase 1 \| chr1:27839465-27841901 REVERSE LENGTH=477 | 1 |
| AT1G20370.1 \| Symbols: no symbol available \| no full name available \| chr1:7051846-7053588 REVERSE LENGTH=549 | 1 |
| AT3G60240.3 \| Symbols: EIF4G, CUM2 \| eukaryotic translation initiation factor 4G, CUCUMOVIRUS MULTIPLICATION 2 \| chr3:22261842-22268295 FORWARD LENGTH=1725 | 1 |
| AT3G46230.1 \| Symbols: HSP17.4, ATHSP17.4 \| heat shock protein 17.4, ARABIDOPSIS THALIANA HEAT SHOCK PROTEIN 17.4 \| chr3:16984263-16984733 REVERSE LENGTH=156 | 1 |
| AT4G13930.1 \| Symbols: SHM4 \| serine hydroxymethyltransferase 4 \| chr4:8048013-8050021 REVERSE LENGTH=471 | 1 |
| AT1G79280.1 \| Symbols: AtTPR, NUA \| TRANSLOCATED PROMOTER REGION, nuclear pore anchor \| chr1:29819176-29832809 REVERSE LENGTH=2093 | 1 |
| AT1G30230.1 \| Symbols: EF1Bb, eEF-1Bb1 \| eukaryotic elongation factor 1B beta 1, elongation factor 1B beta \| chr1:10639286-10640515 FORWARD LENGTH=231 | 1 |
| AT2G18330.1 \| Symbols: SBA3, ATAD3B1 \| ATPase family AAA Domain-containing protein 3B1, SHOT1 Binding ATPase 3 \| chr2:7965829-7968915 FORWARD LENGTH=636 | 1 |
| AT4G26780.1 \| Symbols: MGE2, AR192 \| mitochondrial GrpE 2 \| chr4:13485066-13486560 REVERSE LENGTH=327 | 1 |
| AT3G48870.1 \| Symbols: ATCLPC, HSP93-III, ClpC2, ATHSP93-III \| ClpC2 \| chr3:18122363-18126008 REVERSE LENGTH=952 | 1 |
| AT3G16050.1 \| Symbols: ATPDX1.2, A37, PDX1.2 \| pyridoxine biosynthesis 1.2, ARABIDOPSIS THALIANA PYRIDOXINE BIOSYNTHESIS 1.2 \| chr3:5444121-5445065 REVERSE LENGTH=314 | 1 |
| AT5G21990.1 \| Symbols: AtTPR7, TPR7, OEP61 \| tetratricopeptide repeat 7, outer envelope protein 61 \| chr5:7273395-7276318 FORWARD LENGTH=554 | 1 |
| AT5G19320.1 \| Symbols: RANGAP2 \| RAN GTPase activating protein 2 \| chr5:6505310-6506947 REVERSE LENGTH=545 | 1 |
| AT3G10690.1 \| Symbols: GYRA \| DNA GYRASE A \| chr3:3339612-3346243 REVERSE LENGTH=950 | 1 |
| AT1G29030.1 \| Symbols: no symbol available \| no full name available \| chr1:10129201-10133697 REVERSE LENGTH=556 | 1 |
| AT2G25970.1 \| Symbols: no symbol available \| no full name available \| chr2:11071844-11075604 REVERSE LENGTH=632 | 1 |
| AT1G07360.1 \| Symbols: MAC5A \| MOS4-associated complex subunit 5A \| chr1:2260562-2262795 REVERSE LENGTH=481 | 1 |
| AT1G18500.1 \| Symbols: MAML-4, IPMS1 \| methylthioalkylmalate synthase-like 4, ISOPROPYLMALATE SYNTHASE 1 \| chr1:6369347-6372861 FORWARD LENGTH=631 | 1 |
| AT1G16350.1 \| Symbols: no symbol available \| no full name available \| chr1:5590951-5592872 FORWARD LENGTH=502 | 1 |
| AT3G60450.2 \| Symbols: no symbol available \| no full name available \| chr3:22340886-22342187 FORWARD LENGTH=306 | 1 |
| AT3G49560.1 \| Symbols: Tric1, HP30 \| tRNA import component 1, hypothetical protein 30 \| chr3:18370644-18371821 FORWARD LENGTH=261 | 1 |
| AT1G07510.1 \| Symbols: FTSH10, AtFTSH10 \| FTSH protease 10 \| chr1:2305689-2309380 FORWARD LENGTH=813 | 1 |
| AT4G33010.1 \| Symbols: GLDP1, AtGLDP1 \| glycine decarboxylase P-protein 1 \| chr4:15926852-15931150 REVERSE LENGTH=1037 | 1 |
| AT5G60730.1 \| Symbols: AtGET3c, GET3c \| Guided Entry of Tail-anchored proteins 3b, Guided Entry of Tail-anchored protein 3c \| chr5:24422838-24425352 FORWARD LENGTH=391 | 1 |
| AT4G29010.1 \| Symbols: AIM1 \| ABNORMAL INFLORESCENCE MERISTEM \| chr4:14297312-14302016 REVERSE LENGTH=721 | 1 |
| AT5G56740.1 \| Symbols: HAC07, HAC7, HAG2, HAG02 \| histone acetyltransferase of the GNAT family 2 \| chr5:22953009-22955577 REVERSE LENGTH=467 | 1 |
| AT1G48850.1 \| Symbols: EMB1144 \| embryo defective 1144 \| chr1:18065154-18067956 REVERSE LENGTH=436 | 1 |
| AT2G45240.1 \| Symbols: MAP1A \| methionine aminopeptidase 1A \| chr2:18656059-18658906 FORWARD LENGTH=398 | 1 |
| AT3G48170.1 \| Symbols: ALDH10A9, BADH \| BETAINE ALDEHYDE DEHYDROGENASE, aldehyde dehydrogenase 10A9 \| chr3:17786290-17789918 REVERSE LENGTH=503 | 1 |
| AT1G21080.1 \| Symbols: no symbol available \| no full name available \| chr1:7378822-7382275 REVERSE LENGTH=391 | 1 |
| AT4G31160.1 \| Symbols: DCAF1 \| DDB1-CUL4 associated factor 1 \| chr4:15145936-15152939 FORWARD LENGTH=1883 | 1 |
| AT4G33650.2 \| Symbols: DRP3A, NOXY15, ADL2, APEM1 \| dynamin-related protein 3A, non responding to oxylipins 15, ARABIDOPSIS DYNAMIN-LIKE 2, ABERRANT PEROXISOME MORPHOLOGY 1 \| chr4:16161073-16166587 FORWARD LENGTH=809 | 1 |
| AT5G58710.1 \| Symbols: ROC7 \| rotamase CYP 7 \| chr5:23717840-23719495 FORWARD LENGTH=204 | 1 |
| AT1G48030.1 \| Symbols: mtLPD1 \| mitochondrial lipoamide dehydrogenase 1 \| chr1:17717432-17719141 REVERSE LENGTH=507 | 1 |
| AT2G39990.1 \| Symbols: EIF2, AteIF3f, eIF3F \| Arabidopsis thaliana eukaryotic translation initiation factor 3 subunit F, eukaryotic translation initiation factor 2, eukaryotic translation initiation factor 3 subunit F \| chr2:16698332-16699929 REVERSE LENGTH=293 | 1 |
| AT3G25530.1 \| Symbols: AtGLYR1, GLYR1, ATGHBDH, GHBDH, GR1 \| GLYOXYLATE REDUCTASE 1, glyoxylate reductase 1 \| chr3:9271949-9273514 REVERSE LENGTH=289 | 1 |
| AT1G09770.1 \| Symbols: ATCDC5, ATMYBCDC5, CDC5 \| ARABIDOPSIS THALIANA CELL DIVISION CYCLE 5, cell division cycle 5, ARABIDOPSIS THALIANA MYB DOMAIN CELL DIVISION CYCLE 5 \| chr1:3162002-3165122 FORWARD LENGTH=844 | 1 |
| AT5G47030.1 \| Symbols: no symbol available \| no full name available \| chr5:19090384-19092034 FORWARD LENGTH=203 | 1 |
| AT5G04430.1 \| Symbols: BTR1, BTR1L, BTR1S \| BINDING TO TOMV RNA 1S (SHORT FORM), binding to TOMV RNA 1L (long form), BINDING TO TOMV RNA 1 \| chr5:1250602-1253523 REVERSE LENGTH=313 | 1 |
| AT1G66680.1 \| Symbols: AR401 \| \| chr1:24866352-24868888 REVERSE LENGTH=358 | 1 |
| AT3G07090.1 \| Symbols: Desi1 \| \| chr3:2243153-2244476 REVERSE LENGTH=265 | 1 |
| AT3G55620.1 \| Symbols: emb1624, eIF6A \| embryo defective 1624, eukaryotic initiation facor 6A \| chr3:20634581-20636312 FORWARD LENGTH=245 | 1 |
| AT1G10840.1 \| Symbols: TIF3H1 \| translation initiation factor 3 subunit H1 \| chr1:3607885-3610299 REVERSE LENGTH=337 | 1 |
| AT2G47970.1 \| Symbols: no symbol available \| no full name available \| chr2:19629525-19630973 FORWARD LENGTH=413 | 1 |
| AT4G24280.1 \| Symbols: cpHsc70-1 \| chloroplast heat shock protein 70-1 \| chr4:12590094-12593437 FORWARD LENGTH=718 | 1 |
| AT5G42950.1 \| Symbols: EXA1 \| Essential for poteXvirus Accumulation 1 \| chr5:17224436-17231044 FORWARD LENGTH=1714 | 1 |
| AT1G69250.1 \| Symbols: no symbol available \| no full name available \| chr1:26033163-26035301 FORWARD LENGTH=427 | 1 |
| AT1G65540.1 \| Symbols: AtLETM2, LETM2 \| leucine zipper-EF-hand-containing transmembrane protein 2 \| chr1:24362382-24366011 REVERSE LENGTH=736 | 1 |
| AT2G43130.1 \| Symbols: ARA-4, RABA5C, ATRAB11F, ARA4, ATRABA5C \| ARABIDOPSIS RAB GTPASE HOMOLOG A5C \| chr2:17929899-17930904 REVERSE LENGTH=214 | 1 |
| AT1G77550.1 \| Symbols: no symbol available \| no full name available \| chr1:29138490-29142965 REVERSE LENGTH=855 | 1 |
| AT5G10240.1 \| Symbols: ASN3 \| asparagine synthetase 3 \| chr5:3212934-3216418 REVERSE LENGTH=578 | 1 |
| AT4G26110.1 \| Symbols: NAP1;1, ATNAP1;1 \| ARABIDOPSIS THALIANA NUCLEOSOME ASSEMLY PROTEIN 1;1, nucleosome assembly protein1;1 \| chr4:13232712-13235502 FORWARD LENGTH=372 | 1 |
| AT3G12800.1 \| Symbols: SDRB, DECR \| short-chain dehydrogenase-reductase B \| chr3:4063463-4064757 REVERSE LENGTH=298 | 1 |
| AT5G48030.1 \| Symbols: GFA2 \| gametophytic factor 2 \| chr5:19466298-19469753 REVERSE LENGTH=456 | 1 |
| AT4G17010.1 \| Symbols: no symbol available \| no full name available \| chr4:9576442-9577627 FORWARD LENGTH=160 | 1 |
| AT4G08350.1 \| Symbols: GTA2, SPT5-2, GTA02 \| global transcription factor group A2 \| chr4:5286351-5292072 FORWARD LENGTH=1041 | 1 |
| AT1G43700.1 \| Symbols: SUE3, VIP1, AtVIP1 \| VIRE2-interacting protein 1, sulphate utilization efficiency 3 \| chr1:16484352-16486017 FORWARD LENGTH=341 | 1 |
| AT2G29560.1 \| Symbols: ENOC, ENO3 \| cytosolic enolase, enolase 3 \| chr2:12646635-12649694 FORWARD LENGTH=475 | 1 |
| AT1G79940.1 \| Symbols: Sec. 63-1, ATERDJ2A \| \| chr1:30070023-30073237 FORWARD LENGTH=687 | 1 |
| AT5G63890.1 \| Symbols: HISN8, ATHDH, HDH \| histidinol dehydrogenase, HISTIDINE BIOSYNTHESIS 8 \| chr5:25565600-25567879 REVERSE LENGTH=452 | 1 |
| AT2G47730.1 \| Symbols: GST6, GSTF8, ATGSTF8, ATGSTF5 \| glutathione S-transferase phi 8, Arabidopsis thaliana glutathione S-transferase phi 8, GLUTATHIONE S-TRANSFERASE (CLASS PHI) 5 \| chr2:19558213-19559266 FORWARD LENGTH=263 | 1 |
| AT4G18950.1 \| Symbols: BHP \| BLUE LIGHT-DEPENDENT H+-ATPASE PHOSPHORYLATION \| chr4:10375685-10378129 FORWARD LENGTH=459 | 1 |
| AT5G11240.1 \| Symbols: AtGHS40, NuGWD1, GHS40 \| Nuclear Glucose-responsive WD40 protein1, GLUCOSE HYPERSENSITIVE 40 \| chr5:3582949-3586782 FORWARD LENGTH=615 | 1 |
| AT3G48890.1 \| Symbols: MSBP2, MAPR3, ATMAPR3, ATMP2 \| ARABIDOPSIS THALIANA MEMBRANE-ASSOCIATED PROGESTERONE BINDING PROTEIN 3, membrane-associated progesterone binding protein 3, MEMBRANE STEROID BINDING PROTEIN 2 \| chr3:18129669-18131353 FORWARD LENGTH=233 | 1 |
| AT5G19485.1 \| Symbols: no symbol available \| no full name available \| chr5:6573907-6576352 REVERSE LENGTH=456 | 1 |
| AT2G19730.1 \| Symbols: no symbol available \| no full name available \| chr2:8511752-8512995 FORWARD LENGTH=143 | 1 |
| AT5G42220.1 \| Symbols: no symbol available \| no full name available \| chr5:16872962-16877455 FORWARD LENGTH=879 | 1 |
| AT1G43860.1 \| Symbols: no symbol available \| no full name available \| chr1:16622244-16624385 REVERSE LENGTH=370 | 1 |
| AT1G50380.1 \| Symbols: no symbol available \| no full name available \| chr1:18662480-18666185 FORWARD LENGTH=710 | 1 |
| AT1G27390.1 \| Symbols: TOM20-2 \| translocase outer membrane 20-2 \| chr1:9513469-9514912 REVERSE LENGTH=210 | 1 |
| AT1G63500.1 \| Symbols: BSK7 \| brassinosteroid-signaling kinase 7 \| chr1:23556015-23558403 FORWARD LENGTH=487 | 1 |
| AT1G71860.1 \| Symbols: PTP1, ATPTP1 \| protein tyrosine phosphatase 1 \| chr1:27026866-27028675 FORWARD LENGTH=340 | 1 |
| AT2G24940.1 \| Symbols: MAPR2, AtMAPR2 \| membrane-associated progesterone binding protein 2 \| chr2:10609447-10609749 FORWARD LENGTH=100 | 1 |
| AT5G58590.1 \| Symbols: RANBP1 \| RAN binding protein 1 \| chr5:23680319-23681714 REVERSE LENGTH=219 | 1 |
| AT5G41950.1 \| Symbols: HLB1 \| HYPERSENSITIVE TO LATRUNCULIN B 1 \| chr5:16785825-16789360 FORWARD LENGTH=565 | 1 |
| AT3G01540.1 \| Symbols: IRP6, RH14, DRH1, ATDRH1 \| ARABIDOPSIS THALIANA DEAD BOX RNA HELICASE 1, RNA Helicase 14, DEAD box RNA helicase 1, NVOLVED IN rRNA PROCESSING 6 \| chr3:213077-216142 REVERSE LENGTH=618 | 1 |
| AT1G67230.1 \| Symbols: CRWN1, KAKU2, LINC1, AtLINC1 \| (Japanese for nucleus) 2, CROWDED NUCLEI 1, LITTLE NUCLEI1 \| chr1:25151561-25156032 REVERSE LENGTH=1132 | 1 |
| AT3G22980.1 \| Symbols: no symbol available \| no full name available \| chr3:8160269-8163316 REVERSE LENGTH=1015 | 1 |
| AT4G15940.1 \| Symbols: FAHD1a \| fumarylacetoacetate hydrolase domain containing protein 1a \| chr4:9038361-9040163 FORWARD LENGTH=222 | 1 |
| AT2G31610.1 \| Symbols: no symbol available \| no full name available \| chr2:13450384-13451669 FORWARD LENGTH=250 | 1 |
| AT2G17130.1 \| Symbols: IDH2, IDH-II \| isocitrate dehydrogenase subunit 2, isocitrate dehydrogenase II \| chr2:7461062-7462466 REVERSE LENGTH=367 | 1 |
| AT2G04842.1 \| Symbols: EMB2761 \| EMBRYO DEFECTIVE 2761 \| chr2:1698466-1701271 REVERSE LENGTH=650 | 1 |
| AT2G22450.1 \| Symbols: RIBA2, AtRIBA2 \| homolog of ribA 2 \| chr2:9530654-9532691 FORWARD LENGTH=476 | 1 |
| AT5G46750.1 \| Symbols: AGD9 \| ARF-GAP domain 9 \| chr5:18969950-18971817 REVERSE LENGTH=402 | 1 |
| AT5G39730.1 \| Symbols: no symbol available \| no full name available \| chr5:15901740-15902624 FORWARD LENGTH=172 | 1 |
| AT2G25840.1 \| Symbols: OVA4 \| ovule abortion 4 \| chr2:11021924-11025158 FORWARD LENGTH=408 | 1 |
| AT5G65620.1 \| Symbols: OOP, TOP1 \| thimet metalloendopeptidase 1, organellar oligopeptidase \| chr5:26221951-26225784 FORWARD LENGTH=791 | 1 |
| AT1G05520.1 \| Symbols: AtSEC23B \| \| chr1:1631126-1635703 REVERSE LENGTH=783 | 1 |
| AT5G56360.1 \| Symbols: PSL4 \| PRIORITY IN SWEET LIFE 4 \| chr5:22823586-22827950 REVERSE LENGTH=647 | 1 |
| AT1G26460.1 \| Symbols: no symbol available \| no full name available \| chr1:9151816-9154407 FORWARD LENGTH=630 | 1 |
| AT2G23820.2 \| Symbols: no symbol available \| no full name available \| chr2:10140595-10142672 FORWARD LENGTH=257 | 1 |
| AT5G03660.1 \| Symbols: no symbol available \| no full name available \| chr5:938068-939811 FORWARD LENGTH=173 | 1 |
| AT1G73030.1 \| Symbols: CHMP1A, VPS46.2 \| CHARGED MULTIVESICULAR BODY PROTEIN/CHROMATIN MODIFYING PROTEIN1A \| chr1:27473938-27474848 FORWARD LENGTH=203 | 1 |
| AT5G19370.1 \| Symbols: no symbol available \| no full name available \| chr5:6524247-6526629 REVERSE LENGTH=299 | 1 |
| AT4G23430.2 \| Symbols: TIC32, AtTic32-IVa, Tic32-IVa \| translocon at the inner envelope membrane of chloroplasts 32-IVa \| chr4:12229171-12231493 FORWARD LENGTH=322 | 1 |
| AT3G07100.1 \| Symbols: SEC24A, ERMO2, AtSEC24A \| ENDOPLASMIC RETICULUM MORPHOLOGY 2 \| chr3:2245689-2250077 REVERSE LENGTH=1038 | 1 |
| AT1G10590.1 \| Symbols: no symbol available \| no full name available \| chr1:3502324-3502991 REVERSE LENGTH=139 | 1 |
| AT3G49430.1 \| Symbols: SR34a, SRp34a, At-SR34a \| Serine/Arginine-Rich Protein Splicing Factor 34a, SER/ARG-rich protein 34A \| chr3:18332668-18334829 FORWARD LENGTH=300 | 1 |
| AT1G49600.1 \| Symbols: RBP47A, ATRBP47A \| RNA-binding protein 47A \| chr1:18357236-18360150 REVERSE LENGTH=445 | 1 |
| AT3G08947.1 \| Symbols: no symbol available \| no full name available \| chr3:2724654-2726722 FORWARD LENGTH=657 | 1 |
| AT4G39260.1 \| Symbols: RBGA6, CCR1, ATGRP8, GR-RBP8, GRP8 \| "cold, circadian rhythm, and RNA binding 1", glycine-rich RNA-binding protein 8, GLYCINE-RICH PROTEIN 8, RNA-binding glycine-rich protein A6 \| chr4:18274166-18274958 REVERSE LENGTH=169 | 1 |
| AT1G02500.1 \| Symbols: METK1, SAM-1, AtSAM1, SAM1, MAT1 \| S-adenosylmethionine synthetase 1, S-ADENOSYLMETHIONINE SYNTHETASE-1 \| chr1:519037-520218 FORWARD LENGTH=393 | 1 |
| AT2G38230.1 \| Symbols: ATPDX1.1, PDX1.1 \| pyridoxine biosynthesis 1.1, ARABIDOPSIS THALIANA PYRIDOXINE BIOSYNTHESIS 1.1 \| chr2:16011475-16012404 FORWARD LENGTH=309 | 1 |
| AT2G39080.1 \| Symbols: EMB2799 \| EMBRYO DEFECTIVE 2799 \| chr2:16309981-16312475 REVERSE LENGTH=351 | 1 |
| AT3G56090.1 \| Symbols: FER3, ATFER3 \| ferritin 3 \| chr3:20814350-20815984 REVERSE LENGTH=259 | 1 |
| AT4G11260.1 \| Symbols: EDM1, RPR1, ATSGT1B, SGT1B, ETA3 \| ENHANCED DOWNY MILDEW 1, ENHANCER OF TIR1-1 AUXIN RESISTANCE 3 \| chr4:6851515-6853719 REVERSE LENGTH=358 | 1 |
| AT3G12260.1 \| Symbols: NDUFA6, B14 \| \| chr3:3909252-3910337 REVERSE LENGTH=133 | 1 |
| AT5G30510.1 \| Symbols: ARRPS1, RPS1, PRPS1 \| plastid ribosomal protein S1, ribosomal protein S1 \| chr5:11619262-11621223 REVERSE LENGTH=416 | 1 |
| AT3G49100.1 \| Symbols: no symbol available \| no full name available \| chr3:18196944-18198298 FORWARD LENGTH=103 | 1 |
| AT1G24800.1 \| Symbols: no symbol available \| no full name available \| chr1:8769741-8771042 FORWARD LENGTH=433 | 1 |
| AT5G24810.1 \| Symbols: ABC1K11 \| \| chr5:8516902-8522616 REVERSE LENGTH=1009 | 1 |
| AT3G28480.1 \| Symbols: no symbol available \| no full name available \| chr3:10676266-10678262 REVERSE LENGTH=316 | 1 |
| AT1G21720.1 \| Symbols: PBC1 \| proteasome beta subunit C1 \| chr1:7626394-7628070 FORWARD LENGTH=204 | 1 |
| AT1G08560.1 \| Symbols: SYP111, ATSYP111, KN \| syntaxin of plants 111, KNOLLE \| chr1:2709778-2710710 REVERSE LENGTH=310 | 1 |
| AT1G07830.1 \| Symbols: no symbol available \| no full name available \| chr1:2422549-2423392 FORWARD LENGTH=144 | 1 |
| AT2G47990.1 \| Symbols: SWA1, EDA13, EDA19 \| SLOW WALKER1, EMBRYO SAC DEVELOPMENT ARREST 19, EMBRYO SAC DEVELOPMENT ARREST 13 \| chr2:19637010-19638602 REVERSE LENGTH=530 | 1 |
| AT3G09980.1 \| Symbols: ACIP1 \| ACETYLATED INTERACTING PROTEIN 1 \| chr3:3069358-3071145 FORWARD LENGTH=178 | 1 |
| AT5G23290.1 \| Symbols: PFD5 \| prefoldin 5 \| chr5:7846144-7847428 FORWARD LENGTH=151 | 1 |
| AT2G29960.1 \| Symbols: ATCYP5, CYP5, CYP19-4 \| ARABIDOPSIS THALIANA CYCLOPHILIN 5, cyclophilin 5, CYCLOPHILIN 19-4 \| chr2:12769183-12770528 REVERSE LENGTH=201 | 1 |
| AT3G47370.1 \| Symbols: no symbol available \| no full name available \| chr3:17453671-17454437 REVERSE LENGTH=122 | 1 |
| AT3G53880.1 \| Symbols: AKR4C11 \| Aldo-keto reductase family 4 member C11 \| chr3:19953238-19955171 FORWARD LENGTH=315 | 1 |
| AT2G04650.1 \| Symbols: KJC2 \| KONJAC 2 \| chr2:1621986-1624486 REVERSE LENGTH=406 | 1 |
| AT1G08820.1 \| Symbols: VAP27-2 \| vamp/synaptobrevin-associated protein 27-2 \| chr1:2821810-2824412 REVERSE LENGTH=386 | 1 |
| AT1G16890.2 \| Symbols: UBC13B, AtUBC36, UBC36 \| ubiquitin-conjugating enzyme 36, UBIQUITIN CONJUGATING ENZYME 13B \| chr1:5776550-5778327 REVERSE LENGTH=153 | 1 |
| AT2G32260.1 \| Symbols: ATCCT1, CCT1 \| phosphorylcholine cytidylyltransferase \| chr2:13697645-13700241 FORWARD LENGTH=332 | 1 |
| AT3G53870.1 \| Symbols: no symbol available \| no full name available \| chr3:19951547-19952782 FORWARD LENGTH=249 | 1 |
| AT2G20420.1 \| Symbols: no symbol available \| no full name available \| chr2:8805574-8807858 FORWARD LENGTH=421 | 3 |
| AT5G53460.1 \| Symbols: GLT1 \| NADH-dependent glutamate synthase 1 \| chr5:21700518-21709629 FORWARD LENGTH=2208 | 3 |
| AT2G33210.1 \| Symbols: HSP60, HSP60-2 \| heat shock protein 60-2 \| chr2:14075093-14078568 REVERSE LENGTH=585 | 3 |
| AT5G40770.1 \| Symbols: ATPHB3, PHB3, EER3 \| prohibitin 3 \| chr5:16315589-16316621 REVERSE LENGTH=277 | 3 |
| AT5G42080.1 \| Symbols: DRP1A, ADL1A, DL1, AG68, RSW9, ADL1 \| RADIAL SWELLING 9, DYNAMIN-RELATED PROTEIN 1A, dynamin-like protein \| chr5:16820661-16824536 REVERSE LENGTH=610 | 3 |
| AT2G30970.1 \| Symbols: ASP1 \| aspartate aminotransferase 1 \| chr2:13179012-13181686 FORWARD LENGTH=430 | 3 |
| AT5G41970.1 \| Symbols: no symbol available \| no full name available \| chr5:16791198-16792961 FORWARD LENGTH=373 | 3 |
| AT4G29840.1 \| Symbols: MTO2, TS \| THREONINE SYNTHASE, METHIONINE OVER-ACCUMULATOR 2 \| chr4:14599434-14601014 REVERSE LENGTH=526 | 3 |
| AT1G24180.1 \| Symbols: IAR4 \| IAA-CONJUGATE-RESISTANT 4 \| chr1:8560777-8563382 REVERSE LENGTH=393 | 3 |
| AT1G06220.1 \| Symbols: MEE5, CLO, GFA1 \| MATERNAL EFFECT EMBRYO ARREST 5, CLOTHO, GAMETOPHYTE FACTOR 1 \| chr1:1900524-1904583 FORWARD LENGTH=987 | 3 |
| AT4G00570.1 \| Symbols: NAD-ME2 \| NAD-dependent malic enzyme 2 \| chr4:242817-246522 REVERSE LENGTH=607 | 3 |
| AT4G24830.1 \| Symbols: no symbol available \| no full name available \| chr4:12793085-12795857 REVERSE LENGTH=494 | 3 |
| AT5G65720.1 \| Symbols: NFS1, ATNFS1, NIFS1, ATNIFS1 \| NITROGEN FIXATION S HOMOLOG 1, nitrogen fixation S (NIFS)-like 1, ARABIDOPSIS THALIANA NITROGEN FIXATION S (NIFS)-LIKE 1 \| chr5:26296349-26297710 FORWARD LENGTH=453 | 3 |
| AT4G33680.1 \| Symbols: AGD2 \| ABERRANT GROWTH AND DEATH 2, ARF-GAP domain 2 \| chr4:16171847-16174630 REVERSE LENGTH=461 | 3 |
| AT2G43950.1 \| Symbols: ATOEP37, OEP37 \| chloroplast outer envelope protein 37, ARABIDOPSIS CHLOROPLAST OUTER ENVELOPE PROTEIN 37 \| chr2:18200553-18202644 REVERSE LENGTH=343 | 3 |
| AT3G10050.1 \| Symbols: OMR1 \| L-O-methylthreonine resistant 1 \| chr3:3099164-3101741 REVERSE LENGTH=592 | 3 |
| AT2G43090.1 \| Symbols: IPMI SSU1 \| isopropylmalate isomerase small subunit 1 \| chr2:17918957-17919712 FORWARD LENGTH=251 | 3 |
| AT5G23140.1 \| Symbols: CLPP2, NCLPP7 \| nuclear-encoded CLP protease P7 \| chr5:7783811-7784826 FORWARD LENGTH=241 | 3 |
| AT1G15130.1 \| Symbols: ALIX \| ALG-2 INTERACTING PROTEIN-X \| chr1:5206217-5209848 REVERSE LENGTH=846 | 3 |
| AT1G79340.1 \| Symbols: AtMC4, AtMCP2d, MC4, MCP2d \| metacaspase 2d, metacaspase 4 \| chr1:29842849-29844368 FORWARD LENGTH=418 | 3 |
| AT1G53850.1 \| Symbols: ATPAE1, PAE1 \| 20S proteasome alpha subunit E1, ARABIDOPSIS 20S PROTEASOME ALPHA SUBUNIT E1 \| chr1:20104131-20105792 REVERSE LENGTH=237 | 3 |
| AT3G62030.1 \| Symbols: ROC4, CYP20-3 \| rotamase CYP 4, cyclophilin 20-3 \| chr3:22973708-22975139 FORWARD LENGTH=260 | 3 |
| AT4G04180.1 \| Symbols: no symbol available \| no full name available \| chr4:2020471-2023673 FORWARD LENGTH=600 | 3 |
| AT1G64550.1 \| Symbols: AtGCN20, AtABCF3, ABCF3, SCORD5, GCN20 \| susceptible to coronatine-deficient Pst DC3000 5, general control non-repressible 20, ATP-binding cassette F3 \| chr1:23968850-23973369 FORWARD LENGTH=715 | 3 |
| AT4G00620.1 \| Symbols: EMB3127 \| EMBRYO DEFECTIVE 3127 \| chr4:259265-260788 REVERSE LENGTH=360 | 3 |
| AT1G79500.1 \| Symbols: AtkdsA1, KDO8PS \| 3-Deoxy-D-manno-octulosonate 8-phosphate synthase \| chr1:29903604-29905989 FORWARD LENGTH=290 | 3 |
| AT3G02230.1 \| Symbols: RGP1, ATRGP1 \| ARABIDOPSIS THALIANA REVERSIBLY GLYCOSYLATED POLYPEPTIDE 1, reversibly glycosylated polypeptide 1 \| chr3:415463-417304 FORWARD LENGTH=357 | 3 |
| AT3G49680.1 \| Symbols: BCAT3, ATBCAT-3 \| branched-chain aminotransferase 3 \| chr3:18422768-18425473 FORWARD LENGTH=413 | 3 |
| AT5G17310.2 \| Symbols: AtUGP2, UGP2 \| UDP-GLUCOSE PYROPHOSPHORYLASE 2, UDP-glucose pyrophosphorylase 2 \| chr5:5696955-5700845 REVERSE LENGTH=470 | 3 |
| AT3G52850.1 \| Symbols: GFS1, MTV18, BP-80, ATELP1, BP80-1;1, VSR1;1, ATELP, ATVSR1, VSR1, BP80, BP80B \| VACUOLAR SORTING RECEPTOR 1;1, ARABIDOPSIS THALIANA EPIDERMAL GROWTH FACTOR RECEPTOR-LIKE PROTEIN, modified transport to the vacuole 18, Green fluorescent seed 1, binding protein of 80 kDa 1;1, vacuolar sorting receptor homolog 1 \| chr3:19587999-19591690 FORWARD LENGTH=623 | 3 |
| AT3G27380.1 \| Symbols: SDH2-1 \| succinate dehydrogenase 2-1 \| chr3:10131209-10132673 REVERSE LENGTH=279 | 3 |
| AT1G80560.1 \| Symbols: ATIMD2, IMD2 \| isopropylmalate dehydrogenase 2, ARABIDOPSIS ISOPROPYLMALATE DEHYDROGENASE 2 \| chr1:30287833-30290126 FORWARD LENGTH=405 | 3 |
| AT3G46560.1 \| Symbols: TIM9, emb2474, AtTIM9 \| embryo defective 2474, TRANSLOCASE OF THE INNER MEMBRANE 9 \| chr3:17138632-17139301 FORWARD LENGTH=93 | 3 |
| AT1G32580.1 \| Symbols: MORF5 \| Multiple Organellar RNA editing Factor 5 \| chr1:11784108-11785430 FORWARD LENGTH=229 | 3 |
| AT5G09650.1 \| Symbols: PPa6, AtPPa6 \| pyrophosphorylase 6 \| chr5:2991331-2993117 REVERSE LENGTH=300 | 3 |
| AT5G40370.1 \| Symbols: AtGRXC2, GRXC2, GRX370 \| glutaredoxin C2 \| chr5:16147826-16149052 REVERSE LENGTH=111 | 3 |
| AT1G53000.1 \| Symbols: KDSB, AtCKS, CKS \| CMP-KDO synthetase \| chr1:19745330-19747133 REVERSE LENGTH=290 | 3 |
| AT5G16240.1 \| Symbols: AAD1 \| ACYL-ACYL CARRIER PROTEIN DESATURASE1 \| chr5:5306981-5309639 FORWARD LENGTH=394 | 3 |
| AT5G06140.1 \| Symbols: SNX1, ATSNX1 \| ARABIDOPSIS THALIANA SORTING NEXIN 1, sorting nexin 1 \| chr5:1856212-1858752 REVERSE LENGTH=402 | 3 |
| AT1G60940.1 \| Symbols: SNRK2.10, SNRK2-10, SRK2B \| SNF1-related protein kinase 2.10, SUCROSE NONFERMENTING 1-RELATED PROTEIN KINASE 2-10, SNF1-RELATED KINASE 2B \| chr1:22439398-22441896 REVERSE LENGTH=361 | 3 |
| AT3G62310.1 \| Symbols: no symbol available \| no full name available \| chr3:23057516-23060561 REVERSE LENGTH=726 | 3 |
| AT1G51390.1 \| Symbols: NFU5, ATNFU1 \| NFU domain protein 5 \| chr1:19050427-19051753 FORWARD LENGTH=275 | 3 |
| AT5G18170.1 \| Symbols: GDH1 \| glutamate dehydrogenase 1 \| chr5:6006172-6008248 FORWARD LENGTH=411 | 3 |
| AT5G13120.1 \| Symbols: CYP20-2, Pnsl5, ATCYP20-2 \| cyclophilin 20-2, Photosynthetic NDH subcomplex L 5, ARABIDOPSIS THALIANA CYCLOPHILIN 20-2 \| chr5:4162714-4164720 REVERSE LENGTH=259 | 3 |
| AT5G50960.1 \| Symbols: NBP35, ATNBP35 \| NUCLEOTIDE BINDING PROTEIN 35, nucleotide binding protein 35 \| chr5:20734267-20735824 FORWARD LENGTH=350 | 3 |
| AT3G52390.1 \| Symbols: no symbol available \| no full name available \| chr3:19423105-19425183 REVERSE LENGTH=320 | 3 |
| AT4G26840.1 \| Symbols: SUMO1, ATSUMO1, SUMO 1, SUM1 \| ARABIDOPSIS THALIANA SMALL UBIQUITIN-LIKE MODIFIER 1, small ubiquitin-like modifier 1, SMALL UBIQUITIN-LIKE MODIFIER 1 \| chr4:13497466-13498458 FORWARD LENGTH=100 | 3 |
| AT2G34260.1 \| Symbols: WDR55 \| human WDR55 (WD40 repeat) homolog \| chr2:14465899-14468416 FORWARD LENGTH=353 | 3 |
| AT1G61570.1 \| Symbols: TIM13 \| translocase of the inner mitochondrial membrane 13 \| chr1:22718897-22719473 REVERSE LENGTH=87 | 3 |
| AT1G48860.1 \| Symbols: EPSPS \| 5-enolpyruvylshikimate-3-phosphate synthase \| chr1:18068892-18071331 REVERSE LENGTH=521 | 3 |
| AT3G52190.1 \| Symbols: AtPHF1, PHF1 \| phosphate transporter traffic facilitator1 \| chr3:19354117-19356910 REVERSE LENGTH=398 | 3 |
| AT3G17940.1 \| Symbols: no symbol available \| no full name available \| chr3:6143707-6145244 REVERSE LENGTH=341 | 3 |
| AT1G27310.1 \| Symbols: NTF2A \| nuclear transport factor 2A \| chr1:9484615-9485790 REVERSE LENGTH=122 | 3 |
| AT3G55410.1 \| Symbols: E1-OGDH1 \| \| chr3:20541897-20545728 FORWARD LENGTH=1017 | 4 |
| AT5G09590.1 \| Symbols: MTHSC70-2, HSC70-5 \| mitochondrial HSO70 2, HEAT SHOCK COGNATE \| chr5:2975721-2978508 FORWARD LENGTH=682 | 4 |
| AT3G22330.1 \| Symbols: PMH2, ATRH53 \| putative mitochondrial RNA helicase 2 \| chr3:7892641-7895145 FORWARD LENGTH=616 | 4 |
| AT1G29900.1 \| Symbols: CARB, VEN3 \| carbamoyl phosphate synthetase B, VENOSA 3 \| chr1:10468164-10471976 FORWARD LENGTH=1187 | 4 |
| AT5G50920.1 \| Symbols: DCA1, CLPC, ATHSP93-V, CLPC1, HSP93-V \| HEAT SHOCK PROTEIN 93-V, CLPC homologue 1, DE-REGULATED CAO ACCUMULATION 1 \| chr5:20715710-20719800 REVERSE LENGTH=929 | 4 |
| AT1G20260.1 \| Symbols: AtVAB3, VAB3 \| V-ATPase B subunit 3 \| chr1:7016971-7020290 FORWARD LENGTH=487 | 4 |
| AT1G53750.1 \| Symbols: RPT1A \| regulatory particle triple-A 1A \| chr1:20065921-20068324 REVERSE LENGTH=426 | 4 |
| AT2G45300.1 \| Symbols: no symbol available \| no full name available \| chr2:18677518-18679868 FORWARD LENGTH=520 | 4 |
| AT4G16660.1 \| Symbols: HSP70 \| heat shock protein 70 \| chr4:9377225-9381232 FORWARD LENGTH=867 | 4 |
| AT5G26780.1 \| Symbols: SHM2 \| serine hydroxymethyltransferase 2 \| chr5:9418299-9421725 FORWARD LENGTH=517 | 4 |
| AT4G26970.1 \| Symbols: ACO2 \| aconitase 2 \| chr4:13543077-13548427 FORWARD LENGTH=995 | 4 |
| AT3G13300.1 \| Symbols: VCS \| VARICOSE \| chr3:4304085-4309949 FORWARD LENGTH=1344 | 4 |
| AT2G37690.1 \| Symbols: no symbol available \| no full name available \| chr2:15806111-15810240 FORWARD LENGTH=642 | 4 |
| AT5G14590.1 \| Symbols: no symbol available \| no full name available \| chr5:4703533-4706627 REVERSE LENGTH=485 | 4 |
| AT5G54770.1 \| Symbols: THI1, TZ, THI4 \| THIAZOLE REQUIRING, THIAMINE4 \| chr5:22246634-22247891 FORWARD LENGTH=349 | 4 |
| AT5G58290.1 \| Symbols: RPT3 \| regulatory particle triple-A ATPase 3 \| chr5:23569155-23571116 FORWARD LENGTH=408 | 4 |
| AT5G20080.1 \| Symbols: no symbol available \| no full name available \| chr5:6782708-6786360 FORWARD LENGTH=328 | 4 |
| AT2G39730.1 \| Symbols: RCA \| rubisco activase \| chr2:16570951-16573345 REVERSE LENGTH=474 | 4 |
| AT4G23100.1 \| Symbols: CAD2, GSHA, RAX1, AtGSH1, PAD2, RML1, ATECS1, GSH1 \| PHYTOALEXIN DEFICIENT 2, REGULATOR OF AXILLARY MERISTEMS 1, cinnamyl alcohol dehydrogenase homolog 2, glutamate-cysteine ligase, CADMIUM SENSITIVE 2, proteasome alpha subunit D2, ROOT MERISTEMLESS 1 \| chr4:12103458-12106751 REVERSE LENGTH=522 | 4 |
| AT1G55150.1 \| Symbols: AtRH20, RH20 \| RNA helicase 20 \| chr1:20574634-20577141 FORWARD LENGTH=501 | 4 |
| AT3G29320.1 \| Symbols: PHS1 \| alpha-glucan phosphorylase 1 \| chr3:11252871-11257587 FORWARD LENGTH=962 | 4 |
| AT5G65020.1 \| Symbols: ANNAT2, AtANN2 \| annexin 2 \| chr5:25973915-25975554 FORWARD LENGTH=317 | 4 |
| AT3G54660.1 \| Symbols: EMB2360, MIAO, ATGR2, GR2, GR \| glutathione reductase, GRISEA 2 \| chr3:20230356-20233100 REVERSE LENGTH=565 | 4 |
| AT5G41790.1 \| Symbols: CIP1 \| COP1-interactive protein 1 \| chr5:16727530-16732391 FORWARD LENGTH=1586 | 4 |
| AT5G16760.1 \| Symbols: ITPK1, AtITPK1 \| "inositol 1,3,4-trisphosphate 5/6 kinase 1", "Arabidopsis thaliana inositol 1,3,4-trisphosphate 5/6 kinase 1" \| chr5:5509890-5510849 FORWARD LENGTH=319 | 4 |
| AT1G11860.1 \| Symbols: GLDT \| \| chr1:4001801-4003245 FORWARD LENGTH=408 | 4 |
| AT2G16440.1 \| Symbols: MCM4 \| MINICHROMOSOME MAINTENANCE 4 \| chr2:7126536-7130665 REVERSE LENGTH=847 | 4 |
| AT3G58510.1 \| Symbols: RH11 \| RNA Helicase 11 \| chr3:21640608-21643464 FORWARD LENGTH=612 | 4 |
| AT5G23200.1 \| Symbols: no symbol available \| no full name available \| chr5:7807319-7808988 FORWARD LENGTH=399 | 4 |
| AT1G27090.1 \| Symbols: no symbol available \| no full name available \| chr1:9404041-9406098 REVERSE LENGTH=420 | 4 |
| AT4G01610.1 \| Symbols: AtcathB3 \| \| chr4:694857-696937 FORWARD LENGTH=359 | 4 |
| AT2G36250.1 \| Symbols: FTSZ2-1, ATFTSZ2-1 \| \| chr2:15197661-15199932 REVERSE LENGTH=478 | 4 |
| AT5G55280.1 \| Symbols: CPFTSZ, FTSZ1-1, ATFTSZ1-1, FtsZ1 \| CHLOROPLAST FTSZ, homolog of bacterial cytokinesis Z-ring protein FTSZ 1-1, ARABIDOPSIS THALIANA HOMOLOG OF BACTERIAL CYTOKINESIS Z-RING PROTEIN FTSZ 1-1 \| chr5:22420740-22422527 REVERSE LENGTH=433 | 4 |
| AT5G61510.1 \| Symbols: no symbol available \| no full name available \| chr5:24737084-24738975 REVERSE LENGTH=406 | 4 |
| AT2G42710.1 \| Symbols: no symbol available \| no full name available \| chr2:17782352-17784830 FORWARD LENGTH=415 | 4 |
| AT1G65260.1 \| Symbols: VIPP1, PTAC4, IM30 \| VESICLE-INDUCING PROTEIN IN PLASTIDS 1, plastid transcriptionally active 4 \| chr1:24236329-24240428 FORWARD LENGTH=330 | 4 |
| AT5G26680.1 \| Symbols: SAV6, FEN1 \| Flap Endonuclease I, SHADE AVOIDANCE 6 \| chr5:9311882-9315458 REVERSE LENGTH=453 | 4 |
| AT5G47760.1 \| Symbols: ATPGLP2, ATPK5, PGLP2 \| 2-phosphoglycolate phosphatase 2 \| chr5:19343121-19344979 REVERSE LENGTH=301 | 4 |
| AT5G63620.2 \| Symbols: HER2 \| hexenal response 2 \| chr5:25466380-25468296 REVERSE LENGTH=427 | 4 |
| AT2G45200.1 \| Symbols: GOS12, ATGOS12 \| golgi snare 12 \| chr2:18637689-18639640 REVERSE LENGTH=239 | 4 |
| AT2G33730.1 \| Symbols: SMA1 \| SMALL1 \| chr2:14265679-14267880 REVERSE LENGTH=733 | 4 |
| AT3G01120.1 \| Symbols: AtCYS1, CGS, AtCGS1, CGS1, MTO1 \| CYSTATHIONINE GAMMA-SYNTHASE 1, CYSTATHIONINE GAMMA-SYNTHASE, A. thaliana cystathionine gamma-synthetase 1, METHIONINE OVERACCUMULATION 1 \| chr3:39234-41865 REVERSE LENGTH=563 | 4 |
| AT2G41100.1 \| Symbols: TCH3, CML12, ATCAL4 \| ARABIDOPSIS THALIANA CALMODULIN LIKE 4, calmodulin-like 12, TOUCH 3 \| chr2:17138131-17139406 FORWARD LENGTH=324 | 4 |
| AT1G20920.1 \| Symbols: RCF1 \| regulator of CBF gene expression 1 \| chr1:7285342-7288842 FORWARD LENGTH=1166 | 4 |
| AT3G24430.1 \| Symbols: HCF101 \| HIGH-CHLOROPHYLL-FLUORESCENCE 101 \| chr3:8868731-8872154 REVERSE LENGTH=532 | 4 |
| AT5G19180.1 \| Symbols: ECR1 \| E1 C-terminal related 1 \| chr5:6453375-6455750 FORWARD LENGTH=454 | 4 |
| AT3G62940.2 \| Symbols: OTU5 \| ovarian tumor domain (OTU)-containing DUB (deubiquitilating enzyme) 5 \| chr3:23263106-23264245 REVERSE LENGTH=332 | 4 |
| AT4G08390.1 \| Symbols: SAPX \| stromal ascorbate peroxidase \| chr4:5314999-5317071 FORWARD LENGTH=372 | 4 |
| AT5G23570.1 \| Symbols: ATSGS3, SGS3 \| SUPPRESSOR OF GENE SILENCING 3 \| chr5:7943621-7945874 FORWARD LENGTH=625 | 4 |
| AT5G11340.1 \| Symbols: NAA50 \| N-terminal acetyltransferase 50 \| chr5:3619226-3621068 FORWARD LENGTH=164 | 4 |
| AT2G04350.1 \| Symbols: LACS8 \| long-chain acyl-CoA synthetase 8 \| chr2:1516086-1519178 FORWARD LENGTH=720 | 4 |
| AT5G21150.1 \| Symbols: AGO9 \| ARGONAUTE 9 \| chr5:7193472-7198113 FORWARD LENGTH=896 | 4 |
| AT3G27080.1 \| Symbols: TOM20-3 \| translocase of outer membrane 20 kDa subunit 3 \| chr3:9985212-9986560 REVERSE LENGTH=202 | 4 |
| AT2G29080.1 \| Symbols: AtFTSH3, FTSH3 \| FTSH protease 3 \| chr2:12489911-12492999 REVERSE LENGTH=809 | 4 |
| AT3G27280.1 \| Symbols: PHB4, ATPHB4 \| prohibitin 4 \| chr3:10076904-10078051 FORWARD LENGTH=279 | 4 |
| AT1G02920.1 \| Symbols: ATGSTF8, GSTF7, ATGSTF7, ATGST11, GST11 \| GLUTATHIONE S-TRANSFERASE 11, glutathione S-transferase 7, ARABIDOPSIS GLUTATHIONE S-TRANSFERASE 11 \| chr1:658886-659705 REVERSE LENGTH=209 | 4 |
| AT5G15610.2 \| Symbols: no symbol available \| no full name available \| chr5:5079579-5081929 FORWARD LENGTH=413 | 4 |
| AT5G59160.1 \| Symbols: PPO, TOPP2 \| PROTOPORPHYRINOGEN OXIDASE, type one serine/threonine protein phosphatase 2 \| chr5:23879567-23881102 FORWARD LENGTH=312 | 4 |
| AT3G22660.1 \| Symbols: EBP2 \| \| chr3:8016237-8017118 REVERSE LENGTH=293 | 4 |
| AT5G66420.2 \| Symbols: no symbol available \| no full name available \| chr5:26521893-26524986 REVERSE LENGTH=754 | 4 |
| AT1G05180.1 \| Symbols: AXR1 \| AUXIN RESISTANT 1 \| chr1:1498357-1501775 REVERSE LENGTH=540 | 4 |
| AT5G63030.1 \| Symbols: GRXC1 \| glutaredoxin C1 \| chr5:25286352-25287517 FORWARD LENGTH=125 | 4 |
| AT3G58840.1 \| Symbols: PMD1 \| peroxisomal and mitochondrial division factor 1 \| chr3:21757499-21758455 REVERSE LENGTH=318 | 4 |
| AT5G17530.1 \| Symbols: no symbol available \| no full name available \| chr5:5778168-5781863 FORWARD LENGTH=581 | 4 |
| AT3G63520.1 \| Symbols: ATNCED1, ATCCD1, CCD1, NCED1 \| carotenoid cleavage dioxygenase 1, CAROTENOID CLEAVAGE DIOXYGENASE 1 \| chr3:23452940-23455896 FORWARD LENGTH=538 | 4 |
| AT1G03140.1 \| Symbols: PRP18a \| pre-mRNA processing factor 18a \| chr1:754471-756223 REVERSE LENGTH=420 | 4 |
| AT4G18800.1 \| Symbols: ATRAB11B, ATRABA1D, RABA1d, ATHSGBP \| RAB GTPase homolog A1D \| chr4:10320156-10321339 REVERSE LENGTH=214 | 4 |
| AT5G57950.1 \| Symbols: no symbol available \| no full name available \| chr5:23460765-23462128 FORWARD LENGTH=227 | 4 |
| AT5G51410.1 \| Symbols: LUC7RL \| LETHAL UNLESS CBC 7 RL \| chr5:20881821-20883577 REVERSE LENGTH=334 | 4 |
| AT3G23990.1 \| Symbols: HSP60, HSP60-3B \| heat shock protein 60, HEAT SHOCK PROTEIN 60-3B \| chr3:8669013-8672278 FORWARD LENGTH=577 | 5 |
| AT3G13470.1 \| Symbols: CPNB2, Cpn60beta2 \| chaperonin-60beta2 \| chr3:4389685-4392624 FORWARD LENGTH=596 | 5 |
| AT2G44350.1 \| Symbols: ATCS, CSY4 \| CITRATE SYNTHASE 4 \| chr2:18316673-18320524 FORWARD LENGTH=473 | 5 |
| AT4G32520.1 \| Symbols: AtSHMT3, SHM3 \| serine hydroxymethyltransferase 3, SERINE HYDROXYMETHYLTRANSFERASE 3 \| chr4:15689642-15692334 REVERSE LENGTH=529 | 5 |
| AT3G48560.1 \| Symbols: CSR1, IMR1, TZP5, ALS, AHAS \| ACETOLACTATE SYNTHASE, TRIAZOLOPYRIMIDINE RESISTANT 5, IMIDAZOLE RESISTANT 1, ACETOHYDROXY ACID SYNTHASE, chlorsulfuron/imidazolinone resistant 1 \| chr3:18001530-18003542 REVERSE LENGTH=670 | 5 |
| AT5G35360.1 \| Symbols: CAC2 \| acetyl Co-enzyme a carboxylase biotin carboxylase subunit \| chr5:13584300-13588268 FORWARD LENGTH=537 | 5 |
| AT4G34110.1 \| Symbols: ATPAB2, PAB2, PABP2 \| poly(A) binding protein 2, ARABIDOPSIS POLY(A) BINDING 2, POLY(A) BINDING PROTEIN 2 \| chr4:16336732-16339892 FORWARD LENGTH=629 | 5 |
| AT5G27380.1 \| Symbols: GSH2, AtGSH2, GSHB \| glutathione synthetase 2 \| chr5:9668211-9670912 REVERSE LENGTH=539 | 5 |
| AT1G48900.1 \| Symbols: no symbol available \| no full name available \| chr1:18084972-18087743 REVERSE LENGTH=495 | 5 |
| AT2G17190.1 \| Symbols: DSK2a \| \| chr2:7478272-7481361 REVERSE LENGTH=538 | 5 |
| AT3G46740.1 \| Symbols: 1-Mar, TOC75, TOC75-III \| MODIFIER OF ARG1 1, translocon at the outer envelope membrane of chloroplasts 75-III \| chr3:17216104-17219296 REVERSE LENGTH=818 | 5 |
| AT2G20140.1 \| Symbols: RPT2b \| regulatory particle AAA-ATPase 2b \| chr2:8692736-8694837 FORWARD LENGTH=443 | 5 |
| AT1G79440.1 \| Symbols: ALDH5F1, ENF1, SSADH1, SSADH \| ENLARGED FIL EXPRESSION DOMAIN 1, SUCCINIC SEMIALDEHYDE DEHYDROGENASE, SUCCINIC SEMIALDEHYDE DEHYDROGENASE 1, aldehyde dehydrogenase 5F1 \| chr1:29882525-29887275 REVERSE LENGTH=528 | 5 |
| AT5G16070.1 \| Symbols: CCT6-1 \| Chaperonin containing T-complex polypeptide-1 subunit 6-1 \| chr5:5247549-5251050 REVERSE LENGTH=535 | 5 |
| AT1G14810.1 \| Symbols: no symbol available \| no full name available \| chr1:5102684-5104633 REVERSE LENGTH=375 | 5 |
| AT5G42790.1 \| Symbols: ARS5, PAF1, ATPSM30 \| ARSENIC TOLERANCE 5, proteasome alpha subunit F1 \| chr5:17159270-17160975 REVERSE LENGTH=278 | 5 |
| AT5G14780.1 \| Symbols: FDH, AtFDH1 \| formate dehydrogenase \| chr5:4777043-4779190 FORWARD LENGTH=384 | 5 |
| AT3G09350.1 \| Symbols: Fes1A \| Fes1A \| chr3:2871216-2873109 FORWARD LENGTH=363 | 5 |
| AT1G04410.1 \| Symbols: c-NAD-MDH1 \| cytosolic-NAD-dependent malate dehydrogenase 1 \| chr1:1189418-1191267 REVERSE LENGTH=332 | 5 |
| AT3G53110.1 \| Symbols: LOS4 \| LOW EXPRESSION OF OSMOTICALLY RESPONSIVE GENES 4 \| chr3:19687968-19690423 FORWARD LENGTH=496 | 5 |
| AT3G53580.1 \| Symbols: no symbol available \| no full name available \| chr3:19864784-19866907 FORWARD LENGTH=362 | 5 |
| AT5G61780.1 \| Symbols: Tudor2, AtTudor2, TSN2 \| Arabidopsis thaliana TUDOR-SN protein 2, TUDOR-SN protein 2 \| chr5:24822012-24826641 FORWARD LENGTH=985 | 5 |
| AT1G59900.1 \| Symbols: AT-E1 ALPHA, E1 ALPHA, IAR4L \| pyruvate dehydrogenase complex E1 alpha subunit, IAR4-LIKE \| chr1:22051368-22053660 FORWARD LENGTH=389 | 5 |
| AT3G60820.1 \| Symbols: PBF1 \| \| chr3:22472038-22473809 REVERSE LENGTH=223 | 5 |
| AT4G25340.1 \| Symbols: ATFKBP53, FKBP53 \| FK506 BINDING PROTEIN 53 \| chr4:12959657-12962632 REVERSE LENGTH=477 | 5 |
| AT2G29450.1 \| Symbols: AT103-1A, ATGSTU1, ATGSTU5, GSTU5 \| glutathione S-transferase tau 5, ARABIDOPSIS THALIANA GLUTATHIONE S-TRANSFERASE TAU 1 \| chr2:12624774-12625566 REVERSE LENGTH=224 | 5 |
| AT4G01850.1 \| Symbols: AtSAM2, SAM-2, MAT2, SAM2 \| S-adenosylmethionine synthetase 2, S-ADENOSYLMETHIONINE SYNTHETASE 2 \| chr4:796298-797479 REVERSE LENGTH=393 | 5 |
| AT2G26140.1 \| Symbols: ftsh4, AtFtsH4 \| FTSH protease 4 \| chr2:11131939-11135126 REVERSE LENGTH=717 | 5 |
| AT4G04910.1 \| Symbols: NSF \| N-ethylmaleimide sensitive factor \| chr4:2489696-2495666 REVERSE LENGTH=742 | 5 |
| AT5G37510.1 \| Symbols: EMB1467, CI76 \| embryo defective 1467 \| chr5:14897490-14900352 FORWARD LENGTH=745 | 5 |
| AT1G44900.1 \| Symbols: ATMCM2, MCM2 \| MINICHROMOSOME MAINTENANCE 2 \| chr1:16970291-16974457 FORWARD LENGTH=936 | 5 |
| AT2G32120.1 \| Symbols: HSP70T-2 \| heat-shock protein 70T-2 \| chr2:13651720-13653411 REVERSE LENGTH=563 | 5 |
| AT2G22360.1 \| Symbols: DJA6 \| DNA J protein A6 \| chr2:9498162-9500459 FORWARD LENGTH=442 | 5 |
| AT2G32520.2 \| Symbols: no symbol available \| no full name available \| chr2:13805823-13807676 REVERSE LENGTH=317 | 5 |
| AT3G54640.1 \| Symbols: TRP3, TSA1 \| TRYPTOPHAN-REQUIRING 3, tryptophan synthase alpha chain \| chr3:20223331-20225303 REVERSE LENGTH=312 | 5 |
| AT3G52960.1 \| Symbols: PrxIIE \| peroxiredoxin-II-E \| chr3:19639699-19640403 FORWARD LENGTH=234 | 5 |
| AT2G22475.1 \| Symbols: GEM \| GL2-EXPRESSION MODULATOR \| chr2:9541523-9544778 FORWARD LENGTH=299 | 5 |
| AT3G13160.1 \| Symbols: RPPR3b \| Ribosomal Pentatricopeptide Repeat Protein 3b \| chr3:4229994-4231178 REVERSE LENGTH=394 | 5 |
| AT4G11120.1 \| Symbols: no symbol available \| no full name available \| chr4:6778066-6779934 FORWARD LENGTH=395 | 5 |
| AT2G17200.1 \| Symbols: DSK2, DSK2b \| \| chr2:7482133-7485090 REVERSE LENGTH=551 | 5 |
| AT5G50810.1 \| Symbols: TIM8 \| translocase inner membrane subunit 8 \| chr5:20675875-20676505 REVERSE LENGTH=77 | 5 |
| AT3G61140.1 \| Symbols: FUS6, ATFUS6, EMB78, ATSK31, AtCSN1, COP11, CSN1, SK31 \| ARABIDOPSIS THALIANA FUSCA 6, EMBRYO DEFECTIVE 78, CONSTITUTIVE PHOTOMORPHOGENIC 11, COP9 SIGNALOSOME SUBUNIT 1, FUSCA 6, SHAGGY-LIKE KINASE 31 \| chr3:22626335-22628895 FORWARD LENGTH=441 | 5 |
| AT5G48300.1 \| Symbols: ADG1, APS1 \| ADP-GLUCOSE PYROPHOSPHORYLASE SMALL SUBUNIT 1, ADP glucose pyrophosphorylase 1 \| chr5:19570326-19572557 FORWARD LENGTH=520 | 5 |
| AT3G28940.1 \| Symbols: no symbol available \| no full name available \| chr3:10968324-10969311 REVERSE LENGTH=169 | 5 |
| AT5G62270.2 \| Symbols: GCD1 \| GAMETE CELL DEFECTIVE 1 \| chr5:25012227-25014486 FORWARD LENGTH=420 | 5 |
| AT2G23930.1 \| Symbols: SNRNP-G \| probable small nuclear ribonucleoprotein G \| chr2:10182353-10183026 FORWARD LENGTH=80 | 5 |
| AT3G10670.1 \| Symbols: ABCI6, ATNAP7, NAP7 \| non-intrinsic ABC protein 7, ATP-binding cassette I6 \| chr3:3335325-3337304 REVERSE LENGTH=338 | 5 |
| AT4G08900.1 \| Symbols: ARGAH1 \| arginine amidohydrolase 1 \| chr4:5703499-5705180 FORWARD LENGTH=342 | 5 |
| AT5G58440.1 \| Symbols: SNX2a \| sorting nexin 2A \| chr5:23624154-23626676 REVERSE LENGTH=587 | 5 |
| AT5G08540.1 \| Symbols: no symbol available \| no full name available \| chr5:2763914-2765431 FORWARD LENGTH=346 | 5 |
| AT4G32760.1 \| Symbols: TOL9 \| TOM1-LIKE 9 \| chr4:15799376-15803832 FORWARD LENGTH=675 | 5 |
| AT1G72810.1 \| Symbols: TSY \| THREONINE SYNTHASE 2 \| chr1:27398760-27400393 REVERSE LENGTH=516 | 5 |
| AT3G15020.1 \| Symbols: mMDH2 \| mitochondrial malate dehydrogenase 2 \| chr3:5056139-5057941 FORWARD LENGTH=341 | 5 |
| AT4G29510.1 \| Symbols: ATPRMT11, PRMT11, PRMT1B, ATPRMT1B \| ARABIDOPSIS THALIANA PROTEIN ARGININE METHYLTRANSFERASE 1B, ARABIDOPSIS THALIANA ARGININE METHYLTRANSFERASE 11, PROTEIN ARGININE METHYLTRANSFERASE 1B, arginine methyltransferase 11 \| chr4:14491739-14493752 FORWARD LENGTH=390 | 5 |
| AT5G37475.1 \| Symbols: no symbol available \| no full name available \| chr5:14866328-14867749 REVERSE LENGTH=225 | 5 |
| AT2G39020.1 \| Symbols: GNAT8, NATA2 \| GCN5&#8208;related N&#8208;acetyltransferase 8 \| chr2:16295392-16296102 FORWARD LENGTH=236 | 5 |
| AT1G72680.1 \| Symbols: ATCAD1, CAD1 \| CINNAMYL ALCOHOL DEHYDROGENASE 1, cinnamyl-alcohol dehydrogenase \| chr1:27359346-27360876 REVERSE LENGTH=355 | 5 |
| AT3G60660.1 \| Symbols: no symbol available \| no full name available \| chr3:22421481-22423167 FORWARD LENGTH=272 | 5 |
| AT4G14110.1 \| Symbols: EMB143, COP9, CSN8, FUS7 \| CONSTITUTIVE PHOTOMORPHOGENIC 9, FUSCA 7, EMBRYO DEFECTIVE 143, COP9 SIGNALOSOME SUBUNIT 8 \| chr4:8133049-8134867 REVERSE LENGTH=197 | 5 |
| AT5G19680.1 \| Symbols: PP1R3 \| PROTEIN PHOSPATASE 1 REGULATORY SUBUNIT 3 \| chr5:6649663-6651564 FORWARD LENGTH=328 | 5 |

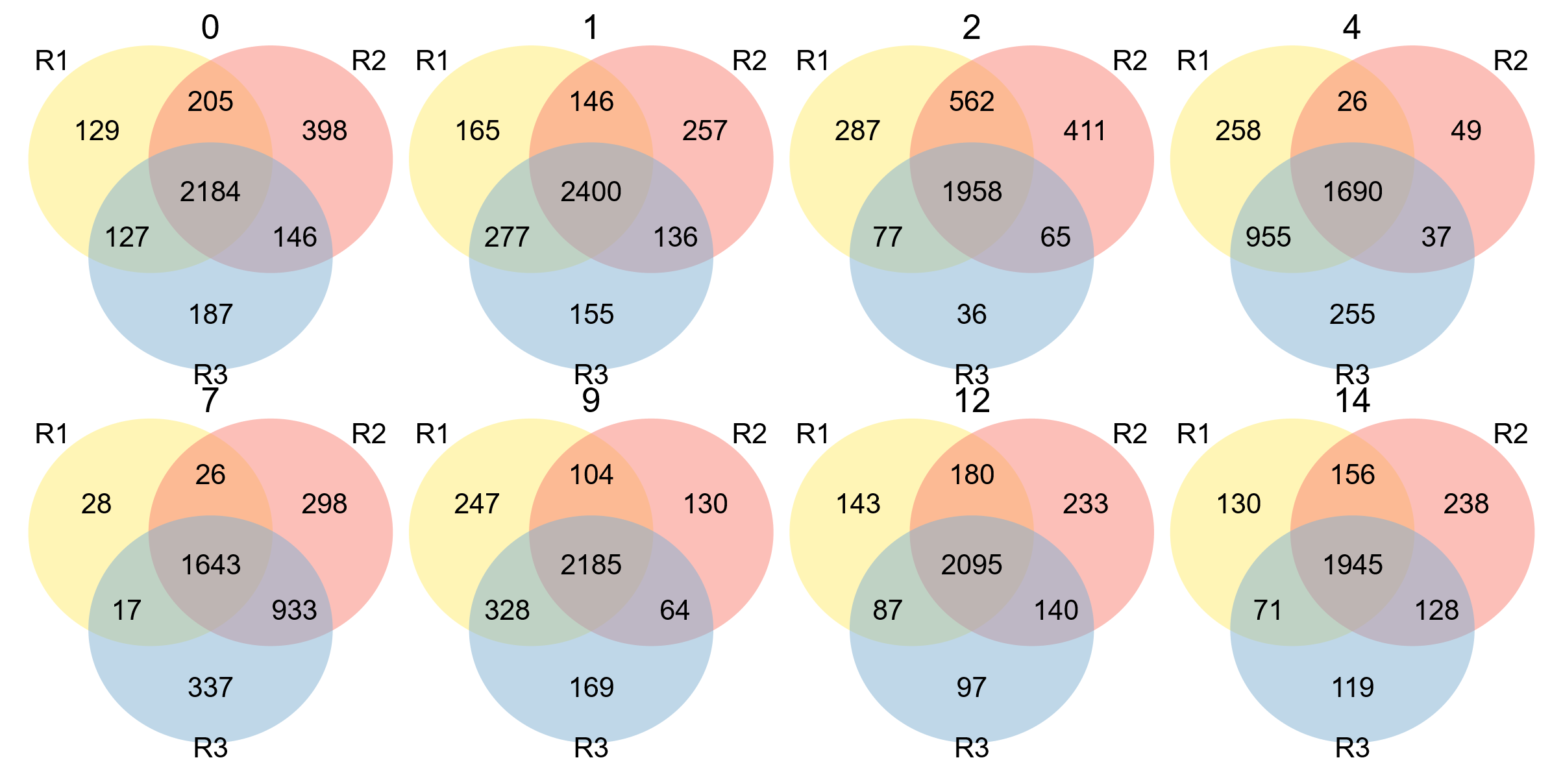

Supplemental Figure S1. Number of unique identified protein in each replicate and time point in samples analysed in this study. After analysis of the peptide mass spectra using FragPipe, data analysis in R revealed that the proteins identified and quantified at each time point were consistent amongst replicates for that timepoint. Approximately 1945–2400 AGIs are shared between all three replicates. The number above each Venn diagram indicates how many days into the chase the samples were taken.

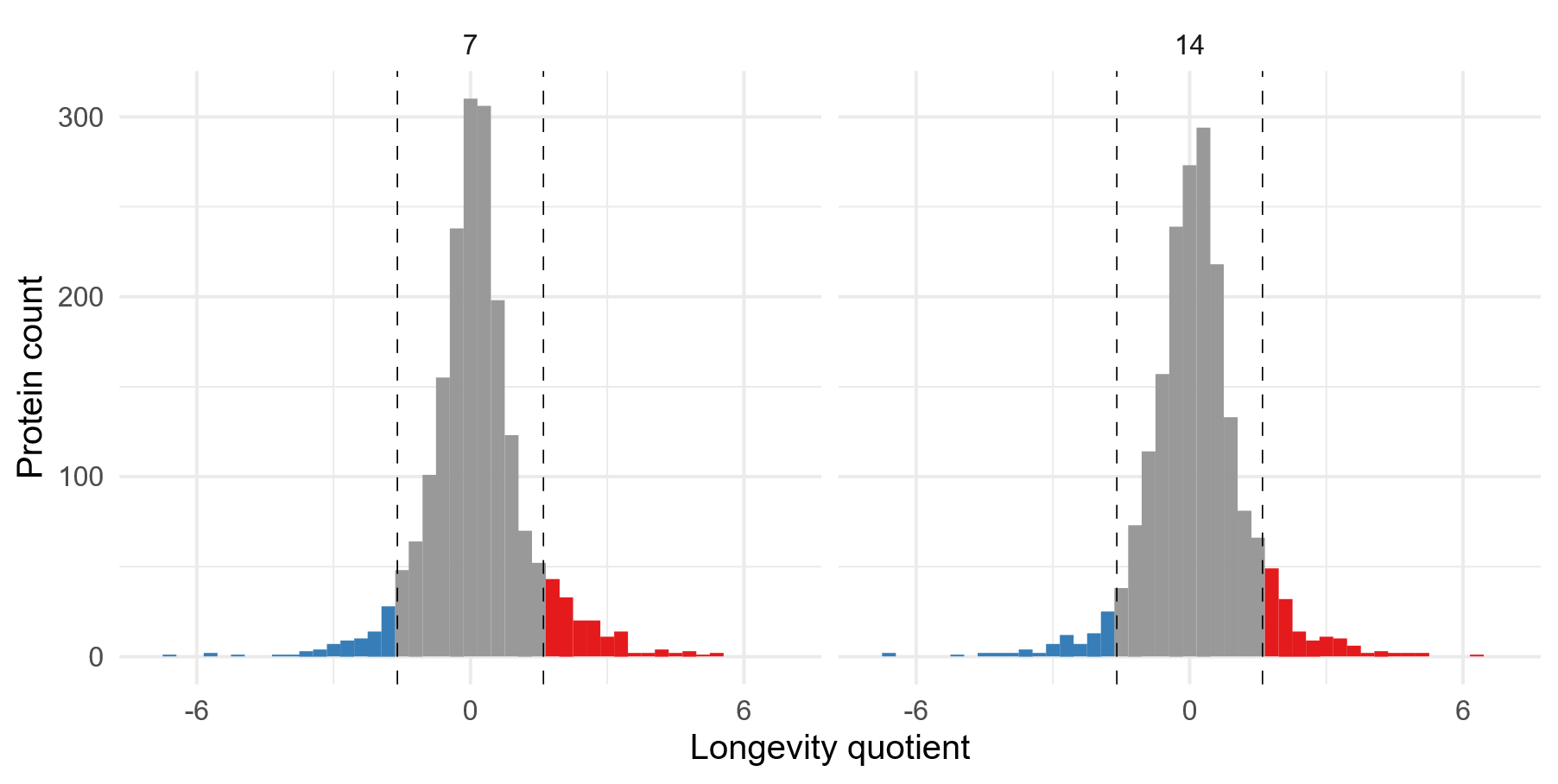

Supplemental Figure S2. Time faceted longevity quotient histograms for proteins in control data (treated with HPG but not enriched).

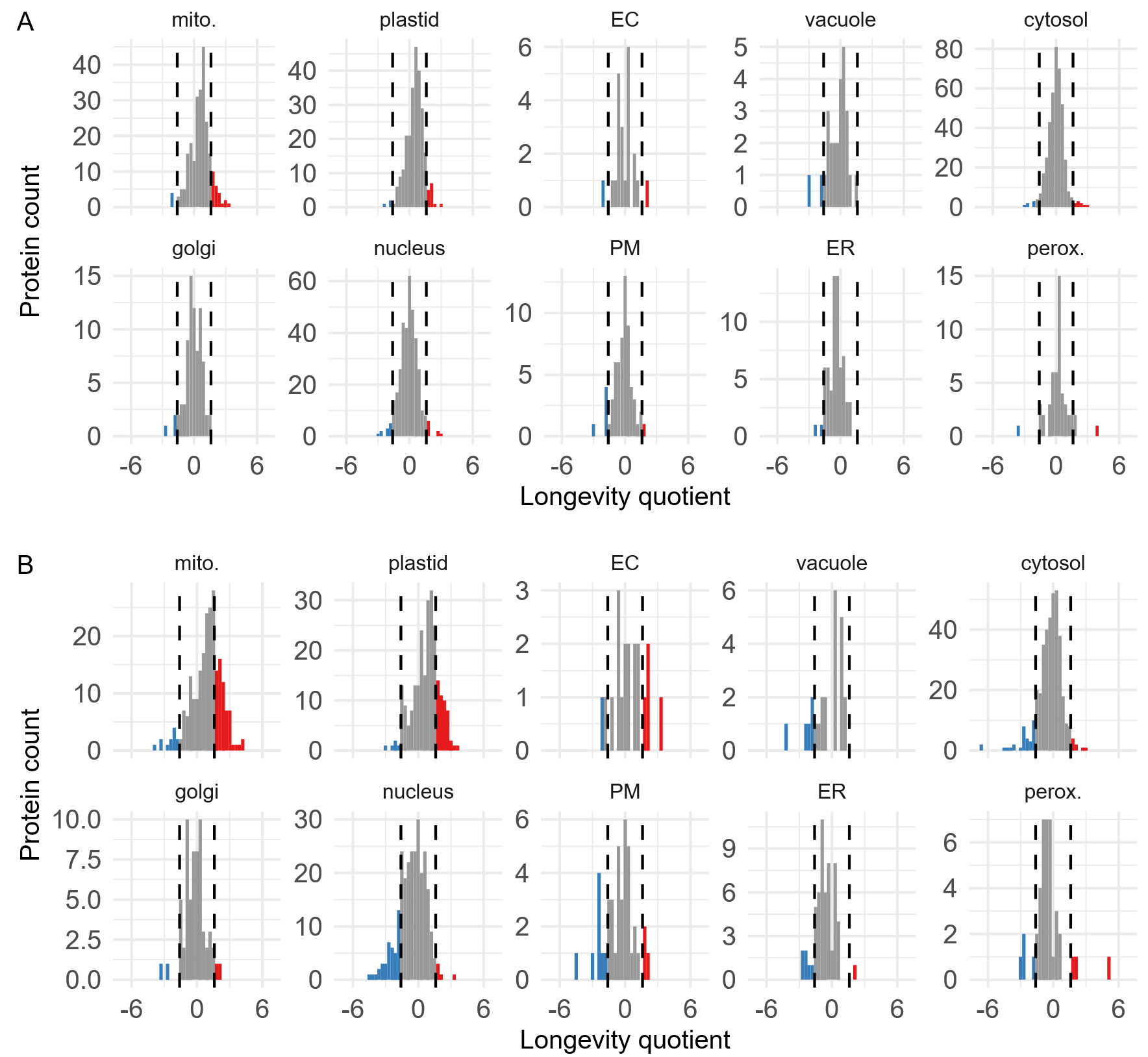

Supplemental Figure S3. (A) Mean longevity quotient histograms, across all timepoints, faceted by SUBAcon location. (B) longevity quotients at 14-day samples, faceted by SUBAcon location.

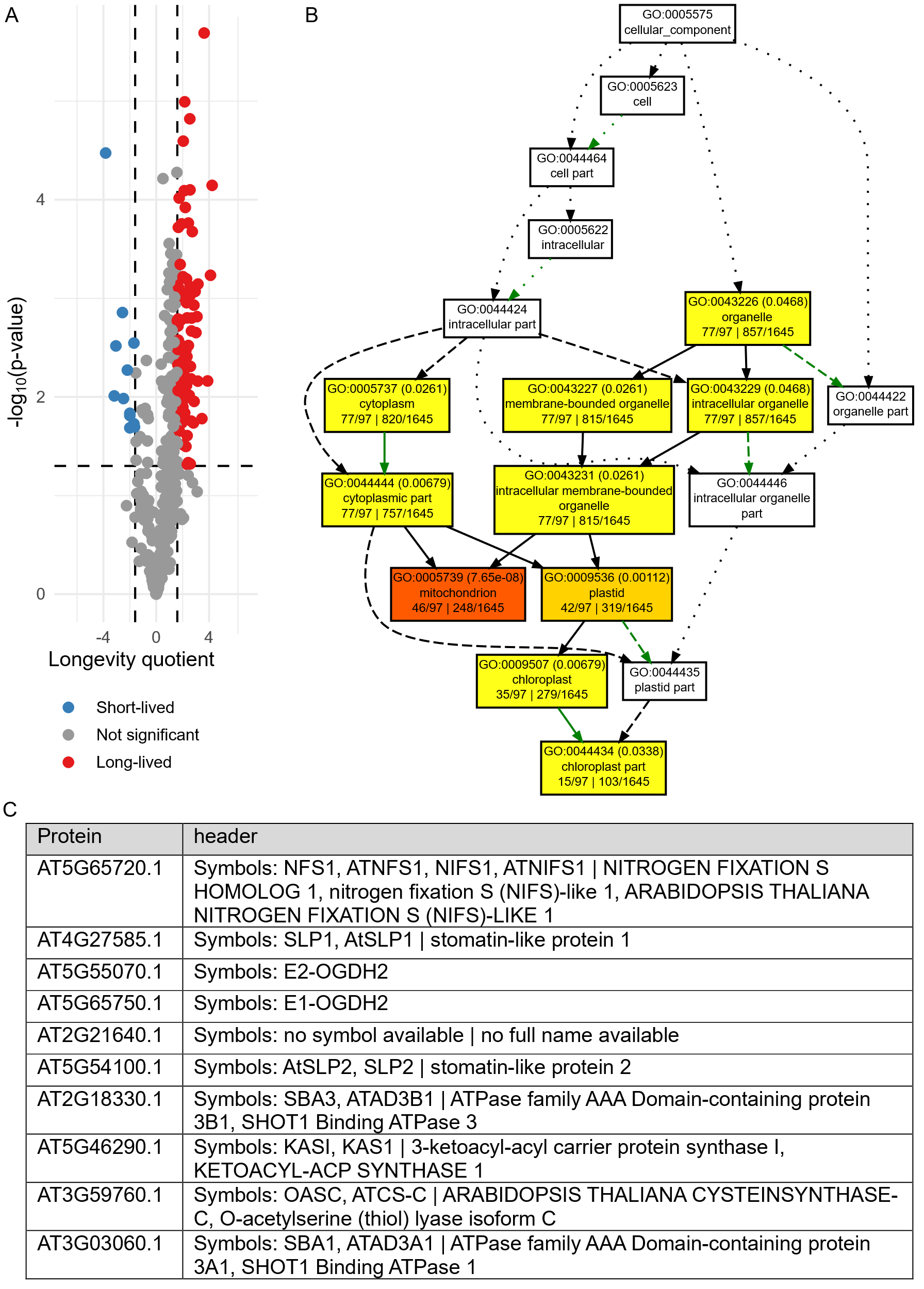

Supplemental Figure S4. (A) Volcano plot for mitochondrial and plastid located proteins. Dashed lines are at 1.3 (-log(0.05), y-axis) and -1.6 and 1.6 (x-axis). Proteins with p-values and longevity quotients outside these parameters (inclusive) were considered short or long-lived as shown in blue and red. (B) GO term enrichment for long-lived proteins (defined as having a longevity quotient >= 1.6 after two weeks). Significant enrichment can be seen in mitochondrial and plastid proteins. (C) Top 10 most significant DAPs in mitochondria or plastids. All except AT2G21640.1 were significantly longer lived.

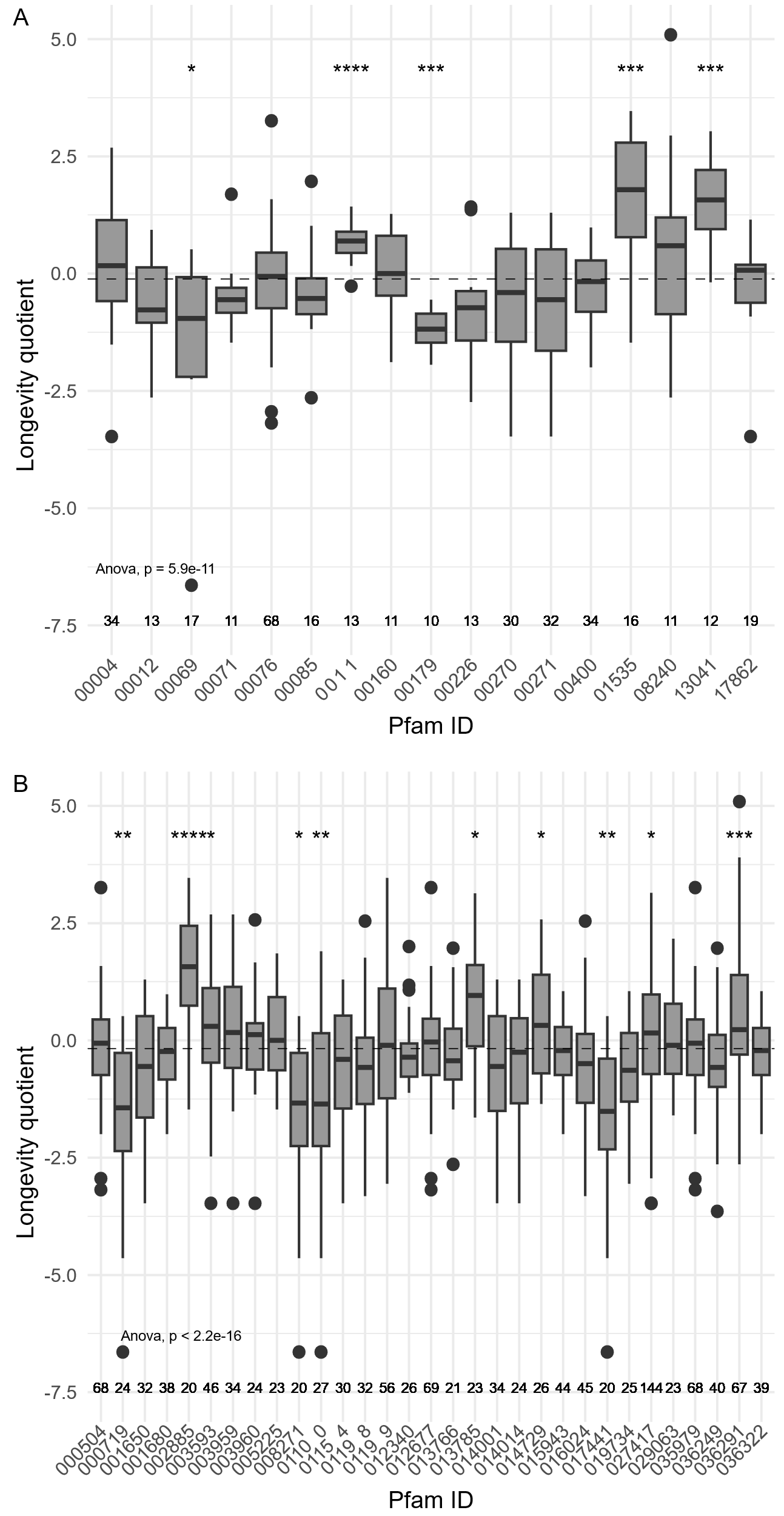

Supplemental Figure S5. (A) Pfam and (B) Interpro boxplot of longevity quotients for proteins with different domains. ‘PF’ and ‘IPR’ portions of the domain numbers removed for breivity. Statistically different groups (t-test) compared to mean (dashed lines) are denoted by asterisks. *p ≤ 0.05, **p ≤ 0.01, ***p ≤ 0.001, ****p ≤ 0.0001.

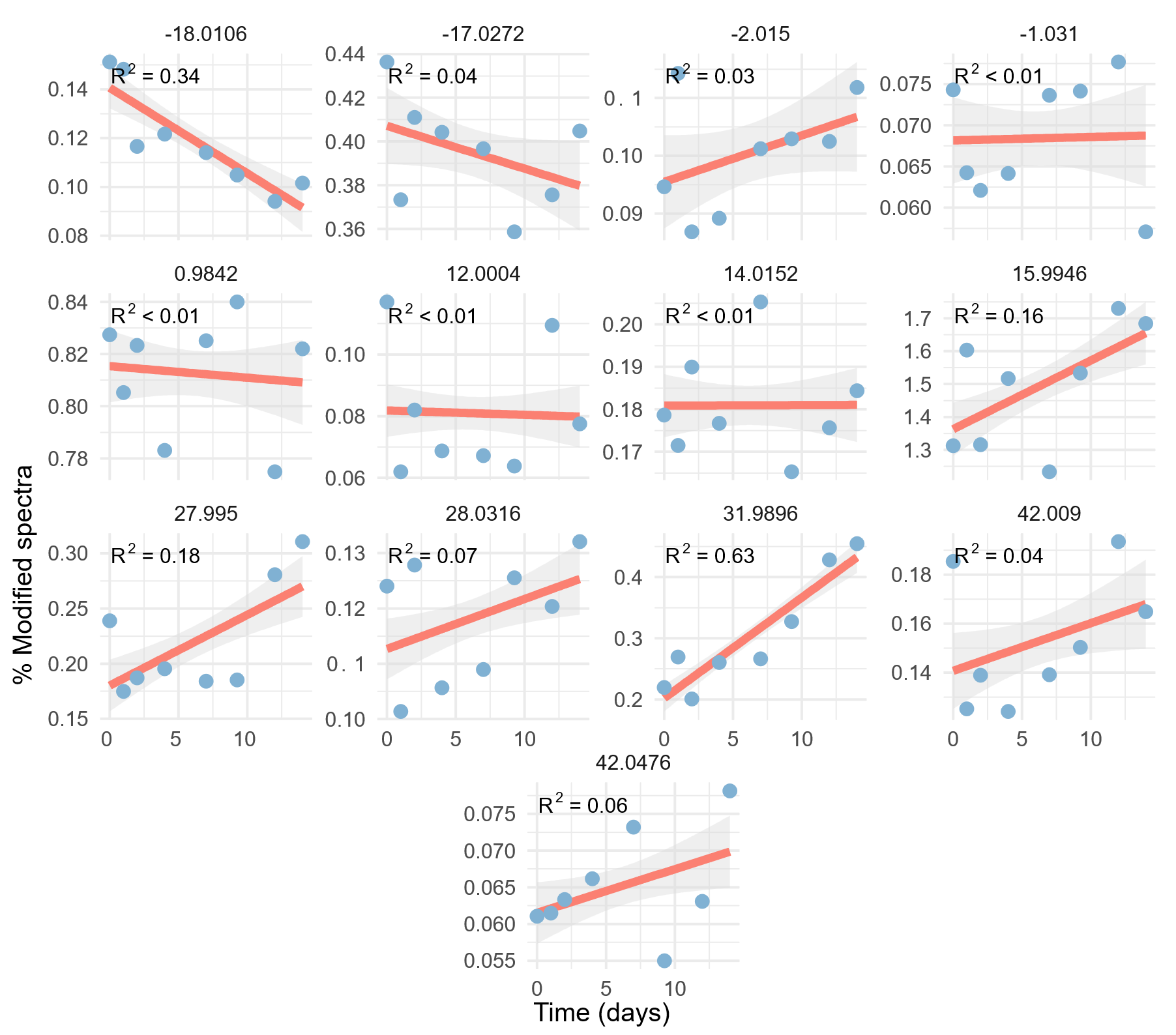

Supplemental Figure S6. Percentage of modified spectral counts vs. time faceted by observed mass shifts in open searches of proteomics data. Chemical identity of mass shifts are shown in Supplemental Table S1. Dots represents means (n = 3); shaded area represents ± 1 standard error.

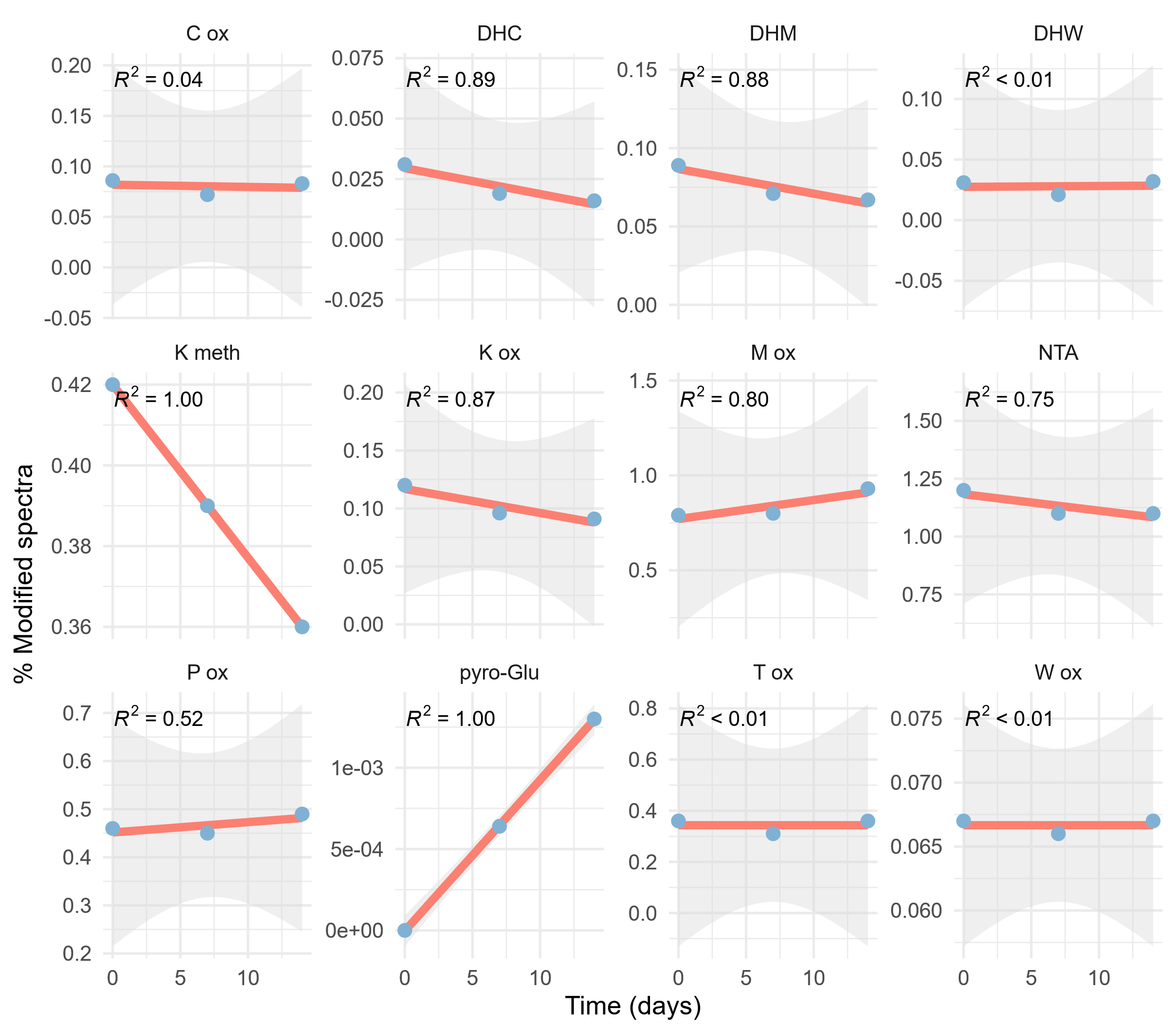

Supplemental Figure S7. Correlations of various PTMs in Arabidopsis cell culture over time, during a two-week chase following a 24-hour pulse with HPG. These data are from samples that have not undergone click-chemistry enrichment. The pyro-Glu from E graph is based on modifications of 3 PSMs out of 4565 PSMs over the three time points, making this general trend unreliable. The perfect fit of the K methylation data also make this correlation unreliable.

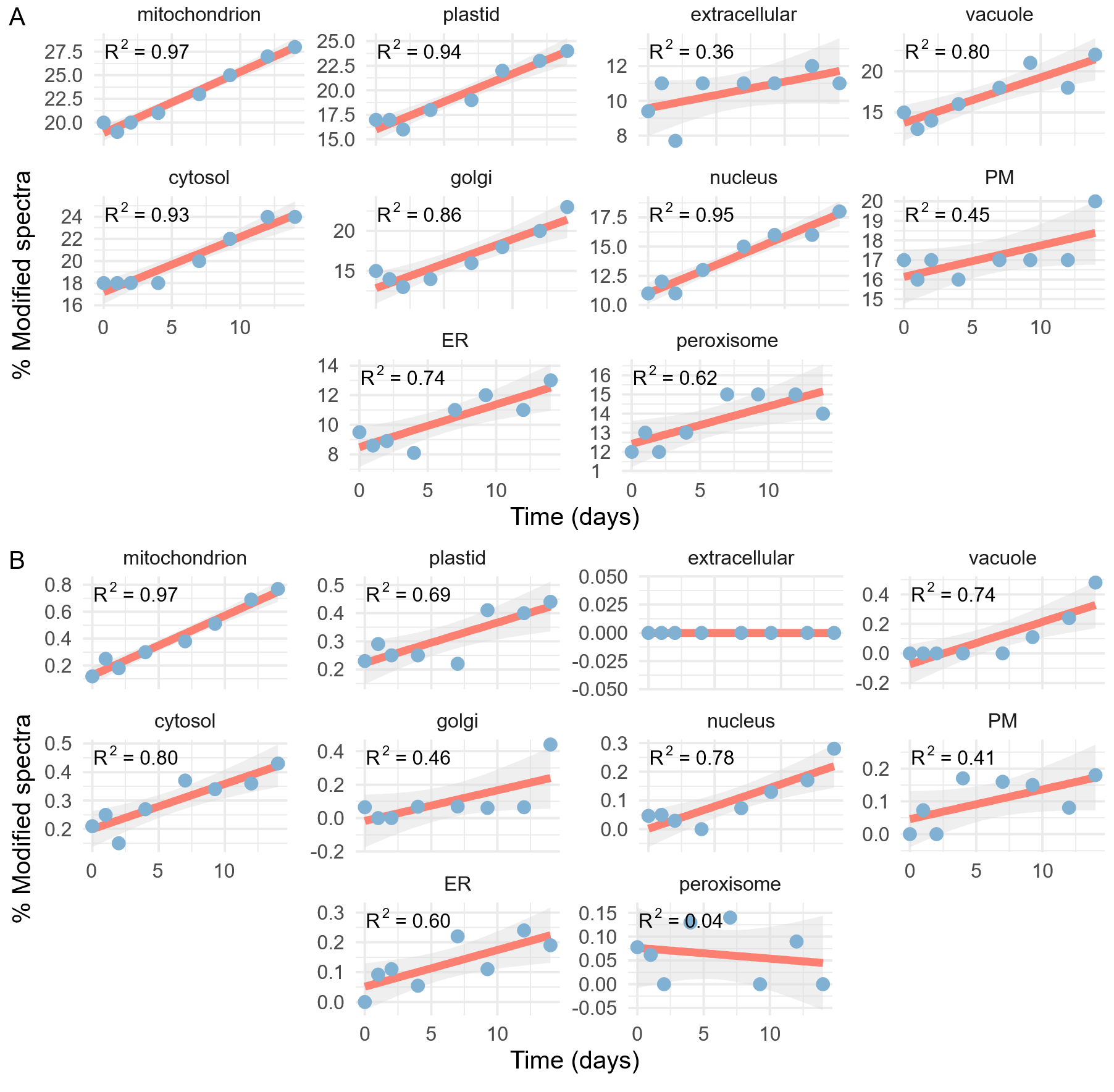

Supplemental Figure S8. Correlation of (A) Met oxidation and (B) DHM PTM modification frequencies in different subcellular compartments.

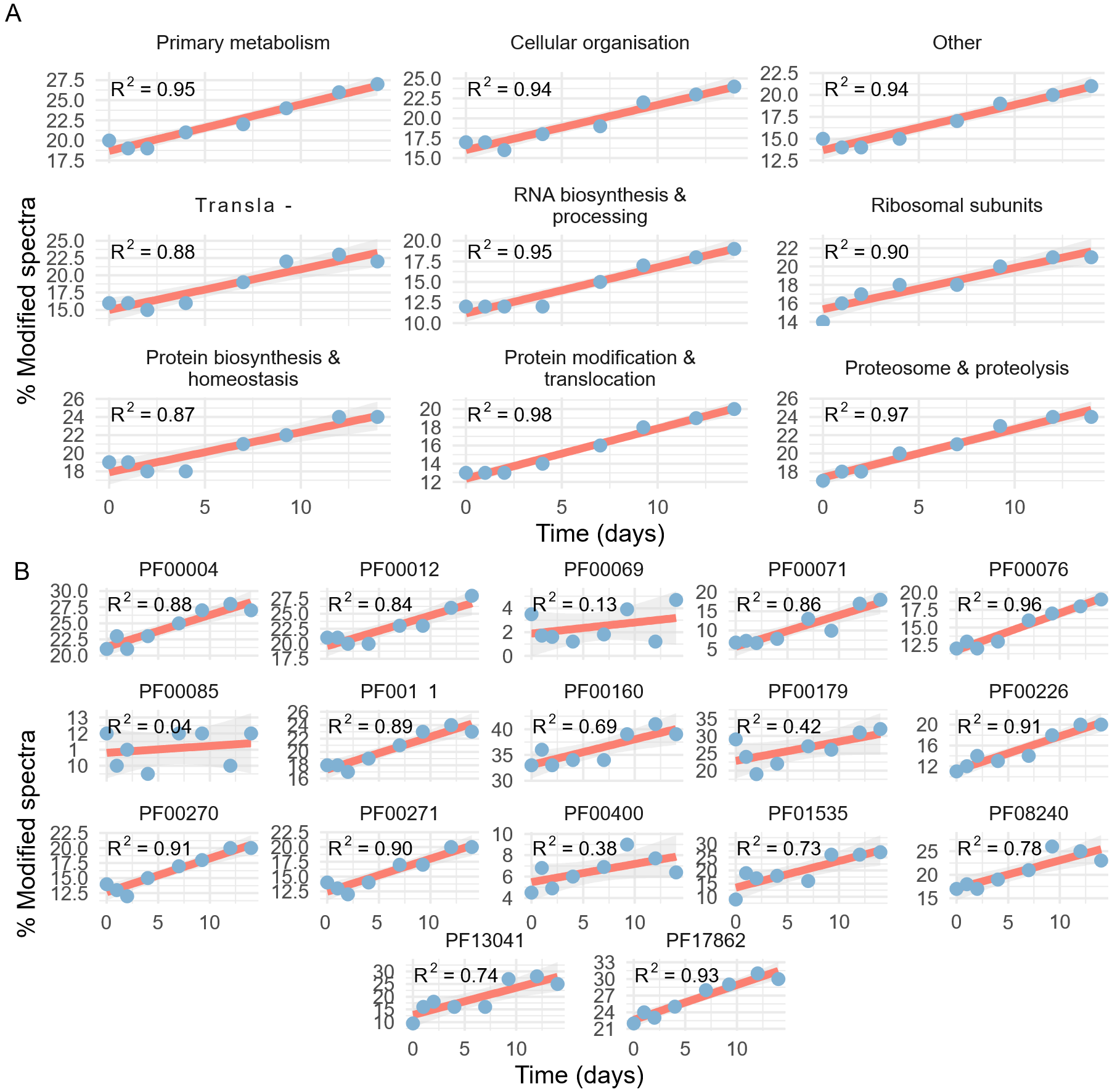

Supplemental Figure S9. Correlation of Met oxidation PTM frequencies with functional categories (A) and Pfam domains (B).

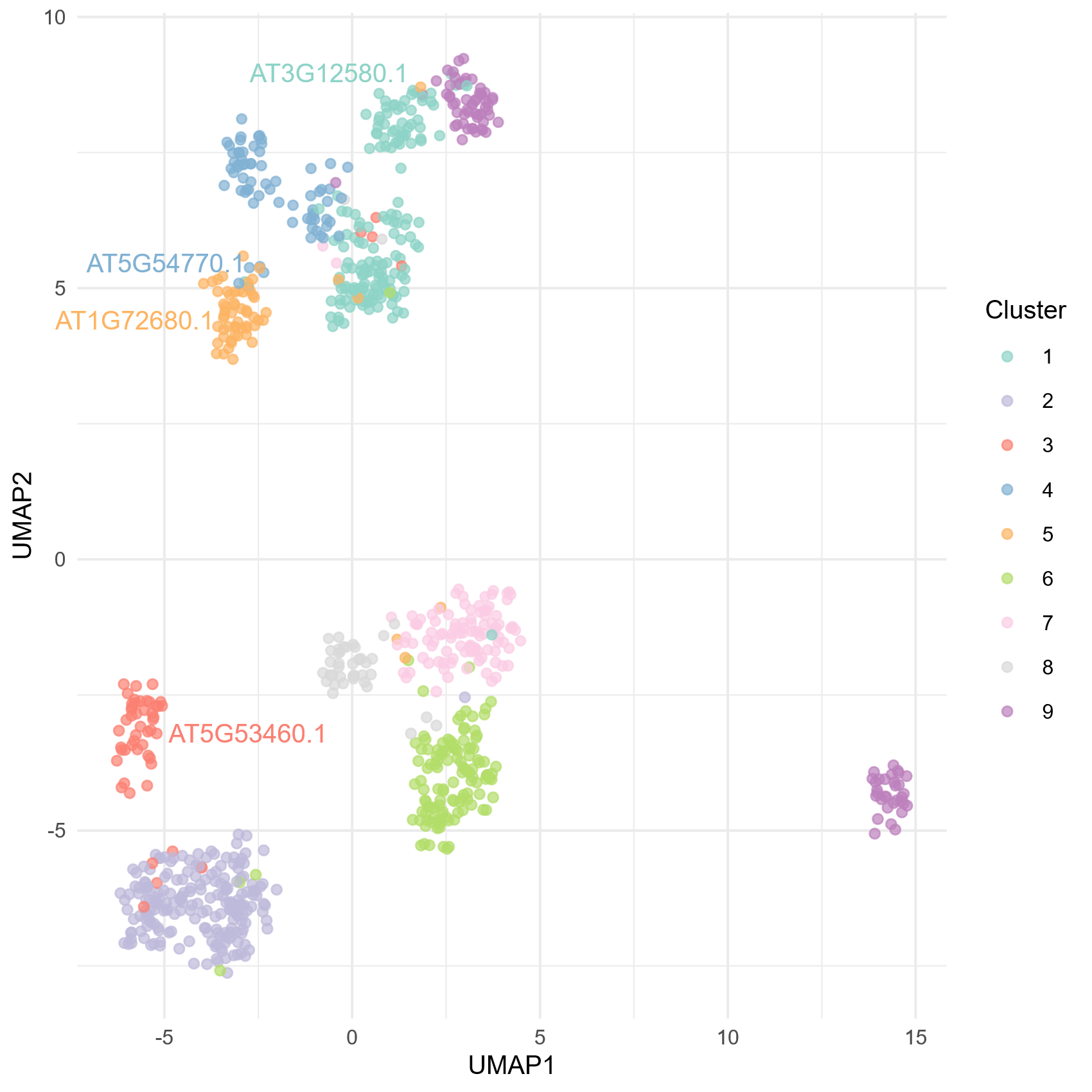

Supplemental Figure S10. K-means clustering of protein groups based on the slope and R^2^ values of regression lines for the relative abundance of 12 PTMs over time. The four exemplar proteins from Figure 5 are labelled in the same colour as the cluster to which they belong and these four clusters are listed in Supplemental Table S8. Cluster membership for all proteins included in the PTM analysis is shown in Supplemental Table S1.

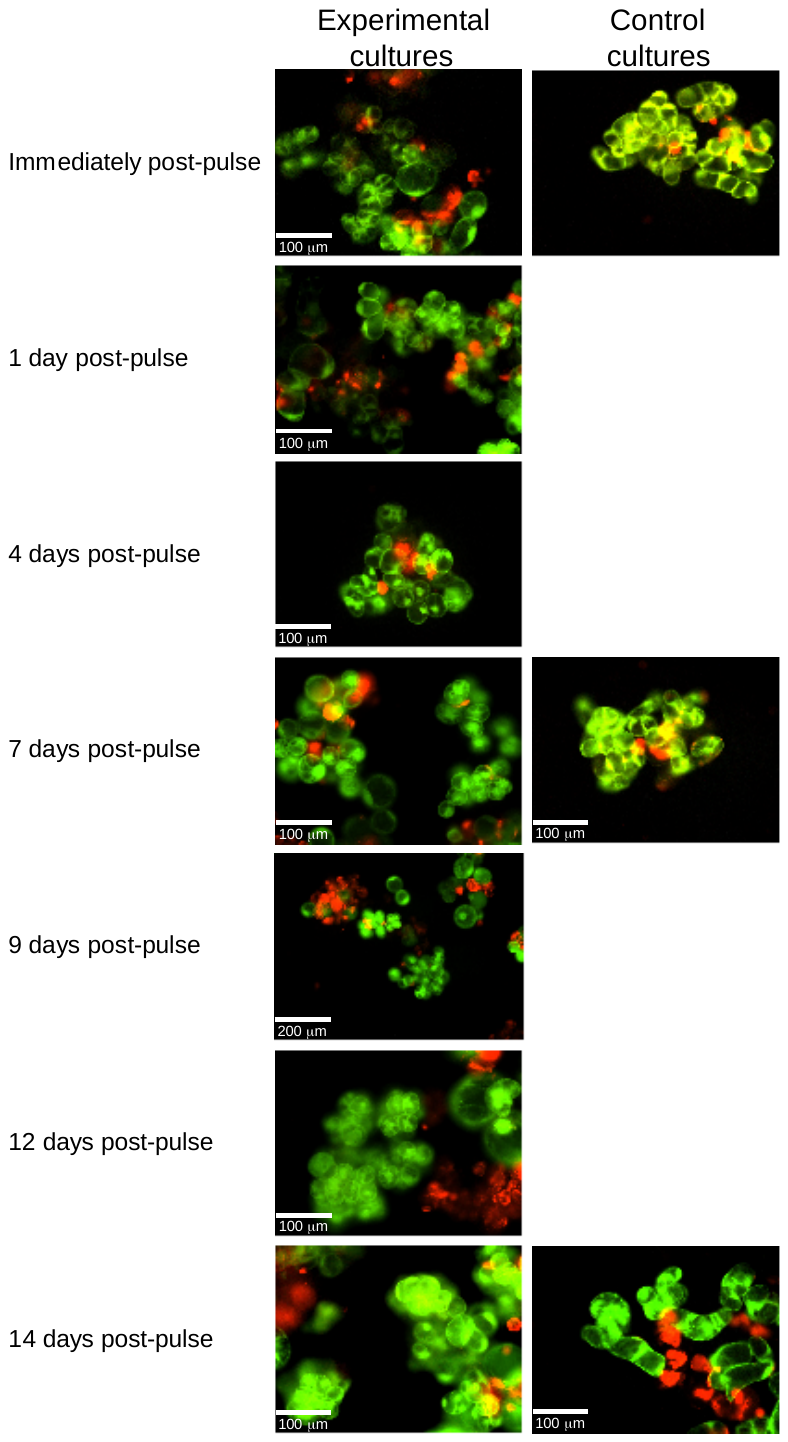

Figure S11. Propidium iodide and fluorescein diacetate viability staining.

Supplemental Figure S12. Cell culture densities after being subject to a 24 h homopropargylglycine pulse followed by a two-week chase. The large drop from 0 days (i.e., at the beginning of the chase) to 1 day is due to loss of cells and dilution of the remaining cells during the washing processes required when switching the cells to HPG-free media.

Density (mg (FW)/mL) normalised to -1 d density

| Time post beginning of chase (days) | R1 | R2 | R3 | R4 | R5 |
| --- | --- | --- | --- | --- | --- |
| -1 | 1 | 1 | 1 | 1 | 1 |
| -0.25 | 1.14387 | 1.0703 | 0.972716 | 1.12155 | 1.01644 |
| 0 | 1.16399 | 1.08333 | 1.22688 | 1.14548 | 1.26791 |
| 1 | 0.554765 | 0.485425 | 0.643106 | 0.510838 | 0.339564 |
| 2 | 0.549075 | 0.606824 | 0.690899 | 0.639847 | 0.381274 |
| 4 | 0.528754 | 0.59928 | 0.682476 | 0.593896 | 0.369505 |
| 7 | 0.714895 | 0.795096 | 0.943051 | 0.719265 | 0.59415 |
| 9.25 | 0.692949 | 0.716907 | 0.819813 | 0.625802 | 0.70855 |
| 12 | 1.08311 | 1.09019 | 1.21333 | 0.941391 | 1.37262 |
| 14 | 2.02906 | 1.66474 | 2.18824 | 1.89808 | 2.25294 |
